# Genomic phylogeography of the White-crowned Manakin *Pseudopipra pipra* (Aves: Pipridae) illuminates a continental-scale radiation out of the Andes

**DOI:** 10.1101/713081

**Authors:** Jacob S. Berv, Leonardo Campagna, Teresa J. Feo, Ivandy Castro-Astor, Camila C. Ribas, Richard O. Prum, Irby J. Lovette

**Affiliations:** Fuller Evolutionary Biology Program, Cornell Lab of Ornithology, 159 Sapsucker Woods Road, Ithaca, NY 14850, USA; Department of Ecology and Evolutionary Biology, Cornell University, 215 Tower Road, Ithaca, NY 14853, USA; Department of Ecology and Evolutionary Biology, and University of Michigan Museum of Paleontology, 1105 North University Avenue, Biological Sciences Building, Ann Arbor, MI 48109-1085, USA; Department of Vertebrate Zoology, MRC-116, National Museum of Natural History, Smithsonian Institution, Washington, DC 20013, USA; Department of Biology, City College of New York and CUNY Graduate Center, City University of New York, New York, NY 10031, USA; Coordenacão de Biodiversidade, Instituto Nacional de Pesquisas da Amazônia, Manaus, AM, Brazil; Department of Ecology and Evolutionary Biology, and Peabody Museum of Natural History, Yale University, New Haven, Connecticut, 06520, USA

**Author notes:** Correspondence to Jacob S. Berv.

**Keywords:** Neotropics, ddRAD, genomics, phylogeography, manakin, suboscine, speciation, introgression, admixture, reticulation, multi-species coalescent, coancestry, isolation by distance, vocalization

## Abstract

The complex landscape history of the Neotropics has generated opportunities for population isolation and diversification that place this region among the most species-rich in the world. Detailed phylogeographic studies are required to uncover the biogeographic histories of Neotropical taxa, to identify evolutionary correlates of diversity, and to reveal patterns of genetic connectivity, disjunction, and potential differentiation among lineages from different areas of endemism. The White-crowned Manakin (*Pseudopipra pipra*) is a small suboscine passerine bird that is broadly distributed through the subtropical rainforests of Central America, the lower montane cloud forests of the Andes from Colombia to central Peru, the lowlands of Amazonia and the Guianas, and the Atlantic forest of southeast Brazil. *Pseudopipra* is currently recognized as a single, polytypic biological species. We studied the effect of the Neotropical landscape on genetic and phenotypic differentiation within this species using genomic data derived from double digest restriction site associated DNA sequencing (ddRAD), and mitochondrial DNA. Most of the genetic breakpoints we identify among populations coincide with physical barriers to gene flow previously associated with avian areas of endemism. The phylogenetic relationships among these populations imply a novel pattern of Andean origination for this group, with subsequent diversification into the Amazonian lowlands. Our analysis of genomic admixture and gene flow reveals a complex history of introgression between some western Amazonian populations. These reticulate processes confound our application of standard concatenated and coalescent phylogenetic methods and raise the question of whether a lineage in the western Napo area of endemism should be considered a hybrid species. Lastly, analysis of variation in vocal and plumage phenotypes in the context of our phylogeny supports the hypothesis that *Pseudopipra* is a species-complex composed of at least 8, and perhaps up to 17 distinct species which have arisen in the last ∼2.5 Ma.

## 1. Introduction

Many kinds of physical and ecological barriers have been proposed to drive population diversification and speciation in the Neotropics (Cracraft and Prum, 1988; Haffer, 1969, 2008; Smith et al., 2014; Wallace, 1854). Such barriers partition biodiversity into areas of endemism (Cracraft, 1985; Cracraft and Prum, 1988; Crother and Murray, 2011; Da Silva et al., 2005; Linder, 2001; Noguera-urbano, 2016) by acting as impediments to gene flow for dispersal-limited organisms (Brumfield, 2012; Cheviron et al., 2005; Fernandes et al., 2015; Moore et al., 2008; Ribas et al., 2012). Three of the most prominent features implicated in structuring the biodiversity of Neotropical forest birds (Figure 1) include the Andes Mountains and other montane regions (Figure 1, grey relief); the Chaco, Cerrado, and Caatinga biomes, which collectively form a “dry diagonal” of open habitat separating the Amazon forest from the Atlantic Forest; and the large rivers of the complex Amazonian drainage system (Brumfield, 2012; Harvey and Brumfield, 2015; Naka and Brumfield, 2018; Smith et al., 2014).

**Figure 1.**
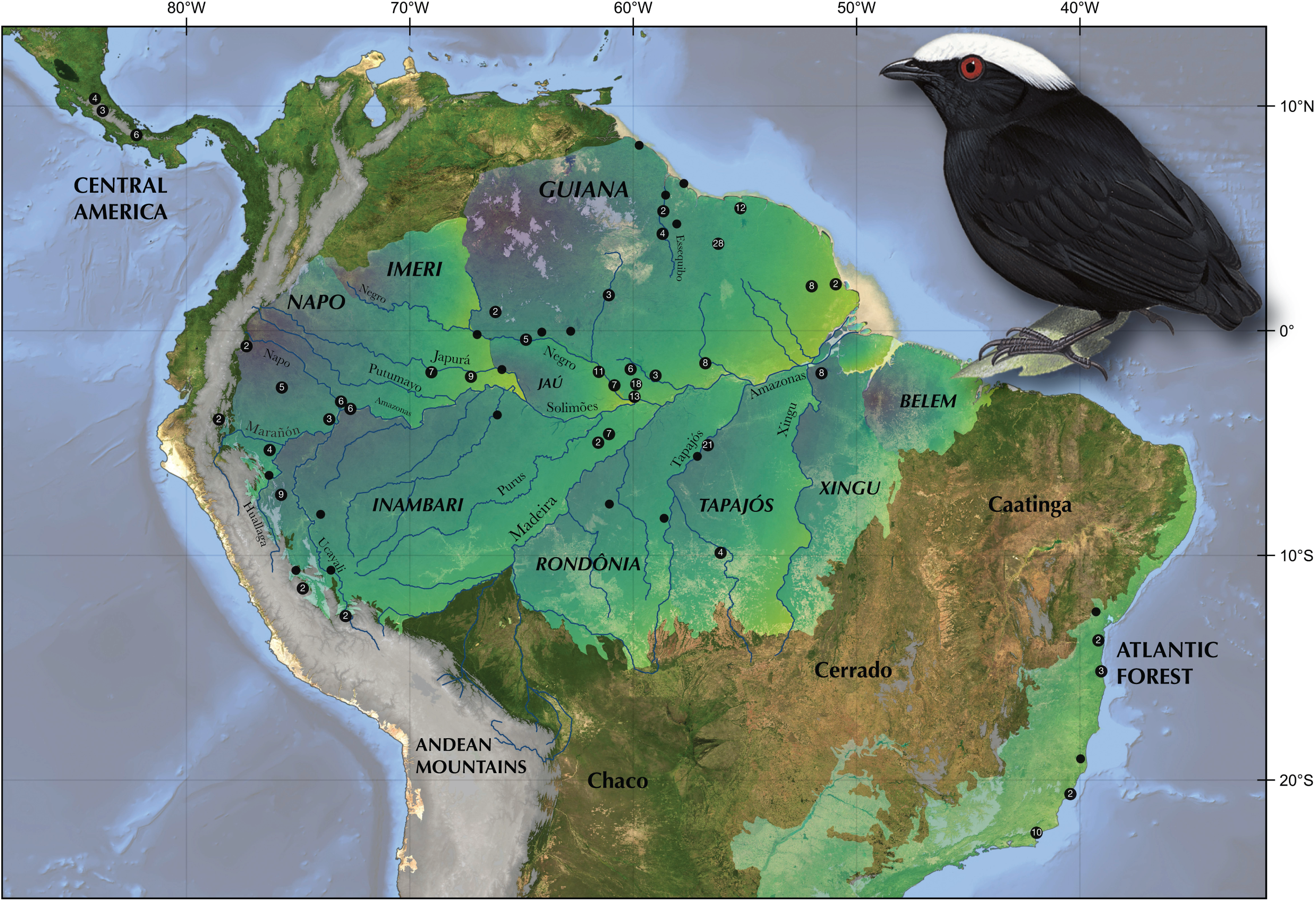
Neotropical areas of endemism and sampling regions. Amazonian lowland areas of endemism as portrayed in Da Silva et al. (2005) are emphasized with colored gradients, as well as the Jaú area of endemism (Borges and Da Silva, 2012). Montane Andean, Central American, Guianan, and dry diagonal regions (> 1000m) are emphasized in grey. As summarized in Da Silva et al. (2005), a sequence of authors identified seven areas of endemism for lowland birds that retained Wallace’s “Guyana” (Wallace, 1854), split “Ecuador” into “Imeri” and “Napo,” renamed “Peru” to “Inambari,” and split “Brazil” into “Rondônia,” “Pará,” and “Belém” (Cracraft, 1985; Cracraft and Prum, 1988; Haffer, 1978; Haffer, 1985; Haffer, 1992). “Pará” was subsequently further partitioned into two regions, “Tapajós” and “Xingu,” (Da Silva et al., 2002). Recently, additional sub-partitions have been proposed for the “Napo” (Jaú - Borges and Da Silva, 2012), and Guianan areas of endemism (Naka, 2011). Plotted markers indicate sampling areas for 286 *Pseudopipra* individuals sequenced for genetic analysis in the present study. The numbers inside markers indicate the number of individuals sampled within 0.25 degrees radius from a given midpoint coordinate (representing samples within a ∼50km diameter circle). Unlabeled markers indicate one sample from a region. Additional sample locality data is provided in Figure 2 and Supplementary Table 1. Inset top right is an illustration of the nominate *pipra* subspecies, reproduced from the Handbook of the Birds of the World, Lynx Edicions. See the Supplementary Data for a high-resolution version of this figure.

Montane Andean regions in the Neotropics are known to be exceptionally biodiverse, encompassing at least 15 areas of avian endemism with biotas shaped by a combination of vicariance and dispersal events (Hazzi et al., 2018; Morrone, 2015). Hazzi et al. (2018) concluded that the Marañon, Apurimac, and Yungas Inter-Andean Valleys, as well as the Colombian Massif, Las Cruces Pass, and the Táchira Depression have all likely played significant roles in structuring Andean biodiversity. Elevational gradients in the Andes also contribute substantially to Neotropical diversification metrics and raise fundamental questions about the historical relationships between lowland and montane endemics in the Neotropics (Janzen, 1967; Musher et al., 2019; Quintero and Jetz, 2018; Weir, 2006). From a biogeographic perspective, avian evolution has proceeded both “into and out of the Andes” (Brumfield and Edwards, 2007; Nylander et al., 2008). A number of examples illustrate colonization of Andean forest regions by lowland forest ancestors (Bates and Zink, 1994; Fjeldså, 1992; Lutz et al., 2013; Ribas et al., 2007). By contrast, some examples of diversification into the lowlands from Andean progenitors are associated with open habitat taxa (da Silva, 1995; van Els et al., 2019; Voelker, 1999), though see Thom and Aleixo (2015). For some tanagers (Aves: Thraupidae), the Northern Andes have been a source of lineages that later dispersed into the Central Andes and Amazonian lowlands (Sedano and Burns, 2010). Notably, Southern Andean forests may have been connected to the Atlantic Forest via Pleistocene expansions of forests into the region occupied today by the Cerrado biome (Cabanne et al., 2019). Indeed, glacial cycles and climatic fluctuations during the Pleistocene have been implicated as an important factor in montane diversification, disproportionally affecting high elevation forest (Hooghiemstra and Van der Hammen, 2004; Weir, 2006) and transiently connecting highland and lowland habitats (Brumfield and Edwards, 2007; Cabanne et al., 2019; Nylander et al., 2008).

In the lowland Amazon basin, a number of areas of endemism have also been described, delimited primarily by major tributaries of the Amazon river (Borges and Da Silva, 2012; Cracraft, 1985; Da Silva et al., 2005; Haffer, 1974). Wallace (1854) initially suggested that the Amazon basin could be divided into four wide bioregions (which he termed “Guyana,” “Ecuador,” “Peru,” and “Brazil”), based on primate distributions (see also Lynch Alfaro et al., 2015). These areas were subsequently partitioned by later biogeographers into at least eight major areas of endemism for terrestrial vertebrates (Figure 1) (Borges and Da Silva, 2012; Cracraft, 1985; Cracraft and Prum, 1988; Da Silva et al., 2002; Da Silva et al., 2005; Haffer, 1978; Haffer, 1985; Haffer, 1992; Naka, 2011). The aggregate of these geographic partitions has been recognized as the “Amazonian areas of endemism” and is the basis for the riverine barrier hypothesis (Figure 1, Antonelli et al., 2018; also see Figure 1A in Silva et al., 2019).

Few studies of birds have applied next-generation sequencing datasets to reconstruct the biogeographic history of a Neotropical radiation distributed across lowland and highland areas of endemism. Here, we investigate lineage diversification across the Neotropics in the continentally distributed *Pseudopipra* genus. The monotypic genus *Pseudopipra* (Kirwan et al., 2016), family Pipridae, currently includes a single species, the White-crowned Manakin (*Pseudopipra pipra* = *Dixiphia*). *Pseudopipra* is found across a variety of known dispersal barriers, including elevational gradients of the Andes, major Amazonian rivers, the dry diagonal, tepuis, and the Isthmus of Panama. Therefore, this group is particularly appropriate for assessing patterns of phylogenomic differentiation across a nested set of spatial scales in Neotropical forests.

In order to assess the degree to which population structure may be related to landscape features of the Neotropics, and the historical relationships among populations, our study design uses fine-scale sampling across a majority of *Pseudopipra’s* geographic distribution. We use double digest restriction site associated DNA (RAD) sequencing (Peterson et al., 2012) to subsample the genomes of hundreds of individuals spanning many well-known Amazonian and Andean areas of endemism, from Costa Rica to the Atlantic Forest of Brazil. We use these data, as well as a sample of mitochondrial sequences, to infer the history of lineages and populations within *Pseudopipra* using a variety of population genomic, landscape genetic, phylogenetic, and comparative approaches. Lastly, we investigated congruence between geographic variation in vocalizations, plumage phenotype, and phylogeographic structure, to evaluate the biological mechanisms that may be contributing to population differentiation and gene flow, and to propose a new taxonomic hypothesis for the genus.

### 1.1. Study organism

The White-crowned Manakin, *Pseudopipra pipra* (Pipridae), is a small (10-12 gram), non-migratory, suboscine passerine bird that is broadly distributed within Central America, the lower montane cloud forests of the Andes (up to ∼2000m) from Colombia to central Peru the Amazon basin, and the Atlantic Forest (Figure 2) (Kirwan et al., 2016; Kirwan and Green, 2012). The male’s striking white crown and jet-black body makes them among the most easily identified manakins, though the grey-green females are often confused with other manakin species in the field. *Pseudopipra* are typically found in dense humid forest and are predominantly frugivorous. Unlike many manakin species that exhibit concentrated or cooperative lek behavior (e.g. Prum, 1990; Prum, 1994), *Pseudopipra* males display in dispersed leks of 2-5 males (Castro-Astor et al., 2007; Snow, 1961). Castro-Astor et al. (2007) described at least 11 components in the display repertoire of the Atlantic forest *Pseudopipra* population, including rapid turning, jumping, “to-and-fro” flights, and an about-face.

**Figure 2.**
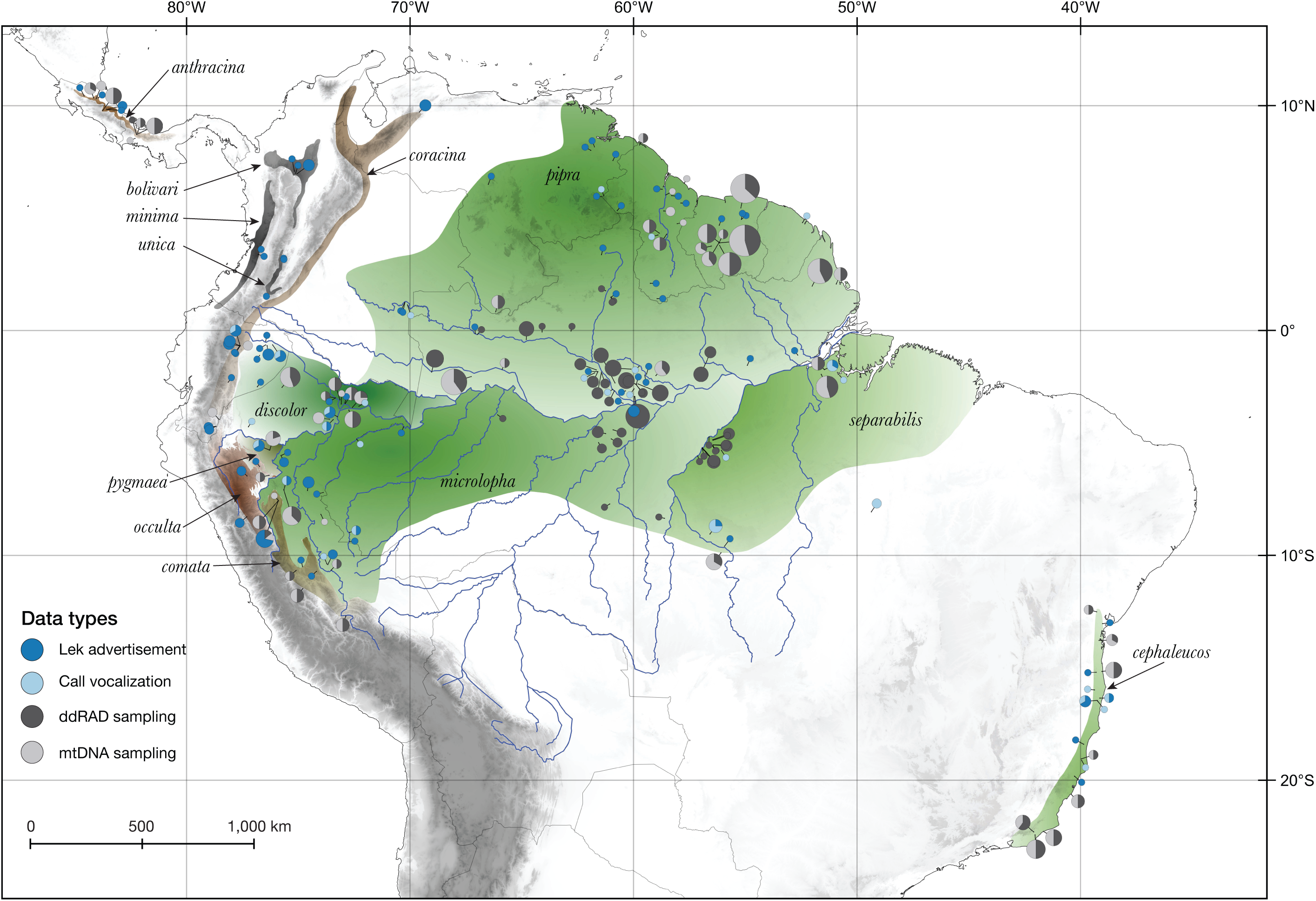
Data type by subspecies. Our total sampling of genetic and vocal diversity across *Pseudopipra* intersects with all recognized subspecies and represents 542 separate data records. Hypothetical distributions are based on descriptions in Kirwan and Green (2012) and indicate maximal ranges which emphasize physical barriers proposed to delimit subspecies (gradients of green: lowland taxa; gradients of brown: highland taxa). Montane areas are emphasized in progressively darker grey colors; Amazonian rivers are indicated in light blue. The overall distribution reflects an approximation from BirdLife International, updated to reflect our best knowledge at the time of writing (BirdLife International Species factsheet: *Pseudopipra pipra*, 2018, also see Supplemental Figure 1). The distributions of northern Andean taxa have been adjusted based on unpublished data provided by Andrés Cuervo (*personal communication*). We use pie charts to present the proportion of records of a particular data type at a given locality (light grey: mtDNA, dark grey: ddRAD, dark blue: lek vocalization, and light blue: call vocalization). Pie chart markers are connected to sampling locality coordinates with black leader lines. Pie charts are sized proportionally (max.: 22 records, min.: 1 record). Pie slices partition records of different types that may come from the same individuals: i.e., a pie chart with 50% dark grey and 50% light grey indicates only that an equal proportion of ddRAD and mtDNA samples are recovered from a locality (see Supplementary Table 1 for additional detail). Also note one confirmed vocalization record from southeast Amazonia outside the typical distribution at time of writing. See the Supplementary Data for a high-resolution version of this figure.

The two most recent phylogenetic hypotheses of manakins based on mitochondrial and nuclear DNA sequences both placed *Pseudopipra* as sister group to the five species of *Ceratopipra* (Leite et al., 2021; Ohlson et al., 2013). While the higher level phylogeny and taxonomy of manakins has received significant attention (also see Tello et al., 2009), work exploring intraspecific genetic variation within manakins has been restricted to relatively few species (e.g. Brumfield et al., 2008; Capurucho et al., 2013; Cheviron et al., 2006; Cheviron et al., 2005; Francisco et al., 2007; Gubili et al., 2016; Luna et al., 2017; McDonald, 2003; Reis et al., 2020).

Based on geographic variation in plumage and vocalizations, many previous authors have suggested that the biological species *Pseudopipra pipra* likely includes multiple, distinct phylogenetic species (Freile, 2014; Kirwan and Green, 2012; Ridgely and Greenfield, 2001; Ridgely and Tudor, 2009; Snow, 2004; Spencer, 2012). Thirteen subspecies of *Pseudopipra pipra* have been recognized based primarily on subtle variations in plumage coloration, which are frequently more marked in females than in males (summarized in Dickinson, 2003; Kirwan and Green, 2012; Snow, 2004). Because suboscine passerines generally do not learn their songs (but see Saranathan et al., 2007), the presence of substantial vocal variation across *Pseudopipra* populations further suggests that the genus may contain unrecognized cryptic species (e.g. Campagna et al., 2012). However, all previous authors awaited new information on genetic differentiation within *Pseudopipra* before making taxonomic recommendations.

It also remains unclear how the current subspecific classification reflects evolutionary history. The genetic variation within *Pseudopipra* has received prior attention by two studies that used mitochondrial DNA to infer population genetic patterns. Milá et al. (2012) reported intraspecific divergences of up to 3.5% (n=19) across three Amazonian populations, and the highest observed nucleotide diversity (π = 0.266) among 14 widely distributed Amazonian birds species. Castro-Astor (2014) used a larger sample (n=57), and discovered that at least four *Pseudopipra* subspecies correspond to well supported mitochondrial clades which were generally congruent with Amazonian areas of endemism. Castro-Astor (2014) used a molecular clock model to estimate the age of the *Pseudopipra* complex to coincide with the onset of the Pleistocene ∼2.457 Ma (95% HPD 1.45-3.97).

## 2. Material and methods

### 2.1. Field and tissue sampling

Muscle tissue samples were obtained from available vouchered avian material from US and Brazilian collections and other institutions (see Acknowledgements and Supplemental Table 1). We sampled from ten of thirteen recognized subspecies (Figure 2) of *Pseudopipra* (Kirwan and Green, 2012). Unfortunately, we were unable to obtain material of adequate quality for ddRAD sequencing from the Andes of Colombia (*bolivari*, *unica*, *coracina*, *minima*), Ecuador (*minima*) and the lowlands of Peru (*pygmaea*). However, we were able to obtain mtDNA data from *coracina* and *pygmaea*, which allowed us to make a preliminary assessment of their phylogenetic affinities (see discussion on mtDNA). In sum, after discarding failed samples and samples with high proportions of missing data, we obtained genetic data from 286 individuals (234 from ddRAD, 168 from mtDNA), representing ∼80 geographic regions and 10 of 13 subspecies. We also obtained comparable ddRAD and mtDNA data for two specimens of *Ceratopipra rubrocapilla* as outgroups (Supplementary Table 1).

### 2.2. Laboratory methods and data processing

We extracted DNA from avian tissue specimens (Supplementary Table 1) using the DNeasy Blood & Tissue Kit (QIAGEN, CA) and generated ddRADtags following the protocol of Peterson et al. (2012), with modifications described in (Thrasher et al., 2018). We used the STACKS 1.44 bioinformatics pipeline (Catchen et al., 2013) to generate assemblies of ddRAD loci, using *Manacus vitellinus* as a reference genome (GCA_001715985.1 from www.ncbi.nlm.nih.gov). We obtained mitochondrial ND2 sequences for a set of individuals using standard Sanger Sequencing protocols described in Berv and Prum (2014). Additional sequencing, assembly, and alignment details are reported in the Supplemental Appendix.

For ddRAD loci, we used several filters in the “populations” module from STACKS to generate different sets of bi-allelic SNPs: a missing data filter, a minimum depth of coverage filter, a filter that exports only the first SNP per RAD locus, and a minor allele frequency filter (MAF). In order to explore the sensitivity of genetic cluster inference to these important parameters (Linck and Battey, 2019; Paris et al., 2017), we produced eight versions of the SNP dataset by combining these different standard filters (Table 1).

**Table 1.** Summary of dataset filtering parameters. We generated eight SNP (single nucleotide polymorphism) datasets under different filtering strategies to evaluate the sensitivity of cluster analyses to these parameters. For eight SNP datasets, dataset D1 – D4mod, we note the parameters for each filter: the required sequencing coverage, the missing data threshold, whether or not a single SNP or all SNPs per locus are retained, the minor allele frequency cutoff (MAF), the number of loci, the total number of SNPS represented in a given dataset, and the average number of SNPs per locus.

| **Dataset** | **coverage** | **SNP present in** | **linkage** | **MAF** | **loci** | **SNPs** | **SNPs/locus** |
| --- | --- | --- | --- | --- | --- | --- | --- |
| D1 | >20x | >80% | 1st SNP | >= 5% | 1960 | 1960 | 1 |
| D1mod | >20x | 100% | 1st SNP | >= 5% | 175 | 175 | 1 |
| D2 | >20x | >80% | 1st SNP | NA | 2581 | 2581 | 1 |
| D2mod | >20x | 100% | 1st SNP | NA | 250 | 250 | 1 |
| D3 | >20x | >80% | all SNPs | >= 5% | 1960 | 5099 | 2.6 |
| D3mod | >20x | 100% | all SNPs | >= 5% | 177 | 391 | 2.2 |
| D4 | >20x | >80% | all SNPs | NA | 2581 | 41024 | 15.9 |
| D4mod | >20x | 100% | all SNPs | NA | 251 | 3832 | 15.3 |

Dataset 1 (D1-labeled arbitrarily) included 1,960 SNPs that were generated by setting the MAF to >= 5%, requiring minimum coverage of 20x, requiring the presence of a SNP in at least 80% of individuals, and filtering out all but the first SNP in each RAD locus to minimize linkage across the dataset. To evaluate the sensitivity of cluster analyses to rare variants (Linck and Battey, 2019; Shafer et al., 2017), Dataset 2 (D2) was exported with identical parameters to dataset D1, but without a minor allele frequency filter; this generated a dataset of 2,581 putatively unlinked SNPs. We generated datasets D3 and D4 to reflect the same filters as D1 and D2 but to also include all SNPs from each locus. Lastly, for each dataset D1-D4, we generated a complimentary dataset which filtered out any sites with missing data. The parameters of these SNP datasets are summarized in Table 1.

### 2.3. Phylogenetic analysis

Double-digest restriction site associated DNA sequence datasets are often characterized by high amounts of missing data (e.g. Eaton et al., 2017; Huang and Knowles, 2016; Shafer et al., 2017). Because phylogenetic analysis of sparse super-matrices may be prone to systematic error (e.g. Roure et al., 2013), we explored the sensitivity of our phylogenetic inferences to missing data using three different diplotype sequence datasets with 20%, 50%, and 80% thresholds for missing data at the RAD locus level. For example, for an 80% threshold, if the RAD locus is present in at least 20% of individuals, then that locus, containing all variable and invariable sites, was exported. This approach generated datasets comprising 2,584, 4,763, and 7,901 RAD loci, respectively. For each dataset, we generated a concatenated supermatrix using Sequence Matrix (Vaidya et al., 2011). These combined datasets had up to ∼40% missing sites across an entire matrix, irrespective of columns or rows (i.e., using the 80% threshold above).

We estimated phylogenies using each dataset with a concatenated maximum likelihood and a coalescent species-tree approach for a total of six phylogenetic analyses. For analysis of concatenated data, we estimated a maximum likelihood phylogenetic hypothesis using RAxML 8.2.9 (Stamatakis, 2014). We estimated 500 bootstrap replicates followed by a full ML search for each dataset, using a GTRGAMMAX substitution model. We avoid the need for ascertainment bias correction because we use whole short-read alignments, including invariant sites (Leaché et al., 2015). In our case, each alignment was 132 bp long (Supplemental Appendix). We also tested for an effect of data partitioning on phylogenetic inference by generating optimized locus partitioning schemes in PartitionFinder 2 (Lanfear et al., 2017), which were then re-analyzed in RAxML. We found that our topological results were insensitive to data partitioning, and as such we do not present these results. Lastly, we generated a mitochondrial DNA ND2 gene tree using IQ-TREE 1.6.10 (Chernomor et al., 2016; Hoang et al., 2017; Kalyaanamoorthy et al., 2017; Schmidt et al., 2014; Trifinopoulos et al., 2016). We partitioned by codon position and generated a maximum likelihood tree using the MFP+MERGE model search and partitioning option (Supplementary Figure 17).

We evaluated coalescent phylogenetic structure using the SVDquartets method implemented in PAUP* 4.0a159 (Chifman and Kubatko, 2014, 2015; Swofford, 2002). SVDquartets may be a useful tool for testing hypotheses of species delimitation because it can generate a lineage-tree under the multispecies coalescent without assigning individuals to species *a priori*. In this approach, each individual in a dataset is considered to be a separate “species.” If a recovered clade in such a lineage-tree has high bootstrap support, this would mean that the descendant taxa have strong support as a group under the multispecies coalescent, which may coincide with species boundaries (*Personal communication*, Laura Kubatko). Our SVDquartets analysis evaluated all possible quartets with 500 bootstrap replicates (other settings left to defaults). SVDQuartets has the advantage of being able to work directly on sequence data (above) and has been shown to perform well in comparison to summary methods which assume fully and correctly resolved gene trees as input (Chou et al., 2015; Schmidt-Lebuhn et al., 2017). SVDquartets has also been shown to be theoretically robust to variation in effective population size, molecular clocks, limited gene flow, and incomplete lineage sorting (Long and Kubatko, 2018).

While a full exploration of our specific usage of SVDquartets is beyond the scope of the present work, we suggest that this approach may be a useful way to assess support for hypotheses of species delimitation under the multispecies coalescent. We do not take the perspective that all well supported clades from this coalescent assessment *necessarily* represent biological species – such an interpretation often leads to an over-inflation of species estimates (Chambers and Hillis, 2020; Hillis, 2020; Sukumaran and Knowles, 2017). In general, we prefer a species agnostic approach to those which require *a priori* delimitation of species boundaries. Our approach instead focuses on discovering well supported coalescent structure without a prior hypothesis of species delimitation, and then assesses the degree to which metrics of support across coalescent and concatenated tree inference methods are concordant. We note that there are many approaches for species delimitation using genetic data (e.g. BP&P, Flouri et al. (2018), BFD, Leaché et al. (2014), PTP, Zhang et al. (2013), also see Leaché et al. (2019)), and that there is lack of consensus on how best to interpret these methods with respect to various species concepts (Carstens et al., 2013). In the present study, we rely on analysis of vocal and plumage phenotypes in combination with genetic patterns, to make taxonomic recommendations (Cadena and Zapata, 2021; Remsen, 2005).

### 2.4. Descriptive statistics

We estimated global and population pairwise F_st_ (Weir and Cockerham, 1984), regional variation in allelic richness (Goudet, 2005), Hardy-Weinberg equilibrium (Paradis, 2010), inbreeding (Goudet, 2005), linkage disequilibrium (Kamvar et al., 2014), sex-biased dispersal (Goudet et al., 2002), and performed an analysis of molecular variance (Kamvar et al., 2014) in *R* (R Core Team, 2018). Some of these statistics and analytical details are reported as supplementary material, while others are discussed below when relevant.

### 2.5. Cluster inference and patterns of genomic admixture

We performed cluster analyses intended to quantify the number of similar genetic groups across the range of *Pseudopipra.* First, we evaluated broad-scale variation across our genomic data using phenetic clustering implemented in the find.clusters function in the adegenet *R* package (Jombart, 2008). This function first transforms the data with a PCA, and then identifies clusters with a K-means algorithm (MacQueen, 1967). Using each SNP dataset, we evaluated K1:20 successively with the Bayesian Information Criterion (BIC). K-means clustering is based on the idea of minimizing the variance within clusters and has the potential advantage of making few assumptions about the underlying processes which generated the data. For each analysis, we retained all PC axes, set 200 randomly chosen starting centroids, and ran each search for 10^9^ iterations.

We also estimated the number of differentiated populations using STRUCTURE 2.3.4 (Pritchard et al., 2000), a widely-used model-based approach which assigns individuals to populations such that Hardy-Weinberg equilibrium is maximized and linkage among loci is minimized within groups. By comparing the output of STRUCTURE runs fixed to different numbers of populations (K), one can assess the degree of fit of the data to various models of K and assess genomic admixture among individuals. We ran STRUCTURE using SNP dataset D1 (Table 1), from K1:20 with 10 iterations per K value. We ran the program for 700,000 generations, discarding the first 200,000 as burn-in, implementing the admixture ancestry model with correlated allele frequencies. We evaluated K by examining Pr(*X*|*K*) or *L*(*K*) after summarizing our results using Structure Harvester (Earl and vonHoldt, 2012) and CLUMPP (Jakobsson and Rosenberg, 2007). We also used the Evanno method to estimate the rate of change in the log probability of data between values of K, which is a useful metric to detect the uppermost hierarchical level of structure in the data (Evanno et al., 2005). Lastly, we generated 95% Bayesian Credible Intervals around admixture coefficients (presented as averages across individuals for population assignment).

To investigate finer scale population genetic differentiation and the genomic composition of populations defined at different hierarchical levels, we used the programs fineRADstructure and RADpainter v.0.2 (Malinsky et al., 2018). RADpainter takes advantage of the information in haplotype data and considers each individual in a dataset as a “recipient” whose genomes are reconstructed using chunks of “donor” DNA from all other available individuals. This approach generates a “coancestry matrix” which combines the information that can be derived from both PCA and model-based clustering approaches and thus can be more sensitive to subtle population structure (Lawson et al., 2012). The fineRADstructure software uses an MCMC approach to explore the space of population assignments based on the coancestry matrix, using an algorithm which merges and splits populations, or moves individuals among populations. We ran fineRADstructure with default priors, but increased the burn in to 200000 iterations, followed by 1000000 iterations, sampling every 1000. We then assessed convergence by 1) considering the assignment of population membership across multiple independent runs, 2) visualizing the MCMC traces of estimated parameters to ensure convergence on the same posterior distributions, and 3) running each chain long enough to achieve effective parameter sample sizes > 100.

Both RADpainter and fineRADstructure are based on the chromoPainter and finestructure software (Lawson et al., 2012), but are optimized to take advantage of the linkage properties of RAD-like datasets. As our data are assembled relative to a *Manacus* reference genome, the order in which loci appear in our data files are related to positions on assembly scaffolds, even though the ddRAD loci in our data appear unlinked when considered at the population level (see Supplementary Appendix). We compared the inferred population assignments from fineRADstructure and STRUCTURE with K-means clustering (as above) of the coancestry matrix as well as SNP datasets. To determine how much of the variation in our dataset was captured by the RADpainter analysis relative to a standard PCoA of SNP data, we used the normalized PCA approach of (Lawson et al., 2012), using the mypca function provided in the “FinestructureLibrary.R” library in the fineRADstructure package.

### 2.6. Phylogenetic reticulation in the western Napo lineage

Model-based, phenetic, and population genetic analyses revealed that a lineage of individuals presently restricted to the western Amazonian Napo area of endemism has a complex history of introgression between southern and northern Amazonian clades (see Results). To further investigate this subset of individuals and their history, we generated two additional versions of the dataset optimized for different analyses. First-we generated a haplotype dataset of 52 individuals, which included all putatively introgressed western Napo individuals (n=10, clade E), as well as samples from proximate populations. These populations consist of a geographically adjacent population with predominantly unadmixed ancestry (Southwestern Amazon: Inambari, n=17, clade C1), and 2), and the source of introgressed genotypes (north-central Amazonian Jaú + eastern Napo, n=25, clade D2, as indicated by admixture analysis). We used a 5% minor allele frequency cutoff, required 20x minimum coverage, and allowed up to 20% missing data per locus—generating a dataset comprising 2,370 loci, including 4,979 SNPs (Dataset H1). Dataset H1 is analogous to that generated for coancestry (fineRADstructure) analysis of the entire dataset (above). We also generated a similar dataset (Dataset H2), without a minor allele frequency cutoff—comprising 2,947 loci including variable and invariable sites—as required for inference of demographic parameters (Gronau et al., 2011).

Hybrid populations are expected to have higher mean coancestry and lower F_st_ with each of their progenitor lineages than their progenitor lineages will have with each other (Barrera-Guzmán et al., 2018). To test this prediction, we first estimated coancestry values using Dataset H1 with RADpainter (as above). We then compared inferred coancestry of population pairs with a standard ANOVA and the glht function in the multcomp R package (Hothorn et al., 2013). We estimated F_st_ with Weir and Cockerham’s estimator (Weir and Cockerham, 1984), and tested the prediction using 1000 bootstrapped datasets to estimate 95% confidence intervals.

To test whether or not the relationships among these focal populations is best represented by a phylogenetic network, we used PhyloNet 3.6.4 (Wen et al., 2018; Zhu et al., 2018). PhyloNet takes SNP data and uses a reversible-jump MCMC technique to explore the posterior distribution of phylogenetic networks and bi-furcating topologies, while accommodating both reticulation and incomplete lineage sorting. PhyloNet searches the set of all possible reticulation models without requiring *a priori* model specification.

For computational tractability, we used SNP dataset D1 (1960 unlinked SNPs), and pruned it to the 52 individuals comprising the groups of interest. We pruned out all sites with missing data, leaving 572 unlinked biallelic SNPs, as required by PhyloNet. For our final runs of the PhyloNet program MCMC_BiMarkers, we randomly subsampled five diploid individuals from each group, as preliminary program runs indicated a severe computational bottleneck with more than five diploid individuals or with larger datasets. We assigned each group of individuals to a lineage, allowed for a maximum of one reticulation, set the chain length for 800000 iterations (sampling every 100), set a burn in of 2.5%, and left all priors as default (i.e., assuming population mutation rates are constant).

Using the final network topology estimated from PhyloNet, we estimated demographic parameters using G-PhoCS version 1.3 (Gronau et al., 2011) with dataset H2. We ran the program for 750,000 iterations with a 10% burn-in, estimating a total of 13 parameters: the effective population sizes of the three focal lineages, two ancestral population sizes, two splitting times, and six directional migration parameters. Parameter MCMC traces were inspected in Tracer 1.7.1 (Rambaut et al., 2018). We converted the median and 95% Bayesian credible intervals from mutation scale to generations and individuals as described in Campagna et al. (2015), assuming an approximate mutation rate of 10^-9^ per base pair per generation (Kumar and Subramanian, 2002; Smeds et al., 2016). We calculated the number of migrants per generation as the product of the per generation migration rate multiplied by ¼θ for the receiving population (m_a-b_ * (θ_b_/4)) (Gronau et al., 2011), and considered migration rates to be significant if the 95% lower bound was > 1 (the level of gene flow expected to counteract the effects of drift; Wright, 1931).

### 2.7. Spatial distribution of genetic variation

Dispersal barriers often create sharp genetic discontinuities between adjacent populations that would otherwise be continuous (Petkova et al., 2015). To assess the degree to which spatial variation in genetic diversity within *Pseudopipra* may be attributed to landscape features, we investigated how discrete and continuous patterns of isolation-by-distance (IBD) vary across the landscape. First, we used Mantel and partial Mantel tests to investigate IBD effects within sampling regions delimited by prominent physical barriers, and then separately tested the roles of specific dispersal barriers in structuring genetic variation. Mantel tests evaluate the correlation among two or more matrices, in our case representing pairwise genetic and geographic distance. Significance is assessed by permuting the rows and columns of one of these matrices (Mantel, 1967). Partial Mantel tests incorporate a third “barrier matrix,” which contains information about environmental or ecological distance. By evaluating the correlation between genetic and ecological distance while controlling for geographic distance, a partial Mantel test can be used to investigate the effect of a particular geographic barrier on genetic differentiation. Because mantel tests may have an inflated type I error rate (e.g. Bradburd et al., 2018; Guillot and Rousset, 2013; Legendre and Fortin, 2010), we follow the recommendations of Diniz-Filho et al. (2013), and only reject the null hypothesis of no correlation if *p* < 0.001 (∼10x more stringent than a standard correction for multiple tests would require).

For these tests and others, we generated a pairwise geographic distance matrix by calculating great circle distances using the distm function in the geosphere *R* package (Hijmans et al., 2015). Individuals sampled at the same GPS coordinates were set to have an arbitrarily small geographic distance of 0.0001 meters. In order to assess spatial variation *within* and *between* populations, we use estimates of pairwise genetic distance at the individual level, rather than at the population level. Our analyses compare patristic phylogenetic distance between all pairs of individuals (estimated from RAxML) as well as coancestry (from fineRADstructure), as proxies of genetic distance among individuals. These procedures, including the generation of an appropriate barrier matrix, are developed in a set of *R* functions and scripts that operate on adegenet GENIND objects (see supplementary *R* code) and internally use the Mantel implementation from the ecodist R package (Goslee and Urban, 2007).

We also analyzed continuous patterns of spatial variation in our data with a population genetic model that relates effective migration rates across geographic space to expected genetic dissimilarities among individuals (Petkova et al., 2015). This method, termed Estimated Effective Migration Surfaces (EEMS), produces a visualization that emphasizes deviations from isolation by distance to highlight migratory corridors and barriers to gene flow. We view this method as complimentary to more explicit Mantel tests, because it seeks to identify where the assumption of constant IBD is violated across continuous geographic space, and thereby can highlight areas where strong barriers to gene flow may exist. In brief, a region is covered with a dense regular grid connecting subpopulations (demes), among which individuals can migrate with rates varying by location. EEMS uses an MCMC approach to estimate expected genetic dissimilarity between two individuals, integrating over all possible migration histories, and adjusting migration rates among graph edges to match the genetic differences in the data. A migration surface is then interpolated across a region to indicate where genetic distances decay faster or slower than predicted by a model of isolation by distance, visually highlighting potential barriers to gene flow and migration corridors. An effective diversity parameter is also estimated for every deme, reflecting local deviations in heterozygosity. We ran several replicates using default priors, iteratively increasing the grid density and chain length, with up to 2000 estimated demes. For each test, we ran three chains, and after examining chain convergence, we assessed model fit by comparing the observed and fitted dissimilarity between and within demes.

### 2.8. Vocal variation

We obtained sound recordings of vocalizations of *Pseudopipra* from the Macaulay Library at the Cornell Lab of Ornithology (https://www.macaulaylibrary.org) and the Xeno-canto (www.xeno-canto.org) collections in May 2019 (Supplementary Table 4). Vocalizations were analyzed in Raven Pro 1.4 (Cornell University, 2011), and converted into spectrograms using a 512-sample Hann window with 50% overlap. Audio recordings ranged from a cut of a single vocalization to longer recordings of a lek with multiple individuals vocalizing. For our initial assessment, we identified qualitatively distinct vocalization types based on our own survey of the available recordings, without regard to the geographic location of the recordings or subspecies identity.

*Pseudopipra* vocalizations have 1-3 buzzy and/or tonal notes. We measured: 1) starting frequency, 2) ending frequency, 3) minimum frequency, 4) maximum frequency, 5) number of notes, and 6) duration of the entire vocalization (Supplementary Figure 13). We performed principal components analysis (PCA) and logistic regression on these vocal measurements to test for significant differences between vocal types and to reduce the dimensionality of the data for comparison to results from analysis of genetic data. Because no sound records were directly associated with genetic samples in this study, we used geographic proximity to vocalization recordings and localization to areas of endemism or areas bounded by clear physical barriers to associate vocal types to genetic samples and taxonomic ranks. This approach assumes that genetically and geographically proximate individuals are likely to share the same vocal type and enabled us to perform a preliminary assessment of how variation in vocalization type maps onto existing genetic variation. Testing the fine-scale association of genetic and vocalization boundaries will require extensive field sampling from individual manakins. Additional analytical details and analyses are reported as supplementary material.

**Figure 3.**
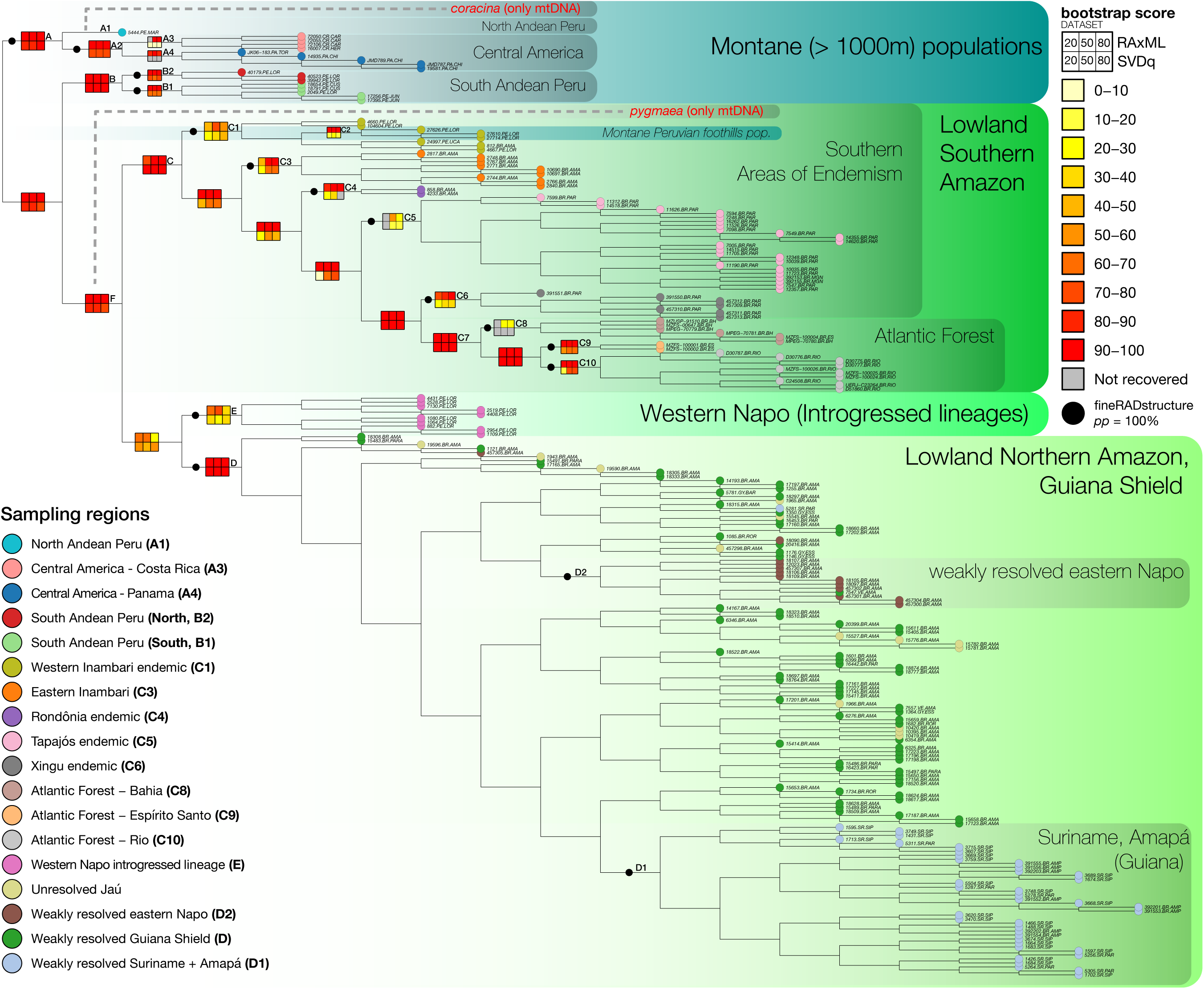
New phylogenetic hypothesis for *Pseudopipra*. Phylogenetic analysis of concatenated ddRAD sequences generated a well-resolved phylogenetic hypothesis which was largely congruent across datasets and analytical approaches. Montane clades, including a Northern Andean (A1) + Central American (A2) clade (A), and a central Peruvian clade (B), are recovered as nested sister groups to two wide ranging lowland clades (C and D), indicating that the lowland clades are descended from Andean lineages (Figure 5). Shown is the RAxML topology generated using the 50% diplotype dataset. Branch lengths are set to equal for display (see Supplementary Data for original tree files). Colored markers at tips indicate group membership to one of eighteen population-areas (matching Figure 4 and 5). Heatmaps at key nodes indicate bootstrap scores from each of six phylogenetic analyses for 20/50/80% datasets, as indicated in the legend. Black markers on branches indicate populations which are identified with 100% posterior probability in fineRADstructure analysis. Deep phylogenetic structure is recovered by both RAxML and SVDquartets with high support, though RAxML recovers additional low-support substructure in the Northern Amazonian + Guiana Shield clade (D) which is coincident with several geographic features (clades D1 and D2). A clade of introgressed western Napo individuals (E) is recovered by all standard phylogenetic analyses as sister to the northern Amazonian + Guiana shield clade (D). As the history of this clade is characterized by a complex introgression scenario between northern and southern lineages, a bifurcating tree model is inadequate to describe its relations with other groups (see text and Figure 6). Also shown are the well supported phylogenetic positions of two distinct subspecies lineages (*pygmaea* and *coracaina*) from which we were able to obtain mtDNA (Supplemental Appendix). Plotting was aided by the phytools (Revell, 2012) and ape (Paradis et al., 2004) *R* packages.

## 3. Results

### 3.1. Genetic *data collection*

Our total genomic data set included an average of 1 ± 0.4 million sequences of 138 bp from 232 individuals of *Pseudopipra* and two *Ceratopipra rubrocapilla,* which were used as an outgroup. Our reference-based assembly produced a catalogue with 47,046 RAD loci. Filtered SNP datasets requiring a SNP to be present in at least 80% of individuals ranged from 1,960 unlinked SNPs (dataset D1: 5% MAF with 1^st^ SNP on each locus) to 41,024 SNPs (dataset D4: all SNPs on all loci, no MAF) (Table 1). Datasets requiring a SNP to be present in 100% of individuals were considerably smaller, as expected, and ranged from 175 SNPs in D1mod, to 3,832 SNPs in D4mod (Table 1). We also obtained 168 mtDNA ND2 sequences using PCR methods which broadly overlapped with our ddRAD dataset. Our mtDNA dataset included two subspecies, *pygmaea* and *coracina*, from which we were not able to obtain enough high-quality material for ddRAD sequencing.

### 3.2. Phylogenetic analyses

RAxML and SVDquartets analyses generated a consistently well-supported phylogenetic hypothesis (Figure 3). The monophyletic lineages of *Pseudopipra* have distributions that are largely congruent with previously recognized avian areas of endemism, and further reveal much finer scale patterns of local biogeographic differentiation. Using the two *C. rubrocapilla* individuals to root the phylogeny (not shown), we detected support for at least five major clades across all phylogenetic analyses of nuclear genomic data, with most receiving high bootstrap support in concatenated or coalescent analysis (clades A-E in Figures 3-5). We first describe the structure of these major clades, followed by descriptions of well supported substructure within each of these regional groupings. We indicate six bootstrap values for each clade with at least some strong support, from each of the three datasets analyzed in RAxML and SVDquartets (see heatmaps, Figure 3). Phylogenetic analysis of mtDNA (Supplemental Figure 17) generated a congruent result, except where noted below.

#### 3.2.1 Phylogenetic structure

The sister group to all the other *Pseudopipra* is a clade including individuals from subtropical Central America and the northern Andes. The northern Andes are represented in our ddRAD dataset by a single individual from subtropical forests (∼1,000 m) of the eastern slopes of the Andes in San Martín, Peru. This subtropical San Martín specimen (identified as *P. pipra occulta*) was the only viable sample in our genomic dataset from the entire northern Andes region (A1 in Figure 3). Mitochondrial data from three additional specimens from the Andes of southern Ecuador (subspecies *coracina*) form a monophyletic group with the San Martín specimen; thus, this northern Andean sister lineage to the Central American lineage likely includes all montane Andean populations of *Pseudopipra* in Ecuador, Colombia, and Venezuela. A well supported Central American clade (A2 in Figure 3) was identified in all analyses, with Costa Rican (A3 in Figure 3) and Panamanian subclades (A4 in Figure 3) supported in RAxML.

The second successively nested clade in *Pseudopipra* includes individuals from subtropical forests of the Andes of Southern Peru, which are sister to all remaining lineages (Clade B in Figure 3). The southern Andean Peru clade is further subdivided into two very well resolved clades: a northern clade (B2 in Figure 3), sampled from the southern Cordillera Azul in southwestern Loreto, and a southern clade from Pasco, Junín, and Cusco (labeled B1 in Figure 3). This southern Andean Peru clade is the sister group to a monophyletic, lowland clade (F in Figure 3) that is further subdivided into a southern Amazon clade + Atlantic Forest clade (Clade C in Figure 3), and a Northern Amazonian + Guiana Shield clade (Clade D in Figure 3). Our analysis of mtDNA with sampling of subspecies *pygmaea*, from the tropical forest of the lower Huallaga river Valley, Peru, places this population as sister group to the entire lowland radiation (Clade F; Supplemental Appendix).

Within the broad lowland Northern Amazonian + Guiana Shield clade (Clade D), virtually no phylogenetic substructure is associated with strong bootstrap support. The maximum likelihood tree however does have substantial biogeographic coherence, which we use as additional justification for population subdivision when monophyletic groups are identified to be coincident with geographic features. For example, all but one individual from coastal Suriname and Amapá form a monophyletic assemblage (D1 in Figure 3), east of the Essequibo river. Likewise, all but one individual from the far eastern Napo area of endemism, east of the Putumayo river, form a clade (D2 in Figure 3). This coherence with landscape features is unlikely to be coincidental, though support for these groups is weak. We treat clade D1 and D2 as well as two other Guianan regions, as separate populations for descriptive population genetic statistics.

A widely distributed set of individuals encompassing the western portion of the Napo area of endemism (from the northern bank of the confluence of the Solimões river and the Napo river north of Iquitos, then west and south, Clade E in Figure 3) was recovered as the sister group to the Guiana Shield + Northern Amazon clade (Clade D in Figure 3). Analysis of mtDNA (Supplementary Figure 17) further resolved (BS > 99) two subgroups delimited by the Napo river. Importantly, and in contrast to nuDNA, we infer that these mtDNA haplotypes are derived from Inambari lineages (BS = 98, Clades C1 + C3 in Figure 3), and are not sister to Clade D. These conflicting patterns are explained by subsequent population genetic analyses, which indicate that this Napo population is a distinct lineage of the southern Amazonian clade (Clade C in Figure 3) that has experienced substantial hybrid introgression from the northern Amazonian clade D (see Population Genetic analyses and Discussion).

Within the large lowland Southern Amazonia clade (Clade C in Figure 3), we recovered eight moderately supported and hierarchically nested clades, which are successive sister groups to each other, and are congruent in distribution with recognized areas of endemism. First is a moderately supported clade (C1 in Figure 3) of individuals from the western Inambari area of endemism, from west of the Ucayali river to the Purus river. Although the first lineages to diverge within clade C1 have low bootstrap support, a monophyletic subgroup of three montane (>1000m) individuals from the highlands between the Huallaga river and Ucayali river cluster together with high bootstrap support (C2 in Figure 3).

The next successively nested clade includes individuals from eastern Inambari, between the Purus river and the Madeira river (C3 in Figure 3). The next successive clade includes two individuals sampled from the Rondônia area of endemism, between the Madeira river and Tapajós river (C4 in Figure 3). The next successive clade includes a group of individuals from the Tapajós area of endemism between the Tapajós river and the Xingu river (C5 in Figure 3). Next, a clade comprised of individuals from the northern Xingu area of endemism, east of the Xingu river (C6 in Figure 3), is sister to a well-supported Brazilian Atlantic Forest clade (C7 in Figure 3), which is further subdivided into three successive clades (north to south) in Bahia, Espírito Santo, and Rio de Janeiro, respectively (C8, C9, and C10 in Figure 3). In summary, the subdivisions of the Southern Amazonian clade reflect a stepwise west-east progression, with a hierarchically nested structure.

### 3.3. Cluster inference and patterns of genomic admixture

#### 3.3.1. Broad-scale population structure and admixture

Weir and Cockerham’s F_st_ was moderate when comparing populations within *Pseudopipra*; for Dataset 1, overall F_st_ was 0.195 [95% CI: 0.187-0.203], and never dipped below 0.171 [0.168-0.175] (Dataset 4). Notably, in SNP datasets filtered to remove sites with missing data, overall F_st_ estimates were on average ∼20% higher and ranged from 0.188 [0.175-0.201] (D4) to 0.261 [0.191-0.336] (D2). Pairwise population F_st_ estimates from Dataset 1 (other datasets showed similar patterns, see Supplementary Figure 10a-h), ranged from essentially undifferentiated, to almost entirely distinct. At the extremes: comparing the geographically proximate Jaú and Guiana Shield populations indicated a particularly low F_st_: 0.005 [0.003 - 0.007]. By contrast, comparing the Atlantic Forest Espírito Santo population to the Panama population recovered an F_st_ of 0.81 [0.79 - 0.83], or almost entirely differentiated. For reference, Hartl and Clark (1997) indicate F_st_ > 0.15 may be considered “great” genetic differentiation.

K-means clustering of SNP datasets on the basis of BIC score identified groups of individuals that differed somewhat with respect to dataset filtering strategies (Summarized in Figure 5, right). For D1, K-means clustering identified five major groups: Central America (Clade A2), South Andean Peru (Clade B), Atlantic Forest (Clade C7), Southern Amazon including the western Napo population (the rest of Clade C + Clade E), all Guiana Shield + Northern Amazon (Clade D) (Supplementary Figure 2a-h). For dataset D1, the montane sample from San Martín (North Andean Peru, 5444.PE.MAR) clustered with Central American populations (Clade A), as it does in all phylogenetic analyses. These clusters are essentially similar to those detected by STRUCTURE at K=5 (Figure 4, 5, below). All other SNP datasets grouped the San Martín specimen with the Southern Amazon populations. D1m further identified the Tapajós + Xingu individuals as a distinct cluster, while D2, D3, D3m, D4, and D4m generated sets of clusters identical to D1, with the exception of the clustering of the San Martín specimen, which clustered with the Southern Amazon in those datasets. Additional field sampling of the North Andean population is required to understand why this lineage may share signals of ancestry with northern and southern groups. K-means clustering of D2m failed to discriminate any lowland Amazonian populations (Supplementary Figure 2d). Thus, only 75 low frequency SNPs which are removed in D1m but are present in D2m obfuscate the signal of population differentiation detected by K-means clustering of D1m. Lastly, across datasets, standard PCoA explained 13-26% of the variance in the SNP data on the first two axes (Supplementary Figure 2).

**Figure 4.**
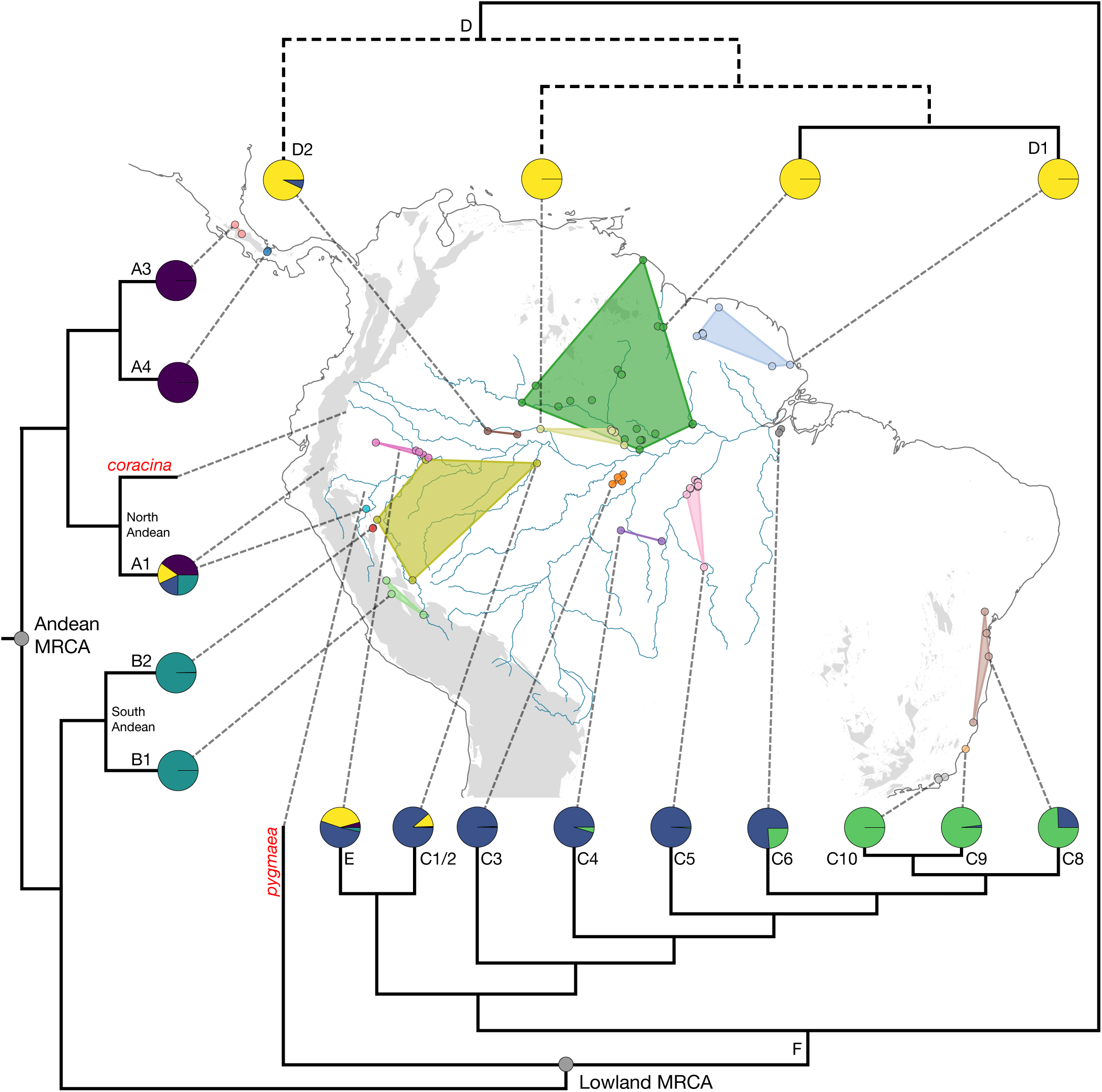
Mapping of phylogenetic relationships into geographic space. Here, a simplified topology is plotted with tips anchored to monophyletic sampling regions (except for clade D). The colors of each convex hull (i.e., hypotheses of minimum clade ranges) match tip marker colors in Figure 3 and 5. These colored hulls represent distinct non-overlapping geographic areas delimited by physical barriers. Note the two weakly resolved lineages (pale yellow Jaú, and green Guiana Shield population groups in clade D) are delimited by the Negro river. Clade D bipartitions are shown with dashed lines to indicate weak phylogenetic support (Figure 3, results). The ancestral habit of the genus is inferred to be Andean (Figure 5, Supplementary Appendix), with a single origin of lowland lineages. Also shown at the tips of the simplified tree are median admixture proportions across sampling localities as inferred with STRUCTURE (Figure 5).

**Figure 5.**
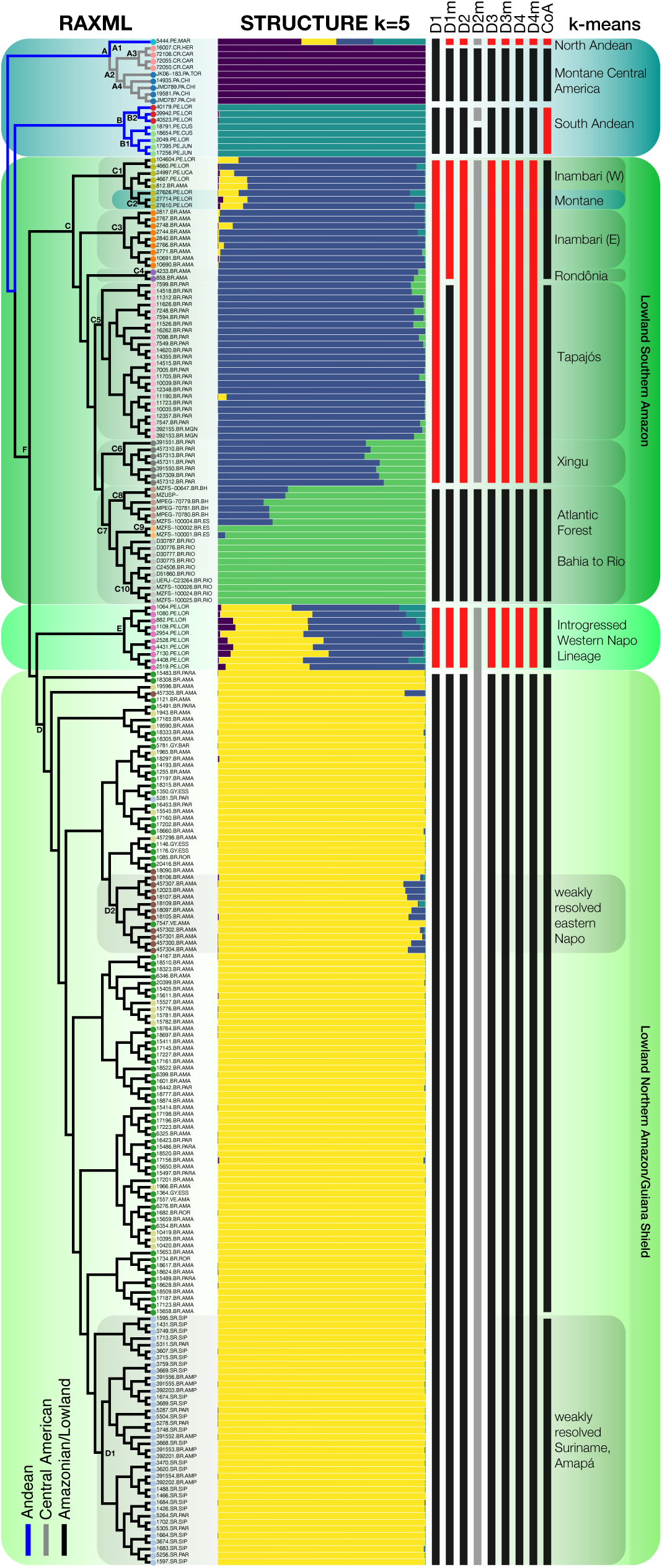
Cluster and admixture analysis of SNP data. Left: RAxML topology using the 50% diplotype dataset, with tip markers colored to indicate group membership to one of eighteen population-areas (matching those in Figures 3, 4). Branch colors indicate inferred ancestral areas, (see Supplemental Appendix and text). Center: STRUCTURE summary for Dataset 1, depicting population assignment and admixture estimates for K=5. The five groups correspond to wide biogeographic regions which coincide with lowland areas of endemism, as well as Central American, and Andean regions (Figure 4). Each labeled region is inferred to have a unique combination of admixture proportions. The admixture gradient inferred from the Xingu to the southern Atlantic Forest may be a product of isolation by distance (see Supplementary Appendix), whereas the signature of admixture inferred for western Napo lineages appears to be a product of historical introgression from a northern lineage, into a distinct Southern Amazonian lineage (see Figure 6 and discussion). See Supplementary Figure 3 for K2-10 results. Right: K-means clustering solutions for SNP dataset D1-D4m (described in Table 1, also see Supplementary Figure 2), as well as for the coancestry matrix (CoA). Black bars indicate distinct clusters which are also detected as monophyletic groups in the phylogenetic analysis. Red bars indicate one distinct K-means cluster from a given dataset, but which is paraphyletic with respect to the phylogenetic analysis. The grey bars for D2m indicate several groups of individuals that were not resolvable via K-means (Supplementary Figure 2). See the Supplementary Data for a version with individual sample IDs.

Visual inspection of Pr(*X*|*K*) from summarized STRUCTURE runs on dataset D1 indicated that the likelihood of each successive K from 1:20 plateaued at K = 5, with the standard deviation across runs increasing rapidly after this point (Supplementary Figure 3). We describe patterns of inferred admixture as they appear in Figure 5, reading from top to bottom. For K5, individuals from Central America are unambiguously assigned to their own cluster (Clade A2 in Figure 3, 5). The north Andean montane specimen from San Martín, (North Andean Peru, 5444.PE.MAR) is reconstructed as being highly admixed, with a genomic composition including 4 of the 5 inferred clusters. However, this inference is tenuous because we have only one individual to represent this likely distinct and highly diverse north Andean lineage (though see RADpainter below, which is consistent with these results). Samples from the southern Amazon and Atlantic Forest (Clade C), have no detected admixture with northern populations, except for some individuals from the western (Clade C1), and to a more limited degree, the eastern (Clade C3) Inambari populations, which are inferred to have limited (∼ <5%) admixture with individuals assigned to the Northern Amazon + Guiana Shield (Clade D, see Figure 5, 6, and discussion below).

**Figure 6.**
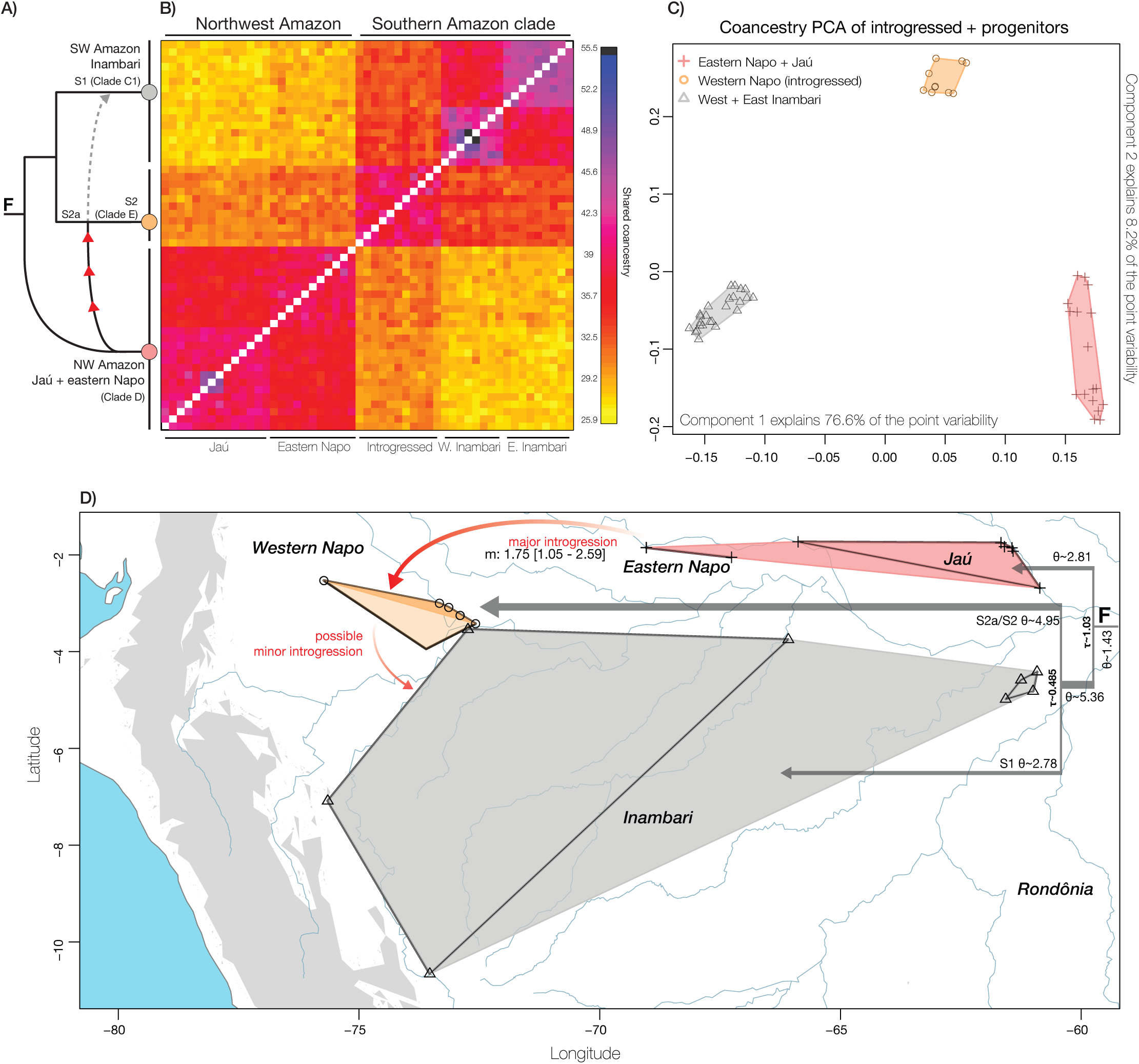
Exploration of the introgressed lineage in the western Napo area of endemism. The available genetic evidence appears consistent with the hypothesis that individuals in the western Napo area of endemism are derived from a distinct southern Amazonian lineage (S2a in Panel A). This lineage (S2a) experienced substantial introgression from the ancestral populations of what are today eastern Napo and Jaú restricted linages. A) depicts the phylogenetic network (in black) inferred by PhyloNet, with red arrowheads indicating a strong signal of asymmetric introgression indicated by G-PhoCS (clade C1 in Figure 3, into a southern Amazonian lineage S2a). The dashed grey line connecting S2a/S2 to S1 indicates a weakly supported hypothesis of introgression into Inambari lineages, suggested only by STRUCTURE (see text and Figure 5). B) displays the individual level coancestry matrix for introgressed Napo individuals and their hypothesized progenitor lineages. Introgressed hybrids (clade E, or abbreviated PH for putative hybrid) have greater coancestry with each of their progenitor lineages [S2a and Northern Amazon + Guiana shield (abbreviated GS)] than their progenitor lineages share with each other (PH vs. GS / progenitors = 1.078, PH vs. Inambari / progenitors = 1.17, see text). The southern progenitor lineage (S2a) is unsampled or extinct. C) summarizes coancestry variance as projected into component space after normalizing. D) summarizes the minimum implied geographic extents of the groups involved. For the convex hull representing the range of the western Napo lineage (beige), darker shading indicates localities for ddRAD sampling, while lighter shading indicates an additional locality of the Napo lineage from mtDNA samples. G-PhoCS demographic parameter estimates are also depicted in panel D (m: migrants per generation, when lower 95% CI > 1, τ: splitting time in generations, θ: effective population size (10^6^), Supplemental Table 5). In A-D, two subpopulations (East and West) are indicated for each progenitor group, as outlined with black convex hulls in D.

Going from West to East across the Southern Amazon, limited admixture with Atlantic Forest alleles (green, Clade C7) is first inferred in the Tapajós (Clade C5) area of endemism. This admixture component increases in the Xingu area of endemism (Clade C8) and increases further in a geographic and genetic cline to 100% assignment probability to a distinct cluster in the Rio de Janeiro clade (Clade C10). Given the geographic context of this cline in admixture, we suggest this pattern may a product of isolation by distance, hierarchical structure, and geographic barriers, and not isolated introgression events. We note that the conStruct method (Bradburd et al., 2018) may be useful for explicitly testing this hypothesis (Supplemental Appendix), however our evaluation of this method suggested that it is presently intractable for a dataset of this size.

Next, all individuals from the western portion of the Napo area of endemism are inferred to be heavily admixed (Clade E; Figure 5). On average, at K=5, STRUCTURE assigns 40.2% of these individuals’ (n=10) genetic composition to the Guiana Shield population (SD 4.7%, average Bayesian credible interval (aBCI): [0.33-0.47]), 50.4% to the Southern Amazon population (SD 3.4%, aBCI: [0.43-0.58]), as well as limited 4.15% probability to Central Peru (SD 3.1%, aBCI: [0.01-0.08]), and 5.2% to Central America (SD 4.8%, aBCI: [0.02-0.1]). The three localities included in this clade– (1) north of Iquitos, on the east bank of the confluence of the Solimões river and the Napo river, (2) southwest of Iquitos in Allpahuayo Mishana Reserve (represented by three mtDNA samples), and (3) near San Jacinto, Ecuador – indicate that this heavily admixed population (corresponding to subspecies *discolor*) may occur across a large area of the western Napo region of endemism, and therefore may not represent a narrow hybrid zone (Figures 2-7).

Lastly, at K=5, all members of the Guiana Shield + Northern Amazon clade (Clade D) are inferred to be virtually unadmixed. Individuals in the far eastern Napo area of endemism on the east bank of the Putumayo river (brown markers in Figure 3, 4) cluster with other Guiana Shield groups in phylogenetic analyses but are detected to have limited (∼5%) admixture with pure individuals of Southern Amazon provenance. At higher values of K, the broad-scale population assignments inferred at K=5 are mostly unchanged, however additional admixture components are inferred for most groups. The introgressed western Napo clade is eventually placed into its own cluster at K=9-10 (see Supplementary Figure 4, Supplementary Figure 8).

#### 3.3.2. Fine-scale population structure and admixture

Admixture analysis using the RAD locus optimized chromosome painting approach in RADpainter and then population assignment with fineRADstructure (Lawson et al., 2012; Malinsky et al., 2018), was congruent with other analyses, and also detected finer partitions of population structure. The fineRADstructure program detected 15 groups of individuals which were each supported with a posterior probability of 100% and which also overlapped with sampling regions which we previously delimited for population genetic analyses based on geographic barriers (Black markers in Figure 3, Supplementary Figure 5). The only sampling regions which fineRADstructure failed to confidently partition into separate groups consisted of a group of individuals in the Jaú area of endemism, and a group of individuals in the eastern Napo area of endemism (both groups are poorly resolved in phylogenetic analyses, but unambiguously members of the broader Northern Amazon + Guiana Shield clade D). The eastern Napo individuals (D2 in figure 3) are identified as a low support group (*pp* = 0.49), while the Jaú individuals are lumped in with other western Guiana Shield individuals (D in figure 3). See Supplementary Figure 5 (fineRADstructure dendrogram), and the full dataset coancestry matrix plots Supplementary Figures 6-7.

Patterns of coancestry show a complex mosaic of shared ancestry among populations (Supplementary Figures 6-7). In general, coancestry patterns appear consistent with admixture patterns detected by STRUCTURE, including the case of the highly introgressed western Napo lineage. Therefore, we take patterns indicated by STRUCTURE to be reliable. Notably, the high level of admixture initially inferred for our San Martín sample is recapitulated in RADpainter coancestry analysis – this sample has relatively high coancestry across most populations (as does the pattern of coancestry we report for the western Napo lineage, below).

Lastly, the Lawson et al. (2012) “normalized PCA” approach provided with fineRADstructure captured 89% of the variance in the genetic data on the first four component axes (51.4%, 24.9%, 7.77%, 4.94%) indicating that the coancestry matrix captures much more information about genetic variance than standard PCoA/PCA of SNP data (Supplementary Figure 8a-c). It is therefore not surprising that K-means clustering of coancestry principal component axes better identified groups of individuals which match previously described subspecies designations and phylogenetic structure (Figure 5, CoA).

### 3.4. Phylogenetic reticulation in the western Napo lineage

Separate RADpainter and PhyloNet analysis of the populations in the western Napo area of endemism and their putative progenitor lineages support the hypothesis that this lineage has a complex history of introgression. One-tailed tests of mean coancestry indicated the null hypothesis that the difference in means among groups is less than or equal to zero can be strongly rejected (ghlt p < 2e-16), and that individuals from the western Napo population have greater median/mean coancestry with each progenitor lineage than their progenitor lineages share with each other (PHvGS/progenitors = ∼1.078, PHvInambari/progenitors = ∼1.16). These analyses establish the asymmetry in admixture patterns detected by STRUCTURE. On average, western Napo individuals share ∼ 7.5% greater coancestry with Inambari individuals (southern Amazon) than they do with other Northern Amazon individuals (glht p < 2e-16), indicating these sampled individuals probably do not include F1 hybrids. We did not observe obvious patterns of clinal variation within our samples of this lineage (spanning ∼20,500 km^2^).

fineRADstructure analysis of the coancestry matrix, as well as visualization of the coancestry matrix indicated that the western Napo population may be differentiated, as well as intermediate when compared to its progenitors (Figure 6a-d). Principle components analysis (Figure 6d) of the coancestry matrix indicate that the western Napo lineage is intermediate on PC1 (compared to progenitor lineages), but differentiated on PC2, which would be expected if the Napo lineage has had sufficient time in reproductive isolation for sorting of ancestral alleles or for the evolution of new alleles (Barrera-Guzmán et al., 2018). This hypothesis is supported by the observation that this lineage has distinct mitochondrial haplotypes derived from Inambari populations (∼separated from Inambari haplotypes by ∼6.5 substitutions or 0.63%), as well as analyses of IBD (below).

Bootstrapped estimates of F_st_ indicated the same general patterns: 95%CI; western Napo vs Eastern Napo (Northern progenitor, 0.07 - 0.090), western Napo v Inambari (Southern progenitor, 0.058 - 0.077): Eastern Napo vs Inambari: (between progenitors, 0.122 - 0.144). These patterns of F_st_ estimates were robust to alternative assignments of geographically proximate individuals to progenitor linages (for example, including an additional adjacent population for both of the putative progenitor lineages). Thus, the introgressed western Napo lineage is less differentiated from *each* of its progenitor lineages than its progenitor lineages are from each other and it is *also* biased in its genetic composition toward Inambari populations.

PhyloNet detected that an evolutionary reticulation is the best model to explain the origin of the western Napo lineage (Figure 6a). This result provides a plausible evolutionary scenario upon which we can interpret other statistics. All parameters in the RJMCMC search achieved an effective sample size > 150 (most >1000), and PhyloNet recovered two network topologies representing 98.6% of the posterior distribution. The top ranked network (*pp* = 57%, Figure 6a) implies the MRCA of the northern Amazonian (Eastern Napo + Jaú) and southern (Inambari + Western Napo) lineages diverged first (clade F). This initial split was followed by a split within the southern lineage that led to the Inambari (S1) and progenitor western Napo (S2a) populations (MRCA of S1 and S2a: Figure 6a). At some point, genetically distinct nuclear genomes from the Northern lineage (clade D/D2) introgressed into S2a (ancestral western Napo), which gave rise to the contemporary western Napo population (S2).

The second-best network (*pp* = 41%, Supplemental Data) depicts a scenario of classic hybrid speciation. First, northern and southern lineages of Clade F diverged, as in the top ranked network. Then, members of S1 and the NW Amazonian lineages merged to form a hybrid lineage. We base our subsequent discussion on the top ranked network, as this hypothesis is more consistent with patterns we detect in other analyses. Support for these two reticulation scenarios is similar, however, and they present interesting alternatives which both require a complex history of reticulation.

Taken in the context of STRUCTURE (Figure 5) and fineRADstructure (Figure 6) results, the gene flow implied by the top ranked network seems to have been asymmetrical, leaving very little evidence of southern admixture among northern lineages (see Figure 5, “weakly resolved eastern Napo,” which are detected to have very limited (< ∼5%) southern admixture). This observation, taken alone, could have been produced by sampling biases. However, demographic parameters estimated with G-PhoCS also support this hypothesis: the only significant migration rate (lower 95% CI > 1 migrant per generation) was inferred from the eastern Napo (member of Guiana Shield lineage) lineage into S2/S2a (Figure 6, Supplementary Table 5).

G-PhoCS also allows us to place a preliminary maximum age constraint on the timing of introgression into S2a. Assuming 2.47 years per generation in *Pseudopipra* (estimated in Bird et al., 2020, though we consider this a lower bound) and a constant mutation rate, the absolute splitting time between S2a and S1 may have been ∼1.2 Ma (Figure 6, Supplementary Table 5). Though we emphasize that this is a rough approximation, this result is loosely consistent with prior divergence time estimates for several groups in the genus based on molecular clock analysis of mtDNA (Castro-Astor, 2014); that study estimated ssp *pipra* (Clade D; Northern Amazon + Guiana Shield lineage) to have branched off from its closest relatives ∼ 0.67 Ma [95% HPD: 0.3506-1.1287]. Therefore, introgression into S2a must have occurred more recently.

Lastly, patterns of admixture (Figure 5, Supplemental Figure 6, 7) suggest some evidence of limited introgression of Northern Amazon alleles into the southwestern Amazonian Inambari population (Figure 5). This could be a consequence of limited gene flow into the Inambari population concurrently with or after the primary introgression event(s) which generated the introgressed western Napo lineage. While our interpretations are conditional on whether or not our genetic sample is indicative of a broader pattern, the product of this process appears to be a distinct, geographically isolated and introgressed lineage within the western Napo area of endemism.

### 3.5. Spatial distribution of genetic variation

Under isolation by distance (IBD), the empirical relationship between geographic distance and genetic distance is related to offspring dispersal distance and population size (Rousset, 1997). Isolation by distance effects are thus predicted when individual dispersal distances are smaller than a species’ range (Teske et al., 2018). Fifteen out of eighteen considered areas (see Figure 4, and the Supplemental Appendix for a description of how these areas were defined) contained > 2 individuals, and thus could be evaluated for isolation by distance effects within them. Five of these regions were identified to have a significant signal of IBD within them at the 0.001 alpha level (Table 2). This signal, however, was not consistent across different proxies of genetic and geographic distance, suggesting that different processes may be contributing to spatial correlations within these areas. Partial Mantel tests of barriers to gene flow indicated that all but one evaluated physical barrier may play significant roles in structuring genetic variation between areas (Table 2). Taken together, Mantel and partial Mantel tests suggest individuals are generally able to disperse within localized regions (with some signal of spatial autocorrelation within regions) but are unable to cross prominent geographic barriers delimiting these areas (Table 2). Notably, the Negro river was the only evaluated barrier which is not detected to exhibit a significant effect— (Table 2 and Supplementary Table 2).

**Table 2.** Summary of Mantel tests. For each geographic barrier and sampling region we were able to evaluate, we report results for analyses using phylogenetic patristic distance and coancestry as proxies for individual genetic distances. Analyses for both linear and log geographic scales are shown. Asterisks indicate *p* values less than 0.001, which we interpret as statistical significance (see discussion in text).

| **Approximate Barrier** | **Patristic distance** |  | **Coancestry** |  |
| --- | --- | --- | --- | --- |
|  | p < 0.001 | p < 0.001 | p < 0.001 | p < 0.001 |
| Cordillera de Talamanca | - | * | - | - |
| Andes (1) | - | * | - | * |
| Andes (2) | - | * | - | * |
| Andes (3) | * | * | - | * |
| Ucayali river | * | * | * | * |
| Eastern Marañón + Hauallaga rivers | * | * | * | * |
| Putumayo river | * | * | * | * |
| Purus river | * | * | - | * |
| Madeira river | * | * | * | - |
| Tapajós river | - | * | - | - |
| Xingu river | * | * | - | - |
| Cerrado (1) | * | * | - | * |
| Cerrado (2) | * | * | * | * |
| Japurá river | - | * | * | * |
| Negro river | - | - | - | - |
| Essequibo river | * | * | * | * |
| Amazonas river | * | * | * | * |
| **Regions** |  |  |  |  |
| Full dataset | * | * | * | * |
| Central America - Costa Rica (Clade A3) | - | - | - | - |
| Central America - Panama (Clade A4) | - | - | - | - |
| North Andean – San Martín (Clade A1) | ? | ? | ? | ? |
| South Andean Peru (North) (Clade B2) | - | - | - | - |
| South Andean Peru (South) (Clade B1) | - | - | - | - |
| weakly resolved Guiana Shield (Clade D) | - | - | - | * |
| Unresolved Jaú | - | * | - | * |
| Weakly resolved eastern Napo (Clade D2) | - | - | - | - |
| Weakly resolved Suriname + Amapá (D1) | - | - | - | * |
| Western Napo introgressed lineage (Clade E) | - | - | - | - |
| Western Inambari endemic (Clade C1) | * | * | - | * |
| Eastern Inambari endemic (Clade C3) | - | - | - | - |
| Rondonia endemic (Clade C4) | ? | ? | ? | ? |
| Tapajós endemic (Clade C5) | - | - | - | * |
| Xingu endemic (Clade C6) | - | - | - | - |
| Atlantic Forest – Bahia (Clade C8) | - | - | - | - |
| Atlantic Forest – Espírito Santo (Clade C9) | ? | ? | ? | ? |
| Atlantic Forest – Rio (Clade C10) | * | - | * | * |
|  | **linear** | **log** | **linear** | **log** |

Model based analysis of migration rates with EEMS largely corroborated the specific hypotheses implied by Mantel tests. For dissimilarities between pairs of demes, the correlation between observed and fitted values was high – R^2^ = 0.871 – within demes somewhat less – R^2^ = 0.579 (Supplementary Figures 10, 11). Thus, the EEMS model appears to do a good job of describing spatially structured variation in this dataset, and a non-stationary migration surface is a better fit to the data than simple isolation by distance. The EEMS visualization detects many strong signals (posterior probability > 0.9) of barriers to gene flow (Figure 7). As expected, the Andes and the Amazon river were unambiguously detected as the most significant barriers to gene flow and are clearly recovered as landscape features, even though the model does not know these features exist. EEMS also performs well in inferring the spatial extents of the lowland Amazonian range (range map outlined in black), despite very limited genetic sampling toward the edge of the distribution.

**Figure 7.**
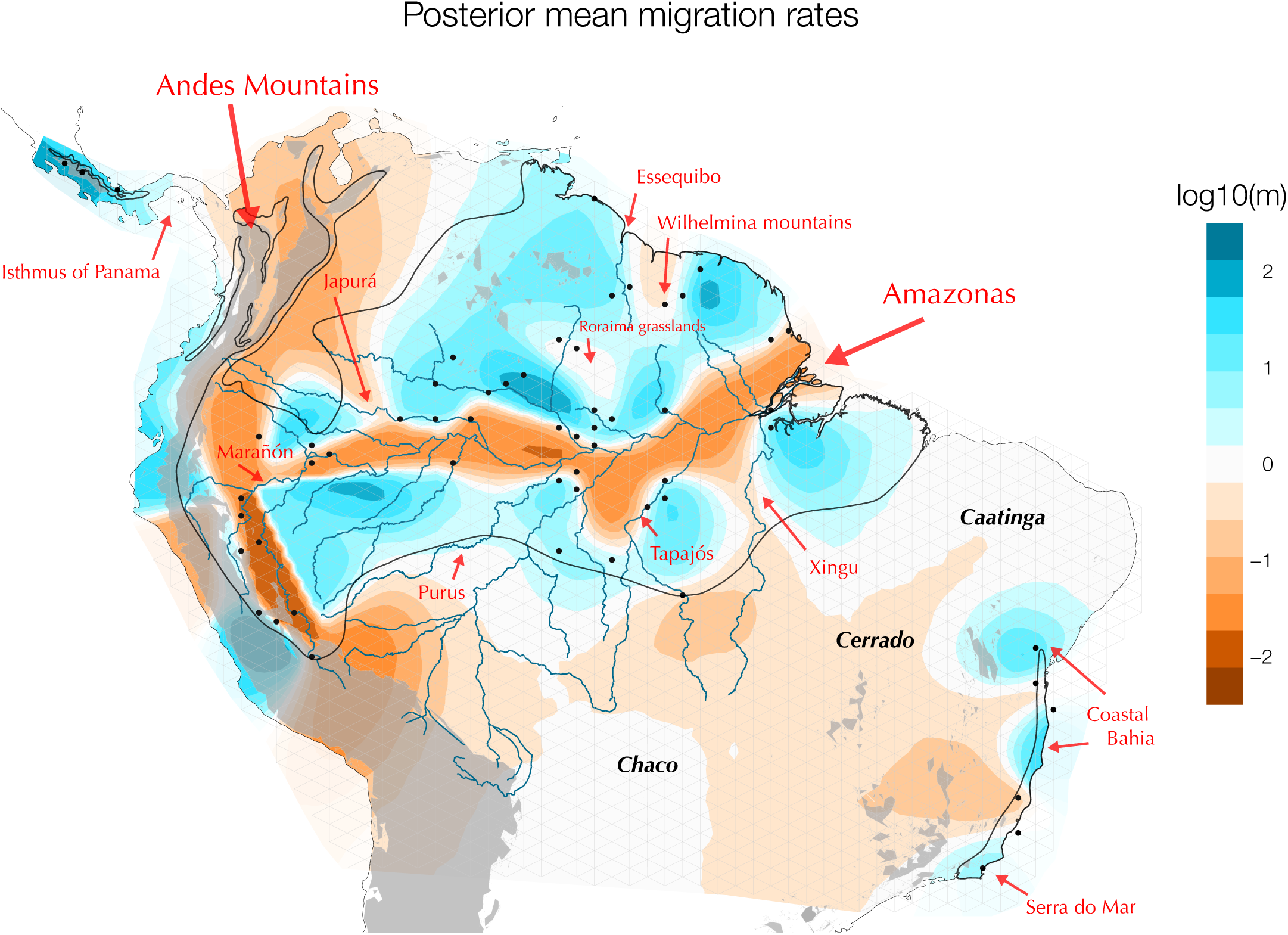
Estimated Effective migration surface (EEMS). The migration surface links the topography and drainage system of the Neotropics to spatial patterns of genetic variation. Shown are results estimated from a model with 2000 demes (vertices). Bluer colors indicate areas where gene flow is more likely to be occurring (i.e., more connectivity than the average signal of isolation by distance), while browner colors indicate areas where barriers to gene flow are probable. The Andes and the Amazon River are the most clearly inferred barriers to gene flow. Major rivers and other barriers, as well as a > 1000m contour (light grey) are overlaid to aid interpretation. See text for additional discussion. Note: in EEMs, input sample coordinates are associated with the closest vertex (i.e., depicted sample markers do not indicate exact sample position). See the Supplementary Data for a high-resolution version of this figure.

EEMS further indicates signals of deviation from constant IBD that correspond to several Amazonian tributaries (Figure 7), which are also recapitulated by Mantel tests (Table 2). As noted above, the Negro river is not clearly identified by EEMS, nor by Mantel tests as a significant barrier in *Pseudopipra*. Simultaneously, uneven sampling, and the predominant signal of the Amazon river in EEMS analysis at this broad scale, may be obscuring more localized effects in the Amazon basin which are detected with Mantel tests; for instance, across the Madeira river (Table 2, *p* < 0.001). An additional strong barrier to gene flow was inferred in the south-central amazon (south-central brown patch in Figure 7), corresponding to the Caatinga - Cerrado – Chaco “dry-diagonal” biomes. Strong signals of genetic connectivity are inferred in many localities (blue), notably west of the Andes, connecting north Andean and Central American lineages (despite a lack of sampling in that region). This link perhaps reflects the common ancestry among montane lineages in these regions. Consistent with other analyses, EEMS detects the large Guianan + Northern Amazonian region (E. Napo + Jaú + Guiana) to have relatively weak population structure across a variety of barriers (including the Negro river). EEMS detects several areas of continuous gene flow within the Atlantic Forest, likely corresponding to the Coastal Bahia and Serra do Mar areas identified by da Silva et al. (2004) (Figure 7). The introgressed hybrid lineage from the western Napo area is inferred to be genetically isolated from neighboring populations, with reductions in gene flow which appear to coincide with the Marañón river in the south and Putumayo/Iça rivers to the north.

Lastly, EEMS detected two broad clusters of relatively high genetic diversity (relative heterozygosity: Supplementary Figure 12). One is centered along the Amazon river and was relatively uniform within the northern Amazonian basin, reflecting relatively high genetic diversity in the Guiana Shield populations. Notably, the other prominent high diversity area overlaps with our sampling of the introgressed western Napo lineage. Areas of relatively low genetic diversity included the Atlantic Forest, the Peruvian Andes, and Central American lineages. These patterns are consistent with direct estimates of regional variation in allelic richness (Supplementary Appendix).

### 3.6. Vocal variation

We examined 198 recordings of *Pseudopipra* and identified 14 distinct vocalizations that can be diagnosed by ear or visual inspection of sonograms (arbitrarily labeled numerically in Figure 8). This set of vocalization types includes both advertisement vocalizations from males at leks and calls by males and females in other contexts. After examining recordist notes, we generated a preliminary hypothesis which distinguishes eleven highly variable and distinct, lekking advertisements (Figure 8a) from three broadly shared call types (Figure 8b). Our delimitation of vocalizations into advertisement and call categories partly reflects expert opinion, as the behavioral context of vocal records was not always clear (see Supplemental Appendix). Work by Castro-Astor et al. (2007) provides important insights of the vocal behavior of Atlantic forest populations, but most *Pseudopipra* populations lack similarly detailed studies. Vocalization types within each category can be statistically discriminated based on a combination of the frequency, duration, and number of notes. PCA and logistic regression on lekking vocalizations (n=114, Supplementary Table 3), and calls (n=33) found significant differences among types (*p <* 0.001). The first three axes of a PCA explained ∼90% of the variation in lekking vocalization characters, with PC1 (∼64%) primarily explaining variation in note number and frequency (Supplementary Figure 15).

**Figure 8.**
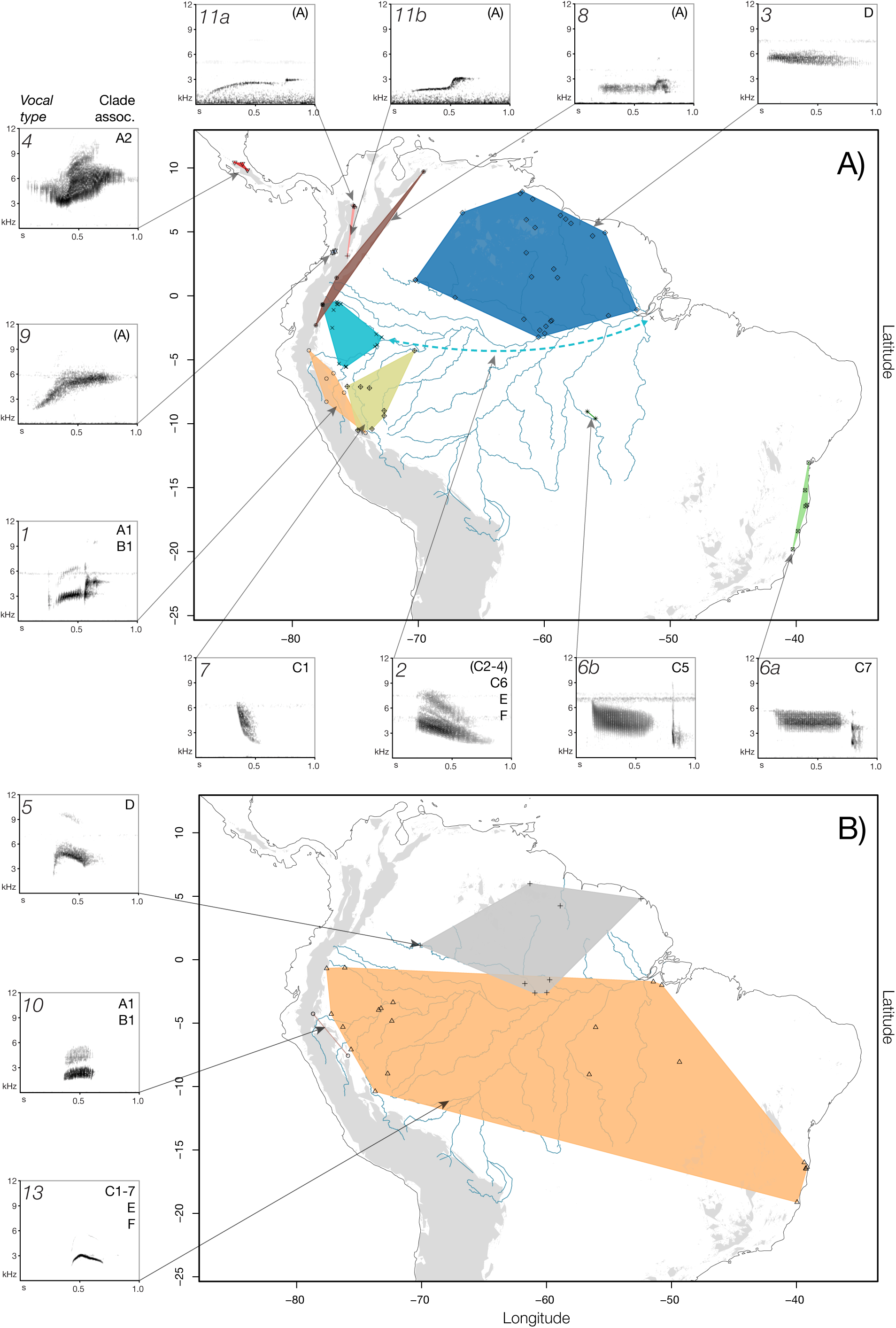
Panel A: Summary of lek vocalization phenotypes. Our analysis identified 11 qualitatively distinct lek vocalization types that can be diagnosed by ear, or by visual inspection of sonograms (arbitrarily labeled numerically). See Supplementary Figure 13 for vocalization measures. The inset map indicates the minimum implied ranges for each identified vocalization type, with colors and plotting symbols matching those in PCA plots (Supplemental Figures 14, 15). The pale blue dashed line indicates where the “southern Amazon” vocalization type 2 may be shared, apparently plesiomorphically (Supplementary Figure 16). Vocalization type 2 may therefore occur in other unsampled areas of the southern Amazon. Type 1 also appears to be shared pleisiomorphically between populations on the eastern slopes of the Andes (Clade A1) and in populations from the southern Cordillera Azul in Loreto (Clade B1). Panel B: Summary of call vocalization phenotypes. Our analysis identified three distinct but similar call vocalization types that can be diagnosed by ear, or by visual inspection of sonograms. Vocalization type 5 was primarily recorded in the northern Amazon basin, coinciding with the broad ranging Northern Amazon + Guiana Shield lineage (Clade D). From our initial classification, one recording from the Atlantic Forest (XC427315) was assigned to type 5, suggesting this type may be shared between the Guiana Shield clade and the Atlantic Forest clade (not shown). As with lek vocalization type 1, call vocalization type 10 was shared pleisiomorphically across Andean clades (A1 + B1). Call vocalization type 13 was entirely restricted to but shared across diverse lowland southern Amazonian forms (Clade C). Letters to the top right of sonograms indicate explicit [e.g., *A*] or implicit clade associations [e.g., (*A*)]. See the Supplemental Appendix for additional details and discussion.

Our assignments of vocal types to well supported genetic groups are preliminary in the absence of recordings from the same individuals from which we have genetic samples. Nevertheless, we found that most of the 11 lek advertisement vocalizations are phylogenetically diagnostic of the populations in which they were recorded (except for two cases of apparent plesiomorphy, Supplementary Figure 16b). In contrast, the three call vocalization types we identified were diagnostic of one lineage (type 5, clade D), or shared plesiomorphically across two ancient clades (type 10 and 13, Supplementary Figure 16a). See Taxonomic Summary, below.

## 4. Discussion

In comparison to the great diversity of Neotropical organisms, there have been few studies focused on understanding both fine and broad scale diversity patterns within broadly distributed Neotropical taxa, and this has slowed the biogeographic interpretation of the effects that many landscape features have on the evolution of Neotropical organisms (Fernandes et al., 2013; Harvey et al., 2017; Harvey and Brumfield, 2015; Marks et al., 2002; Milá et al., 2012; Nyári, 2007). We investigated the phylogenetic and population genetic history of a continentally distributed taxon currently recognized as a single species– *Pseudopipra pipra*– using a genomic dataset which samples thousands of loci across the genomes of 232 individuals representing eight of thirteen *Pseudopipra* subspecies. Two additional subspecies, *pygmaea* and *coracina*, were represented by mtDNA. Our genetic sampling encompasses a majority of the broad range of *P. pipra*, spanning Neotropical forests from Central America to the Atlantic Forest of Brazil, including highland and lowland areas of endemism. Importantly, our sampling of vocal variation fills in gaps in our genetic sampling—therefore, our intersecting datasets allow us to assess at least some data from all thirteen previously identified subspecies (Figure 2).

The predominant biogeographic signal we detect in our dataset is of two successively nested montane clades as the sister groups to several major lowland Amazonian and Atlantic Forest clades (Figure 5, see *Historical Biogeography*, below). This pattern implies that lowland South American and montane Central American populations of *Pseudopipra* expanded historically “out of the Andes.” Ancient lineages endemic to the Andes may have originated around the onset of the Pleistocene ∼2.5 Ma (molecular clock estimates from Castro-Astor, 2014). The lowland clade within *Pseudopipra* originated (∼1.085 [0.57-1.79] Ma) in southwestern Amazonia (Figure 4) and expanded, occupying Amazonia on both margins of the Amazon river, and eventually reached the Atlantic forest (see below). Therefore, our study provides an unusual example of a widespread suboscine passerine which originated in the Andean highlands.

### 4.1 Population genetic variation

We examined our dataset using a variety of population genetic approaches intended to assess the number of natural genetic groupings that may be present in *Pseudopipra*. Phenetic K-means clustering of SNP data and analysis with STRUCTURE estimated a minimum of 5 genetic clusters with broad geographic ranges (Figure 5, Supplementary Figure 2, Supplementary Figure 4). K-means clustering of the coancestry matrix (which captured substantially more variation than standard PCoA) more finely partitioned the dataset into 8 genetic clusters (Figure 5 CoA, Supplementary Figure 8), which closely recapitulates previously identified subspecies boundaries (Figure 2). Phylogenetic (RAxML, SVDquartets) and model-based population genetic (fineRADstructure) analyses of nuclear genomic data detected fifteen identical groups of individuals which coincided with sampling regions delimited by clear geographic barriers (results, Figure 3, 4, Supplementary Appendix). Two additional subspecies represented by mtDNA (results) bring the total number of distinct genetic populations delimited by geographic barriers to at least seventeen. The close concordance observed between population structure and landscape features implies that the evolutionary history of *Pseudopipra* within the Neotropics is deeply connected to the South American landscape, adding support to a rich body of literature endorsing this hypothesis (Brumfield, 2012; Cracraft and Prum, 1988; Silva et al., 2019).

Our sampling reveals a “horseshoe” pattern of genetic differentiation across the Amazon basin (Figure 4), with no evidence of gene flow across the Amazon river, except in the western headwaters where the Amazon is narrowest and floodplains are dynamic and young (Haffer, 1992; Pupim et al., 2019; Räsänen et al., 1987). Given that the genus originated in the west, we hypothesize that northern and southern clades diverged across the Amazon river in this region. For virtually all evaluated cases, we find significant effects of geographic barriers on subdividing genetic variation within this species complex, including across the Amazon river and most of its larger tributaries, with the notable exception of the Negro river (Figure 7 and Supplementary Table 2a, 2b). As expected, the “dry-diagonal” Caatinga, Cerrado, and Chaco belt is a strong barrier to gene flow, isolating Atlantic Forest lineages from their southeastern Amazonian Xingu relatives— as are the Andes, which exhibit a disproportionate effect on divergence between Peruvian foothills populations and Central American lineages (see Supplementary Appendix for additional discussion).

Multiple analyses indicate that the western Napo population has a complex history of initial and early differentiation from the broader Southern Amazonian clade, followed by subsequent, asymmetric genetic introgression from Jaú and eastern Napo populations of the Guianan Shield + Northern Amazon clade (Clade D; see discussion below). When the genetic data are analyzed under either concatenated or coalescent phylogenetic frameworks which do not allow for phylogenetic reticulation, the western Napo population is reconstructed as most closely related to Guiana Shield + Northern Amazon populations (Figure 3, Figure 5, Clade D). In contrast, phylogenetic network analysis indicates that this population is derived from individuals of Inambari (southern Amazon) provenance (Figure 6, also summarized in Figure 4). This incongruence between methods is likely due to the complexity of the constituent gene trees in this population-level differentiation induced by asymmetric introgression from the Jaú and eastern Napo populations (Thom et al., 2018). We hypothesize that introgression has introduced more recently derived variation from the Guiana Shield + Northern Amazon clade D into an older, differentiated, population of the Southern Amazonian clade, pulling the phylogenetic affinity toward the northern clade. A similar scenario may have occurred in another manakin lineage, *Lepidothrix coronata*, in which nuclear and mitochondrial markers seem to show contrasting topologies regarding the western Amazonian Napo lineage (Reis et al., 2020).

### 4.2 History of an introgressed hybrid lineage

The substantial level of observed introgression (>40%) in the western Napo individuals, as well as a suite of genomic and phenotypic evidence, suggests that this genetic population may be best recognized as a hybrid lineage, though it is uncertain if it can be considered a product of hybrid speciation. At the time of writing, five prior examples of hybrid species formation in birds have been evaluated with genomic data: (1) the Golden-crowned manakin *Lepidothrix vilasboasi* (Barrera-Guzmán et al., 2018), (2), the Audubon’s warbler *Setophaga auduboni* (Brelsford et al., 2011), (3) the Italian sparrow *Passer italiae* (Hermansen et al., 2011), (4) the Hawaiian duck *Anas wyvilliana* (Lavretsky et al., 2015), and (5) a putative species of Darwin’s finches *Geospiza* (Lamichhaney et al., 2018). These examples of mosaic genome hybrid speciation, in which the fusion of two or more progenitor lineages leads to a reproductively isolated hybrid species (Counterman, 2016; Jiggins et al., 2008), repeatedly demonstrate that modern patterns of avian diversity may have been significantly influenced by historical phylogenetic reticulation (Suh, 2016). Our analyses of genomic data across *Pseudopipra* indicate that the western Napo lineage may be a differentiated, allopatric phylogenetic species which has a genomic composition derived from heavy historical admixture with a non-sister lineage. This historical scenario is not the same as those examples cited above, but it has produced a population genetic signature which is quite similar (compare to Barrera-Guzmán et al., 2018). These observations force us to ask whether historical process, or contemporary genetic pattern should be paramount in delineating a hybrid species.

Some authors have proposed strict population genetic criteria for identifying homoploid hybrid speciation, including demonstrating (1) evidence of reproductive isolation from parental species, (2) evidence of past hybridization, and (3) evidence that intrinsic reproductive isolation was derived from the hybridization event (Schumer et al., 2014). The first two conditions seem to be satisfied by the western Napo *Pseudopipra* lineage. First, phylogenetic and population genetic analyses are consistent with contemporary reproductive isolation. Phylogenetic analyses of nuclear genome data, including concatenated and coalescent analyses (Figure 3, 4, 6), recover this lineage as a well-supported reciprocally monophyletic group. Differentiated mtDNA haplotypes from a broader geographic sample (Figure 6) also form subgroups derived from southern Inambari haplotypes (results, Supplementary Figure 17), implying reproductive isolation from female northern progenitors.

PCA-based and model-based population genetic cluster analyses (Figure 6b, 6c, Supplementary Figures 4-9), help establish this lineage as intermediate *and* differentiated when compared to the living descendants of its inferred progenitors. The level of differentiation on PC2 corresponds to about 10% of the total variation explained by PC1 and PC2 (∼85%). At the same time, Mantel tests, as well as model-based analysis of isolation by distance, support the hypothesis that contemporary gene flow into and out of this population is restricted (Figure 7, Supplementary Figure 12). Thus, we interpret our G-PhoCS results of a significant migration parameter (results) as indicative of a window of historical introgression, which could be further explored with simulation studies. Estimates of inbreeding coefficients also support this hypothesis and suggest similar levels of inbreeding among *Pseudopipra* populations in the western Amazon (F_IS_ > 0.1, Supplementary Figure 9, and Supplementary Appendix). The western Napo population is thus not detected to be more outbred than most populations (i.e., recently introgressed). In sum, these patterns observed in our genetic data are consistent with the hypothesis of contemporary reproductive isolation for the introgressed Western Napo populations.

The second criterion from Schumer et al. (2014), evidence of past hybridization, is directly supported by multiple model-based analysis of genomic admixture and phylogenetic reticulation (Figure 5, 6), and indirectly supported by model based and phenetic cluster analysis (Supplementary Figures 4-9). We hypothesize that hybridization initially occurred between progenitor lineage S2a and western lineages of the Guiana Shield + Northern Amazon clade (Figure 6a). Lineage S2a was then replaced by the novel introgressed lineage, which exchanged genes with neighboring populations within a finite time window.

The third criterion from Schumer et al. (2014), evidence that intrinsic reproductive isolation was derived from the hybridization event, is recognized as the most difficult to demonstrate, and probably the most tenuous. As discussed by Nieto Feliner et al. (2017) in critique of Schumer et al. (2014), this criterion may be useful for framing the strength of evidence for or against the existence of a homoploid hybrid species, but only if one adheres strictly to the biological species concept (Mayr, 1942). If one applies a more evolutionary definition of species, like the metapopulation lineage concept, in which the only necessary requirement is demonstration of separately evolving lineages (de Queiroz, 2005), the importance of demonstrating intrinsic reproductive isolation to claim species status for a putative hybrid lineage (or any lineage at all) is less clear to us. Speciation in birds in particular is thought to proceed relatively fast, and may be more affected by external factors than internal ones (Grant and Grant, 1997). Further, requiring hybridization to be the *source* of intrinsic isolation excludes the possibility that extrinsic or pre-zygotic isolation may evolve from changes that enable hybrids access to new niches, which may also be geographically isolated relative to parental forms (Nieto Feliner et al., 2017).

Intriguingly, the evolutionary scenario implied by phylogenetic reticulation analysis for the introgressed western Napo hybrids provides a mechanism to explain a confusing distribution of vocalization types (Figure 8, Supplementary Appendix). The males from the Napo area of endemism share vocalization type 2 with birds in the southeastern Amazon (Figure 8, Supplementary Figure 16), and with the restricted Huallaga valley *pygmaea* subspecies (the sister group to all lowland Amazonian forms, according to mtDNA). Manakins are suboscine passerines and have innate vocalizations (though see Saranathan et al., 2007). Therefore, the introgressed Napo lineage appears to have inherited its vocalizations from the MRCA of groups S1 and S2 (Figure 6a) and retained it through subsequent introgression with northern lineages that have a different vocal type (Type 3 and/or 5). In this scenario, vocal type 7 in the western Inambari represents more recently derived variation in the southern Amazonian clade (Figure 8, Supplementary Figure 16).

As noted earlier, the western Napo lineage also has mtDNA haplotypes derived from Inambari populations (Supplementary Figure 17). In addition to implying reproductive isolation from the Guiana Shield clade, this mitochondrial pattern implies that the introgressed Napo lineage may be descended from the union of progenitor female S2a and progenitor western males of the Guiana Shield + Northern Amazon clade, perhaps through male biased dispersal or mate choice (a preliminary test of male biased dispersal did not detect a signal, see the Supplemental Appendix). Regardless, this pattern raises fascinating questions about why and how the western Napo lineage may have retained its southern vocalization type.

Taking all of the available genetic and phenotypic evidence into account, we suggest that the western Napo lineage may be an additional example of the formation of a hybrid lineage in birds, and one that was produced via an underappreciated historical mechanism. Assuming that our limited sampling is representative of the *discolor* subspecies, the patterns we observe appear to be most similar to “hybrid trait speciation,” in which a hybrid species is formed after introgression from one species into a genomic background of a close relative (Counterman, 2016; Jiggins et al., 2008; Marques et al., 2019). This type of combinatorial hybrid speciation was originally proposed from studies of *Heliconius* butterflies (Jiggins et al., 2008; Salazar et al., 2010).

### 4.3 Comparative and Historical Biogeography in South American Lowlands

The spatial pattern of population structure and evolutionary relationships we identify here is broadly congruent with other recent studies of Neotropical biogeography in upland *terra firme* forest birds, contributing another example of the substantial phylogeographic pseudocongruence across taxa at this wide scale (e.g. Cunningham and Collins, 1994; Harvey et al., 2017; Harvey and Brumfield, 2015; Ribas et al., 2012; Silva et al., 2019; Thom and Aleixo, 2015). Southern Amazonian tributaries, such as the Xingu, Tapajós, and Purus, delimit recently isolated populations, while northern *Pseudopipra* populations show weaker evidence of being isolated by tributaries with larger discharges, like the Negro and Japurá rivers. The widespread occurrence of white sands habitats, which are used by *Pseudopipra* along the Negro river basin (Borges et al., 2001) may weaken the effect of this river barrier, as has been observed in white sand vegetation associated species, including the piprid *Xenopipo artronitens* (Capurucho et al., 2013).

A recent comparative analysis shows that large rivers delimit genetic clusters in as many as 23 groups of upland Amazonian forest birds, strongly arguing in favor of these rivers acting as barriers to gene flow (Silva et al., 2019). Our study thus adds *Pseudopipra* to a list of similarly distributed Neotropical bird species recognized to have spatial patterns of genetic diversity influenced by Amazonian rivers. We speculate that the somewhat distinct pattern found for *Pseudopipra* may be related to its recent expansion throughout the eastern Andean lowlands and its habitat tolerance.

Within the manakins, Ohlson et al. (2013) placed *Pseudopipra* as sister to the genus *Ceratopipra*, which includes five well-recognized species that are extensively codistribtuted with *Pseudopipra.* The montane origin of *Pseudopipra* may be congruent with the observation that *Pseudopipra* is the sister group to *Ceratopipra,* which is also broadly distributed in the Neotropical lowlands from Central America to the Atlantic Forest and has a montane sister group (*C. cornuta*) (Leite et al., 2021; Ohlson et al., 2013; Prum, 1992). The breakpoints among these *Ceratopipra* species are highly concordant with the breakpoints among the genetic clusters within the *Pseudopipra* complex that we have presented here, implying that these closely related taxa have components of their phylogeographic history in common. See the Supplemental Appendix for additional notes, as well as Castro-Astor (2014). Future studies of these patterns should be fruitful.

A key feature of the phylogeny of *Pseudopipra* is that montane Andean lineages are sister groups to both the montane Central American and the lowland Amazonian-southeast Brazilian lineages (Figure 5). Ancestral character reconstruction under Bayesian stochastic mapping and maximum parsimony therefore unambiguously reconstructs the most recent common ancestor of *Pseudopipra* as Andean (Figure 3-5, Supplemental Appendix). Thus, it appears that the most recent common ancestor of *Pseudopipra* was an Andean lineage restricted to subtropical, lower montane forest. The earliest diversification event in the genus was likely the differentiation of the ancestral Andean populations across the region where the headwaters of the Huallaga river, Peru are currently located. The northern lineage gave rise to the subtropical montane lineages of the northern Andes and Central America (Clade A). The southern lineage gave rise to a subtropical montane Southern Peruvian lineage (Clade B) and a southwestern lowland Amazonian lineage (*pygmaea*, represented in our dataset by mtDNA). Subsequently, the Amazonian lineage expanded to the eastern lowlands and differentiated into the northern lowland Amazonian + Guiana Shield (Clade D in Figure 3) and southern lowland Amazonian/Atlantic Forest (Clade C in Figure 3) clades, perhaps tracking available upland forest habitats at a time when western Amazonia was dominated by wetlands (Pupim et al., 2019). Within the Southern Amazonian clade, differentiation proceeded from west to east (Figure 3, 4). Contrastingly, the northern Amazonian lineage also expanded all the way to the Atlantic Ocean, but without producing strong phylogenetic structure (Figures 3, 4).

As upland forest became available in western Amazonia, replacing the seasonally flooded terraces of the Solimões river (Pupim et al., 2019), the area was probably occupied by neighboring upland lineages. At the same time, the main river channels became entrenched, promoting contact and isolation among western lowland populations (Bicudo et al., 2019). Both the Solimóes and Japurá rivers seem to have undergone recent changes due to sedimentation dynamics and channel reorganization (Ruokolainen et al., 2019), and such processes likely affected upland forest bird populations. The pattern of introgression in the western Napo *Pseudopipra* lineage may be temporally congruent with a similar pattern described for *Lepidothrix coronata,* with a split across the upper Solimões around 1-1.5 Ma, and hypothesized introgression from Northeastern lineages (Reis et al., 2020).

The effects of these recent landscape reconfigurations in western Amazonia probably varied according to the habitat affinities and intrinsic characteristics of the different clades, with some retaining Northern and Southern lineages delimited by the main Solimões channel (Ribas et al., 2012; Silva et al., 2019); some in which southern lineages are more closely related to lineages in the Solimões/Japurá interfluve (Silva et al., 2019, counterclockwise pattern); and some in which northern lineages group with lineages at the Solimões/Madeira interfluve (Ribas et al., 2018; Silva et al., 2019). These varying patterns point to a dynamic history of western Amazonian upland forests, and introgression with the formation of hybrid lineages may have been common in the history of many taxa. As genomic data becomes available for more clades, hybrid lineages may turn out to be common in this region of high diversity and extremely dynamic landscape history.

Lastly, a well-supported distinct montane (>1000m) (BS: 100) (Clade C2 in Figure 3; Cushabatay area, northern Cordillera Azul, Loreto Prov., Peru) is recovered as nested within the well supported lowland western Inambari subclade (Clade C1 in Figure 3). Thus, there may have been at least one secondary invasion of montane Andean habitats from a lowland ancestor. Patterns of admixture support this hypothesis – montane individuals from the northern Cordillera Azul and their proximate lowland relatives all show limited genomic admixture from derived lineages of the Guiana Shield + Northern Amazon clade (yellow in Figure 5). This result is consistent with a scenario of secondary invasion of the highlands from lowland Inambari lineages after weak introgression from the western Napo (see below).

### 4.4 North Andean diversity

Our final genomic data set does not include any samples from Andean Ecuador or Colombia, a region that includes four previously recognized subspecies and at least four distinct vocalization types (see Taxonomic Summary). Unfortunately, the three Andean Ecuadorian tissue (representing subspecies *coracina*) specimens in our original sample were not of high enough preservation quality to be viable for ddRAD sequencing (Supplementary Table 1). We were able to obtain mtDNA from these samples however, which unambiguously formed a monophyletic cluster with our single San Martín (North Andean Peru) specimen in mtDNA gene tree analyses (Supplemental Appendix, Supplementary Figure 17). A prior study by Castro-Astor (2014) based on mtDNA also included one Andean Ecuadorian sample and one Central American sample, which unambiguously clustered together (*pp* = 100). Thus, it is likely that our unsampled Andean lineages of Ecuador and Colombia (including the subspecies *minimus, bolivari, unica,* and *coracina*) are members of the northern Andean clade represented in our genomic data set by our single San Martín specimen. As noted earlier, our San Martín specimen had approximately equal probability of assignment to multiple populations, which could indicate heavy admixture, or that this is a single sample from a highly distinct population which has its own distinct history. These patterns may also be consistent with the hypothesis that the North Andean clade is the source of montane and lowland *Pseudopipra* diversity.

### 4.5 Phenotypic and Vocal Evolution

Thirteen subspecies have been previously described based on variations in plumage color and, to a lesser extent, size (See Taxonomic Summary in Supplementary Appendix). Male plumage coloration varies among the subspecies of *Pseudopipra* in the glossiness of the black body feathers, the length of the white crown feathers, and the color– white, gray, or black– of the bases of these crown feathers. Female plumage coloration often shows more striking differentiation among subspecies than does male plumage, including variation in the shade of olive green on the body, the shade and extent of gray on the crown and face, and olive, yellow, or gray coloration on the belly (Chapman et al., 1914; de Schauensee, 1945, 1950; Ridgway, 1906; Wetmore, 1972; Zimmer, 1936).

Although we did not conduct a detailed analysis of plumage variation among populations of *Pseudopipra*, we did observe study skin specimens of all but one (*bolivari*) of the previously described subspecies in museum collections. All of the ten subspecies that were included in our genetic samples were identified as distinct, diagnosable, monophyletic groups. In other words, in all cases that we were able to test, traditional taxonomic practices conducted between 1758 through 1936 successfully identified and named distinct evolutionary lineages within *Pseudopipra*. We did not have genetic samples of the northern Andean taxa *minima, unica,* and *bolivari*, but we note that they all share entirely white crown feather bases (forecrown feathers only in *bolivari*)(de Schauensee, 1945, 1950), implying that this unique plumage character state is derived within *Pseudopipra*, and that *minima*, *unica*, and *bolivari* form a clade. One lineage containing two subspecies– *separabilis* and *cephaleucos-* has evolved another unique, shared, derived plumage character– a distinctive, second, predefinitive male plumage which has also further differentiated between the two subspecies. In *cephaleucos,* predefinitive males have an olive green back, a white crown, and slate gray face and belly. In *separabilis,* predefinitive males are similar with lighter gray belly, and a medium gray, instead of white, crown. In conclusion, plumage coloration appears to provide highly informative evidence of evolutionary lineage status in this genus.

Our analysis of vocal behavior indicates that vocalizations are also highly informative of lineage identity within *Pseudopipra*. The complex phylogenetic history we have documented with genetic data has clearly had a strong impact on the evolution of vocalizations in *Pseudopipra.* Because lek vocalizations are a focus of female choice in manakins and because suboscine passerines have largely innate vocalizations, the extensive vocal and genetic differentiation among populations of *Pseudopipra* strongly indicates the existence of many species within *Pseudopipra* (Remsen, 2005).

We identified fourteen distinct vocalization types (Figure 8), and nearly all were restricted to and diagnostic of a single previously recognized subspecies or a broader monophyletic group (See Expanded Taxonomic Summary in Supplementary Appendix for details). One of the two exceptions was vocalization type 2, which appears to be shared plesiomorphically across the Amazon basin between populations of *pygmaea* in the Huallaga Valley, *discolor* in the western Napo region of Ecuador and Peru, and *separabilis* in Para, Brazil, but not in the intervening southwestern Amazonian populations of *microlopha* (Supplementary Figure 16). Unfortunately, available vocal sampling is very limited in the intervening regions along the south side of the Amazon Basin, which are populated by a nested series of genetically distinct lineages with successively closer relationships to southern Amazonian *separabilis* and *cephaleucos* from the Atlantic forest of Brazil.

The other exception is the sharing of vocal type 1 (’*trill-dink*’) between populations of *occulta* from the eastern slopes of the Andes from San Martin, Peru to southeastern Ecuador (A1) and populations of *comata* from the southern Cordillera Azul in Loreto (B1). The song of the southern clade (B2) from southern Huánuco (AMNH 820866, 820952), Pasco, Junín, and Cusco is currently unknown. Surprisingly, these geographically adjacent lineages sit on either side of the deepest phylogenetic separation within *Pseudopipra*. The *comata* populations are phylogenetically closer to northern Andean and Central American lineages that exhibit substantial vocal diversity. Likewise, the southern Cordillera Azul populations are more closely related to other south Peruvian and Amazonian populations which also exhibit great vocal diversity. This result implies that the vocalization type 1 was shared by the ancestral lineage and has been retained in these lineages (Supplemental Figure 16). We identified the same pattern for call vocalization type 10.

Our phylogenetic analysis also identified a distinct montane clade from the northern Cordillera Azul (Clade C2; Cushabatay area; Alverson et al. 2001) which is closely related to populations from lowland forests of eastern Peru south of the Marañón river and east to Purus river, Brazil, currently recognized as *P. p. microlopha*. The hypothesis that this lineage is a secondary expansion into montane forests from the nearby Amazonian populations is strongly supported by the evidence that this montane population shares same song type 7 (’*jeer’*) with the lowland populations of *microlopha* (ML-190390, ML-190388, and ML-190262).

### 4.6 Taxonomy and Revised Classification

The present set of analyses demonstrate a number of genetically well-differentiated and phenotypically diagnosable lineages and provides compelling new evidence for evaluating the species status of *Pseudopipra*. Although there are gaps in our sampling, we find that there are at least eight genetically well-differentiated, and phenotypically diagnosable lineages of *Pseudopipra*, and this is reflected in our proposed taxonomic revision below. Nonetheless, it should be noted that there is more genetic variation than observed vocal variation implies, and more vocal variation beyond our geographic coverage of genetic samples (especially in northern Andes). Thus, our taxonomic recommendations for *Pseudopipra* almost certainly represent an underestimate of extant diversity, underscoring the need for continued reassessment of species limits and of diversity patterns in the Neotropics.

Our analysis provides the first comprehensive opportunity to reevaluate species limits within *Pseudopipra* based on phylogenetic, population genetic, vocal, and plumage differentiation among populations and named subspecies (See the Expanded Taxonomic Summary in the Supplementary Appendix). This proposed classification is a conservative treatment that recognizes the limitations on our genetic sampling of populations in the northern Andean clade from Ecuador and Colombia. Four of these unsampled (and under-sampled) northern Andean subspecies – *coracina*, *minima*, *bolivari*, and *unica* – have unique and highly differentiated vocalization types, and diagnosable plumage differences. Our phylogenetic results strongly suggest that most of the *Pseudopipra* subspecies are distinct evolutionary lineages deserving species status under either the phylogenetic or biological species concepts. In sum, *Pseudopipra* may include 15-17 distinct species that have rapidly arisen in the last ∼2.5 Ma, but further reclassification will require additional sampling of genetic, vocal, and plumage variation.

## 5. Taxonomic Summary

*Pseudopipra coracina* (Sclater, 1856) **Andean White-crowned Manakin**

**Distribution**: Subtropical Andes from Venezuela south to Esmeraldas, Ecuador and San Martín, Peru

**Phylogenetic Position**: Clade A1

***P. c. coracina*** (Sclater, 1856)
**Type Locality:** Villavicencio, Meta, Colombia.
**Distribution**: Subtropical forests of the eastern slope of the Andes from western Venezuela to Morona-Santiago, Ecuador
**Phylogenetic Position**: (mtDNA) Member of Clade A1
**Lek Vocal Type**: 8 (*errrwer*).
**Call Vocal Type**: Unknown
***P. c. minima*** (Chapman et al., 1914)
**Type Locality:** West of Popayan, Cauca, Colombia
**Distribution**: Subtropical forests of western Cauca, Colombia south to Esmeraldas, Ecuador
**Phylogenetic Position**: (unsampled) Member of Clade A/A1
**Lek Vocal Type**: 9 (*reeee*)
**Call Vocal Type**: unknown
***P. c. bolivari*** (de Schauensee, 1950)
**Type Locality:** Murucucu, Bolivar, Colombia
**Distribution**: Subtropical forests of southern Córdoba, Colombia (not sampled)
**Phylogenetic Position**: (unsampled) Member of Clade A1
**Lek Vocal Type**: Unknown
**Call Vocal Type**: Unknown
***P. c. unica*** (de Schauensee, 1945)
**Type Locality:** Lomas de Isnos, Huila, Colombia
**Distribution**: Subtropical forests of Magdalena Valley, Antioquia to Huila, Colombia
**Phylogenetic Position**: (unsampled) Member of Clade A1
**Lek Vocal Type**: 11a (*weer-dink*) and 11b (*shureeep*)
**Call Vocal Type**: Unknown
***P. c. occulta*** (Zimmer, 1936)
**Type Locality:** Ucho, east of Chachapoyas, Amazonas/San Martin, Peru
**Distribution**: Eastern slope of the Andes from Zamora-Chinchipe, Ecuador (Freile, 2014) south to San Martín, and Huánuco, Peru, west of the Huallaga river
**Phylogenetic Position**: Member of Clade A/A1
**Lek Vocal Type**: 1 (*trill-dink*)
**Call Vocal Type**: 10 (*bree*)

*Pseudopipra anthracina* (Ridgway, 1906) **Western White-crowned Manakin**

**Type Locality:** Moravia, Costa Rica

**Distribution**: Subtropical Costa Rica to Western Panama

**Phylogenetic Position**: Clade A2

**Lek Vocal Type**: 4 (*jureeee*)

**Call Vocal Type**: Unknown

*Pseudopipra comata* (Berlepsch and Stolzmann, 1894) **Junín White-crowned Manakin**

**Type Locality:** Garita del Sol, Junin, Peru.

**Distribution**: Subtropical Andes of Peru from Cordillera Azul, Loreto (east and south of the Huallaga river) to southern Huánuco Pasco, Junín, and northern Cusco

**Phylogenetic Position**: Clade B

**Lek Vocal Type**: 1 (*trill-dink*)

**Call Vocal Type**: 10 (*bree*)

*Pseudopipra pygmaea* (Zimmer, 1936) **Huallaga White-crowned Manakin**

**Type Locality:** Chamicuros, Loreto, Peru

**Distribution**: Tropical forest of lower Huallaga river Valley, Peru

**Phylogeneic Position:** (mtDNA) Sister to Clade F

**Lek Vocal Type**: 2 (*deeeer*)

**Call Vocal Type**: 13 (*weeu*)

*Pseudopipra discolor* (Zimmer, 1936) **Napo White-crowned Manakin**

**Type Locality:** Puerto Indiana, Loreto, Peru

**Distribution**: Tropical forest in Napo, Ecuador and Loreto, Peru south to the Marañón river

**Phylogenetic Position:** Clade E

**Lek Vocal Type**: 2 (*deeeer*)

**Call Vocal Type**: 13 (*weeu*)

*Pseudopipra pipra* (Linnaeus, 1758) **Northern White-crowned Manakin**

**Type Locality:** Suriname

**Distribution**: Tropical forest of eastern Colombia, southern Venezuela, the Guianas, and Brazil north of the Amazon and west to the right (west) bank of the Putumayo river, Colombia

**Phylogenetic Position**: Clade D

**Lek Vocal Type**: 3 (*buzzzz*)

**Call Vocal Type:** 5 (*zeee*)

*Pseudopipra microlopha* (Zimmer, 1929) **Southern White-crowned Manakin**

**Distribution**: Tropical forest of eastern Peru south of the Huallaga river, Marañón river, and the Amazon river east to Pará, Brazil, and subtropical forests between the Huallaga river and Ucayali river

**Phylogenetic Position:** Paraphyletic, including Clade C without Clade C7

*P. m.* undescribed subspecies
**Distribution:** Subtropical forest from the Cushabatay area, northern Cordillera Azul, between the Huallaga river and Ucayali river, Loreto Province, Peru
**Phylogenetic Position:** Clade C2
**Lek Vocal Type**: 7 (*jeer*)
**Call Vocal Type**: 13 (*weeu*)
*P. m. microlopha* (Zimmer, 1929)
**Distribution**: Eastern Peru south of the Marañón river and Huallaga river, west to the Juruá river and Purus river, Brazil
**Phylogenetic Position:** Clade C1 excluding C2
**Lek Vocal Type**: 7 (*jeer*)
**Call Vocal Type**: 13 (*weeu*)
*P. m.* undescribed subspecies
**Distribution:** Right (east) bank of the Purus river to the left (west) bank of the Madeira river
**Phylogenetic Position:** Clade C3
**Plumage:** Not examined
**Lek Vocal Type:** Unknown
**Call Vocal Type**: 13 (*weeu*)
*P. m.* undescribed subspecies
**Distribution:** Right (east) bank of the Madeira river to the left (west) bank the Tapajós river
**Phylogenetic Position:** Clade C4
**Plumage:** Not examined
**Lek Vocal Type:** Unknown
**Call Vocal Type**: 13 (*weeu*)
*P. m.* undescribed subspecies
**Distribution:** Right (east) bank of the Tapajós river to the left (west) bank of the Xingu river
**Phylogenetic Position:** Clade C5
**Plumage:** Not examined
**Lek Vocal Type:** 6b (*zeeeee-tonk*)
**Call Vocal Type**: 13 (*weeu*)
*P. m. separabilis* (Zimmer, 1936)
**Type Locality:** Tapara, Xingu river, Brazil
**Distribution**: Tropical forest of the Xingu river east to central and southern Pará
**Phylogenetic Position:** Clade C6
**Lek Vocal Type:** 2 (*deeeeer*)
**Call Vocal Type**: 13 (*weeu*)

*Pseudopipra cephaleucos* (Thunberg, 1822) **Atlantic White-crowned Manakin**

**Type Locality:** Bahia, Brazil

**Distribution**: Tropical forest from Bahia south to northern Rio de Janeiro, Brazil

**Phylogenetic Position:** Clade C7

**Lek Vocal Type:** 6a (*zeeeee-tonk*)

**Call Vocal Type**: 13 (*weeu*)

## Supplementary Material

Archived Supplementary Data, including data files and *R* code can be found in the Mendeley Data Repository (LINK REMOVED). Mitochondrial sequences have been archived at NCBI GenBank (REMOVED). SNP datasets have additionally been archived at the European Variation Archive (Project: REMOVED, Analyses: REMOVED and REMOVED). *R* code will also be made available at the author’s GitHub repository: https://github.com/jakeberv.

## Acknowledgements

We thank the numerous researchers whose field collection efforts made this research possible, including Angelo P. Capparella, Camila Ribas, Ivandy Castro-Astor, Cecilia Fox, David E. Willard, Diego Ocampo, Donna Dittmann, Jacob S. Berv, John P, O’Neill, Karla Conejo, Katya Balta, Kenneth V Rosenberg, Louise M. Augustine, Peter E. Scott, Richard O. Prum, Steven W. Cardiff, Susan Allen-Stotz, Thomas Valqui, Tristan J. Davis, Ivan Prates and others. Tissue loans or skins were kindly received from the Divisão de Aves do Museu de Zoologia da Universidade Estadual de Feira de Santana (MZFS), the Field Museum of Natural History (FMNH), Instituto Nacional de Pesquisas da Amazônia (INPA), Kansas University Natural History Museum (KUNHM), Louisiana Museum of Natural History (LSUMZ), Museu de Zoologia da Universidade de São Paulo (MZUSP), Museu Paraense Emilio Goeldi (MPEG), the Academy of Natural Sciences of Drexel University (ANSP), Universidade do Estado do Rio de Janeiro (UERJ), the University of Alaska Museum (UAM), the University of Nevada, Las Vegas (UNLV), the Yale Peabody Museum of Natural History (YPM), the Cornell Museum of Vertebrates (CUMV), the American Museum of Natural History (AMNH), the Museum of Comparative Zoology (MCZ), and the United States Museum of Natural History (NMNH). We thank the curators and collections managers of these institutions for facilitating these loans: Camila Ribas, John Bates, Mark Robbins, Donna Dittman, Alexandre Aleixo, Caio Graco Machado, Joel Cracraft, Luís Fábio Silveira, Kevin Winker, Maria Alice Santos Alves, Kristof Zyskowski, Nate Rice, Charles Dardia, John Klicka, and others. We thank Eliot Miller, Emma Greig, Matthew Medler, Greg Budney, Eduardo Iñigo and Andrew Spencer at the Macaulay Library for assistance with audio recordings. We thank members of the Lovette Lab, Daniel Lane and four anonymous reviewers, for comments on the research and the manuscript. We gratefully acknowledge the bioRxiv for hosting the pre-print version of this article. The illustration depicted in Fig. 1 is reproduced with permission from the Handbook of Birds of the World, Lynx Editions, Barcelona. J. S. B. and R. O. P. conceived of the research. J. S. B. and L. C. performed laboratory and bioinformatic processing, with input and support from I. J. L. We thank Bronwyn Butcher for help with laboratory protocols. J. S. B. and L. C. analyzed the genomic data, and interpretation of the results was supported by input from all authors. T. J. F. downloaded, processed, and analyzed lekking vocalization data. C. C. R. and I. C. R. facilitated critical sample acquisitions from Amazonia and the Atlantic Forest. I. J. L. and R. O. P. provided detailed guidance and feedback on the manuscript. R. O. P. examined plumage phenotypes and proposed a revised taxonomy. The manuscript was written by J. S. B. with input from all authors.

