## Supplementary Appendix for "Genomic phylogeography of the White-crowned Manakin *Pseudopipra pipra* (Aves: Pipridae) illuminates a continental-scale radiation out of the Andes"

#### Expanded Taxonomic Summary

*Taxonomic history*

For most of its history, the manakin species *pipra* was placed in the genus *Pipra* (Snow, 1979). Prum (1990a, 1990b); and Prum (1992) recognized that the traditional *Pipra* (*sensu* Snow 1979) was polyphyletic and placed the species *pipra* in *Dixiphia* (Reichenbach, 1850) which was recognized as a junior synonym of *Pipra*. Recently, Kirwan et al. (2016) demonstrated that the genus *Dixiphia* was unavailable for *pipra* because it is actually a junior synonym of the tyrannid genus *Arundinicola*. They named the new genus *Pseudopipra* for *pipra*.

*Summary preface*

There are at least eight genetically well-differentiated, and phenotypically diagnosable lineages of *Pseudopipra*, and this is reflected in our proposed taxonomic revisions. Three of these species are polytypic (i.e., contain multiple subspecies). One of the species we recognize– *P. microlopha* – is paraphyletic with respect to another species, *P. cephaleucos,* because we recognize that speciation may not threaten the lineage identity of a paraphyletic group. Plumage and vocal data from other newly identified clades may support the recognition of additional species including a lowland-derived montane population from the northern Cordillera Azul (Clade C2; Fig. 3), and the currently unnamed lineages from the southern Amazon Basin (Clades C3, C4, and C5).

*Linnaean Taxonomy*

***Pseudopipra coracina*** (Sclater, 1856) **Andean White-crowned Manakin**
**Distribution**: Subtropical Andes from Venezuela south to Esmeraldas, Ecuador and San Martín, Peru.
**Phylogenetic Position**: Clade A1, plus multiple unsampled subspecies from the Colombian and Ecuadorian Andes (Figure 3).
**Comments:** This apparently monophyletic group of northern Andean populations includes five currently recognized subspecies, each of which may be a distinct species. Three of these subspecies–*coracina, minima,* and *occulta*– have unique, highly differentiated vocal types, and diagnosable plumage differences.

***P. c. coracina*** (Sclater, 1856)
**Type Locality:** Villavicencio, Meta, Colombia
**Distribution**: Subtropical forests of the eastern slope of the Andes from western Venezuela to Morona-Santiago, Ecuador.
**Phylogenetic Position**: Based on mtDNA sampled, a member of Clade A1.
**Plumage:** Males are moderately glossy on the back. White crown feathers are long with extensive black bases. Crowns are sometimes slightly grayish. Females are olive green with lighter yellow belly, and olive gray crown with more olive cheeks.
**Lek Vocal Type**: 8 (*errrwer*).

**Call Vocal Type**: Unknown

***P. c. minima*** (Chapman et al., 1914)
**Type Locality:** West of Popayan, Cauca, Colombia
**Distribution**: Subtropical forests of western Cauca, Colombia south to Esmeraldas, Ecuador
**Phylogenetic Position**: Not Sampled, but a likely member of Clade A1 (Figure 3).
**Plumage:** Males are moderately glossy; crown feathers are entirely white to their bases. No females were observed. (Chapman et al., 1914) reported that *minima* is smaller than *anthracina*, and that males lack prominent gray tips on undertails. Freile (2014) reported one specimen of a female from San Javier, Esmeraldas, Ecuador (100 meters) and provisionally identified it as *minima.* The specimen is bright olive above and below with a slightly grayish olive grown. However, this specimen is from a substantially lower altitude than Colombian records of *minima*, so it may represent an altitudinal migrant or a distinct population.
**Lek Vocal Type**: 9 (*reeee*)
**Call Vocal Type**: Unknown

***P. c. bolivari*** (de Schauensee, 1950)
**Type Locality:** Murucucu, Bolivar, Colombia
**Distribution**: Subtropical forests of southern Córdoba, Colombia. (Not Sampled)
**Phylogenetic Position**: Not Sampled, but a likely member of Clade A1 (Figure 3).
**Plumage:** None observed. Apparently known only from the type specimen from Cerro Murucucú, Córdoba, Colombia. de Schauensee (1950) described this male specimen as having entirely white feathers in the forecrown, like *minima* and *unica,* but hindcrown feathers basally black like *coracina.*
**Lek Vocal Type**: 11a (*weer-dink*)
**Call Vocal Type**: Unknown

***P. c. unica*** (de Schauensee, 1945)
**Type Locality:** Lomas de Isnos, Huila, Colombia
**Distribution**: Subtropical forests of Magdalena Valley, Antioquia to Huila, Colombia.
**Phylogenetic Position**: Not Sampled, but a likely member of Clade A1 (Figure 3).
**Plumage:** Males are moderate glossy, with long crown feathers that are white to their bases. Females are olive green above, and slightly gray on the crown; underparts uniform olive. (de Schauensee, 1945) described *unica* as glossier than coracina, with longer tail and very long crest.
**Lek Vocal Type**: 11a (weer-dink) and 11b (shureeep)
**Call Vocal Type**: Unknown

***P. c. occulta*** (Zimmer, 1936)
**Type Locality:** Ucho, east of Chachapoyas, Amazonas/San Martin, Peru
**Distribution**: Eastern slope of the Andes from Zamora-Chinchipe, Ecuador Freile (2014) south to San Martín, and Huánuco, Peru, west of the Huallaga river.
**Phylogenetic Position**: Clade A1 (Figure 3).

**Plumage:** Males are glossy with dark gray bases to crown feathers. Females are dark olive with dark gray crown and gray throat. Zimmer (1936) described occulta as similar to *comata* but adult males with the occipital feathers slightly shorter and with the crown and occipital feathers sooty at the base instead of entirely white.
**Lek Vocal Type**: 1 (*trill-dink*)
**Call Vocal Type**: 10 (*bree*)

***Pseudopipra anthracina*** (Ridgway, 1906) **Western White-crowned Manakin**
**Type Locality:** Moravia, Costa Rica
**Distribution**: Subtropical Costa Rica to Western Panama
**Phylogenetic Position**: Clade A2 (Figure 3).
**Plumage:** Males less lustrous on back than all other *Pseudopipra* populations, white crown feathers gray or dark gray at base. Females are olive green with slaty crown and face. Ridgway (1906) considered *anthracina* to have shorter wings, smaller beak, less lustrous plumage than *pipra* with undertails tipped with gray.
**Lek Vocal Type**: 4 (*jureeee*)
**Call Vocal Type**: unknown

***Pseudopipra comata*** (Berlepsch and Stolzmann, 1894) **Junín White-crowned Manakin**
**Type Locality:** Garita del Sol, Junin, Peru
**Distribution**: Subtropical Andes of Peru from Cordillera Azul, Loreto (east and south of the Huallaga river) to southern Huánuco, Pasco, Junín, and northern Cusco.
**Phylogenetic Position**: Clade B (Figure 3).
**Plumage:** Males are glossy black above, crown feathers longer and entirely white to their bases. Females are bright olive green above, gray on crown and face, slightly gray on throat, dark olive below, and slightly dark gray on the belly.
**Lek Vocal Type**: 1 (*trill-dink*)
**Call Vocal Type**: 10 (*bree*)
**Comment:** *P. comata* is also composed of two well differentiated subclades. The northern clade (B1) is known from southern Cordillera Azul in Loreto, Peru. The southern clade (B2) is known from Pasco, Junín, and Cusco. The type locality of *comata* is Vitoc, Junín within the southern clade. Birds of from the Cerros del Sira (9°30'S 74°47'W; AMNH 820866, 820952) are apparently *comata,* but we do not have any DNA or vocal data to confirm that. Further investigation plumage and behavioral is necessary to determine whether the southern Cordillera Azul populations should be recognized as a distinct, new taxon.

***Pseudopipra pygmaea*** (Zimmer, 1936) **Huallaga White-crowned Manakin**
**Type Locality:** Chamicuros, Loreto, Peru
**Distribution**: Tropical forest of Lower Huallaga river Valley, Peru
**Phylogeneic Position:** Sister to Clade F (mtDNA) (Figure 3).
**Plumage:** Males: Glossy, with black bases to crown feathers. Females are olive above and gray below with a band of olive across the chest, crown and face only slightly darker than back, not gray. (Zimmer, 1936) described males as having long crest with gray bases, crown sometimes slightly ashy; females are much paler than *occulta*; throat and belly decidedly more whitish, breast paler duller green; lighter even than *microlopha*.
**Lek Vocal Type**: 2 (*deeeer*)

**Call Vocal Type**: 13 (*weeu*)
**Comment**: Lowland populations along the Huallaga river have been named *pygmaea* (Zimmer, 1936). Our four samples of *pygmaea* from Jeberos, Peru did not yield sufficient quality DNA for RADseq, but all four had a phylogenetically distinct mtDNA haplotype which placed this lineage as the sister group to all other lowland populations of *Pseudopipra*. These populations have song type 2, which appears to be shared plesiomorphically with *P. discolor* and *P. microlopha separabilis* from Para, Brazil.

***Pseudopipra discolor*** (Zimmer, 1936) **Napo White-crowned Manakin**
**Type Locality:** Puerto Indiana, Loreto, Peru
**Distribution**: Tropical forest in Napo, Ecuador and northern Loreto, Peru south to the Marañón river.
**Phylogenetic Position:** Clade E (but actually derived from Clade C) (Figure 3).
**Distribution**: Tropical forest in Napo, Ecuador south to the Marañón river.
**Plumage:** Males are glossy black above, white crown feathers with black or dark gray bases. Females are dusky olive overall, slightly grayer on crown, and grayer belly. (Zimmer, 1936) described male *discolor* as glossier and bluer above than *pipra.*
**Lek Vocal Type**: 2 (*deeeer*)
**Call Vocal Type**: 13 (*weeu*)
**Comment:** This lineage was found to have a distinct unique history, with subsequent introgression with adjacent populations of the northern Amazonian clade. The nature of this introgression indicates this lineage may be best recognized as a distinct hybrid species.

***Pseudopipra pipra*** (Linnaeus, 1758) **Northern White-crowned Manakin**
**Type Locality:** Suriname
**Distribution**: Tropical forest of eastern Colombia, southern Venezuela, the Guianas, and Brazil north of the Amazon. West to the right (north) bank of the Putumayo river, Colombia.
**Phylogenetic Position**: Clade D (Figure 3).
**Plumage:** Males are glossy black above, crown feathers longer with extensive black bases. Females are dark olive above, olive below, grayer on belly, and occasionally only slightly darker gray on crown.
**Lek Vocal Type**: 3 (*buzzzz*)
**Call Vocal Type:** 5 (*zeee*)

***Pseudopipra microlopha*** (Zimmer, 1929) **Southern White-crowned Manakin**
**Distribution**: Tropical forest of eastern Peru south of the Marañón river, and south of the Amazon east to Pará, Brazil, and subtropical forests between the Huallaga river and Ucayali river.
**Phylogenetic Position:** Paraphyletic, including Clade C without Clade C7 (Fig. 3).
**Comments:** A paraphyletic group (with respect to *P. cephaleucos* from Brazilian Atlantic forest) which includes three, currently recognized subspecies, and four additional genetically well-supported monophyletic subgroups that may be recognized as new taxa. Furthermore, we identified a genetically distinct montane clade from the highlands between the Huallaga river and the Ucayali river that has not been previously described and may have distinct plumage and vocal characters.

***P. m.* undescribed subspecies**
**Distribution:** Subtropical forest from the highlands between Huallaga river and Ucayali river. All samples are from a single locality in Cushabatay area, northern Cordillera Azul, Loreto, Peru; 7.08333° S, 75.65° W.
**Phylogenetic Position:** Clade C2 (Figure 3).
**Plumage:** Not examined.
**Lek Vocal Type**: 7 (*jeer*)
**Call Vocal Type**: 13 (*weeu*)

***P. m. microlopha*** (Zimmer, 1929)
**Type Locality:** Puerto Bermudez, Pasco, Peru
**Distribution**: Eastern Peru south of the Marañón river and Huallaga river west to the Juruá river and the Purus river, Brazil.
**Phylogenetic Position:** Apparently paraphyletic, Clade C1 excluding C2 (Figure 3).
**Plumage:** Males are glossy black above, with black or dark gray bases to white crown feathers. Females are dark olive above, occasionally with slightly gray crown, olive below, and graying on the belly.
**Lek Vocal Type**: 7 (*jeer*)
**Call Vocal Type**: 13 (*weeu*)

***P. m.* undescribed subspecies**
**Distribution:** Right (east) bank of the Purus river to the left (west) bank Madeira river.
**Phylogenetic Position:** Clade C3 (Figure 3).
**Plumage:** Not examined.
**Lek Vocal Type:** Unknown
**Call Vocal Type**: 13 (*weeu*)

***P. m.* undescribed subspecies**
**Distribution:** Right (east) bank of the Madeira river to the left (west) bank of the Tapajós river.
**Phylogenetic Position:** Clade C4 (Figure 3).
**Plumage:** Not examined.
**Lek Vocal Type:** Unknown
**Call Vocal Type**: 13 (*weeu*)

***P. m.* undescribed subspecies**
**Distribution:** Right (east) bank of the Tapajós river to the left (west) bank of the Xingu river.
**Phylogenetic Position:** Clade C5 (Figure 3).
**Plumage:** Not examined.
**Lek Vocal Type:** 6b (*zeeeee-tonk*)
**Call Vocal Type**: 13 (*weeu*)

***P. m. separabilis*** (Zimmer, 1936)

**Type Locality:** Tapara, Xingu river, Brazil
**Distribution:** Right (east) bank of the Xingu river east to central and southern Pará.
**Phylogenetic Position:** Clade C6 (Figure 3).
**Plumage:** Males are moderately glossy above, crown long with large, dark gray feather bases. Predefinitive male plumage light olive above, gray below, with medium gray crown.
Females are light olive above, light grayish below with olive wash on the breast. Zimmer (1936) commented that adult males and females not distinguishable from *separabilis,* but he identified the distinct predefinitive male plumage.
**Lek Vocal Type:** 2 (*deeeer*)
**Call Vocal Type**: 13 (*weeu*)

***Pseudopipra cephaleucos*** (Thunberg, 1822) **Atlantic White-crowned Manakin**
**Type Locality:** Bahia, Brazil
**Distribution**: Tropical forest from Bahia south to northern Rio de Janeiro, Brazil.
**Phylogenetic Position:** Clade C7 (Figure 3).
**Plumage:** Males are glossy black with a long and slightly gray crown. Crown feather have extensive dark gray bases. Predefinitive males have olive backs, pure white or grayish white crowns, and slate gray on the face, throat, and belly. Females have olive back, dusky gray on head, gray below, slightly olive on breast, lighter on belly.
**Lek Vocal Type:** 6a (*zeeeee-tonk*)
**Call Vocal Type**: 13 (*weeu*)

#### Additional Results and Discussion

*Expanded ddRAD laboratory methods*

Briefly, we digested equal quantities of genomic DNA from each sample in individual reactions with two restriction enzymes, SbfI and MspI (New England Biolabs, MA), and ligated adapters on both ends. The 5' adapters contained one of 20 unique 5-7 bp barcodes, whereas the 3' adapter was common to all samples. We pooled groups of samples with unique 5' barcodes and subsequently size selected DNA fragments that were between 400-700 bp using a Blue Pippin (Sage Science, MA). For each group of size-selected samples, we incorporated unique Illumina TruSeq adapters by performing 11 cycles of PCR. The combination of 5' barcodes and TruSeq adapters was unique to each sample. A total of 13 groups of pooled samples with different Illumina TruSeq adapters were combined in equimolar proportions into two libraries. We sequenced both libraries on four lanes of Illumina HiSeq 2500 at the Cornell University Institute for Biotechnology, obtaining single-end 151 bp sequences.

*Assembly of sequencing reads into RAD loci*

We obtained 497 million raw 151 bp reads (~75 Gb) for 241 individuals. We first assessed the overall read quality with FastQC (Andrews, 2010) and trimmed lower quality bases at the 3' end with FASTX-Toolkit (Gordon and Hannon, 2010). The trimmed sequences were 145 bp in length. Using FASTX-Toolkit we filtered-out lower quality reads if they had a single base below a Phred quality score of 10 and/or more than 5% of bases with quality between Phred 10 and 20.

The quality filtered reads were demultiplexed with the "process_radtags" program from the STACKS 1.44 bioinformatics pipeline (Catchen et al., 2013). During the demultiplexing step we also discarded reads that had not passed the Illumina filter, had adapter contamination, lacked barcodes used for multiplexing, or did not contain an SbfI cut site. This step removed the inline barcodes and trimmed all reads to an equal length, the length of the reads that contained a 7 bp barcode.

After demultiplexing and filtering, we retained an average of 1.1 ± 0.4 million 138 bp sequences per individual. We downloaded the *Manacus vitellinus* (GCA_001715985.1) reference genome from www.ncbi.nlm.nih.gov, and aligned the reads from each individual using bowtie2 2.3 (Langmead et al., 2009), as recommended by Paris et al. (2017); Shafer et al. (2017). We assembled the mapped reads into RAD loci using the reference-based pipeline in STACKS, executed with the ref_map script which runs the modules "pstacks/cstacks/sstacks". We subsequently ran the error correction module "rxstacks" and a final iteration of "cstacks/sstacks". We set the parameters to a minimum coverage of 20 (*m*) and up to two differences among aligned loci of different individuals (*n*). The reference-based assembly produced a catalogue with 47,046 RAD loci. Seven samples were discarded due to high proportions of missing data in the final assemblies; our final RAD dataset thus comprised 234 individuals. We used several filters in the "populations" module from Stacks to generate different sets of bi-allelic SNPs (see main text). For sequence datasets, an additional 5' 6bp was trimmed to remove the SbfI cut site (below).

*Additional phylogenetic results and comments on mutational spectra*

At the level the concatenated alignments, the "20% missing" dataset had a total of 2,548 132 bp loci (340,956 sites), 7.92% missing sites, 8,063 parsimony informative sites, and 5,709 variable parsimony-uninformative sites. The "50% missing" dataset had a total of 4,763 132 bp loci (626,868 sites), 20.48% missing sites, 15,365 parsimony informative sites, and 11,221 variable parsimony-uninformative sites. The "80% missing" dataset had a total of 7,907 132 bp loci (1,039,632 sites), 38.89% missing sites, 24,450 parsimony informative sites, and 17,839 variable parsimony-uninformative sites. Across these datasets, alpha from the GTR+G model was << 1 (0.069, SD=$6.033\times10^{-4}$), indicating high among site rate heterogeneity for these ddRAD loci. Chi-square tests of base compositional heterogeneity rejected the hypothesis of compositional homogeneity (Chi-sq= 7822.93, df=702, P << 0.05), with slight bias observed on the AT-GC axis of compositional variation (A: 0.24968, C: 0.25225, G: 0.24301, T: 0.24301, on the largest 80% dataset). Maximum likelihood estimates of transition rates were ~8x transversion rates (A<-->G: 7.34 x G <--> T, C <--> T: 8.13 x G <--> T), as estimated in RAxML. Estimated rates among other nucleotide classes were ~1 relative to the fixed G <--> T rate, suggesting that GTR+G may be over-parameterized for this ddRAD dataset.

*Reconstruction of mitochondrial ND2 gene tree*

After obtaining mitochondrial DNA sequences for 168 individuals (Supplementary Table 1), we aligned these sequences using MAAFT (Standley and Katoh, 2013). The alignment was visually inspected and trimmed in Sequencher (Gene Codes Corporation, 2010), and then analyzed in IQ-TREE 1.6.10 (Chernomor et al., 2016; Hoang et al., 2017; Kalyaanamoorthy et al., 2017; Schmidt et al., 2014; Trifinopoulos et al., 2016). We partitioned by codon position and generated a maximum likelihood tree using the MFP+MERGE model search and partitioning option, with 1000 ultrafast bootstrap replicates. MFP+MERGE detected that an optimal scheme comprised of three partition-models for each of the three codon positions (CP1: TIM2+F+I; CP2: TIM2+F+G4; CP3: TIM2+F+G4). Nodes recovered with ultrafast bootstrap scores < 95 were collapsed. The recovered topology was entirely congruent with the topology presented in the main text as derived from ddRAD data, with a few exceptions (Supplementary Figure 17). Our mtDNA dataset included individuals from subspecies *coracina* and *pygmaea* which were derived from low quality tissue samples (and hence were not suitable for ddRAD sequencing). This enabled us to make a preliminary assessment of their phylogenetic affinities (main text), though nuclear genomic data should be collected in future studies. Notably, the introgressed western Napo lineage has mtDNA haplotypes which are members of the southern amazon clade (BS 98), which is consistent with the scenario of hybrid origin and introgression we develop in the main text. Because mtDNA is inherited matrilineally, a potential implication of this pattern is that the introgressed Napo lineage (S2a/S2 in Figure 6) was created when southern progenitor females were introgressed with northern males.

*Reconstruction of ancestral ranges*

We performed a Bayesian stochastic character mapping analysis (Bollback, 2006; Huelsenbeck et al., 2003) to estimate the ancestral habit of *Pseudopipra*. In brief, we coded lineages as Andean, Central American, or Lowland, applied a bi-directional Mk model ("ARD") and performed 100 simulations (see supplemental R scripts) using the RAxML topology. We used the SIMMAP implementation in phytools (Revell, 2012). We also performed a similar analysis under maximum parsimony using the phangorn R package (Schliep, 2011) and recovered identical results, which unambiguously reconstruct the ancestral habit of *Pseudopipra* to be montane Andean (see Figure 5 in the main text for a summary of these results).

*STRUCTURE - additional notes*

The Evanno method applied to the whole dataset detected a significant shift in the rate of change of the log probability of the data between K1 and K2, indicating a deep hierarchical split in the data. As STRUCTURE also infers the degree of admixture among individuals, these assignments are not directly analagous to K-means phenetic cluster assignments, which lump individuals categorically based on overall genetic similarity.

#### Descriptive Population Genetic Statistics

*Methods*

For population genetic statistics, we considered eighteen population-areas (Figure 3, 4). Most of these populations are delimited by clear geographic barriers (e.g., rivers in the cases of previously identified areas of endemism, the Andes, or the Cerrado belt) and have strong phylogenetic support. Two subgroups within the broad Northern Amazonian + Guiana Shield lowland clade were defined on the basis of low support monophyly in the RAxML analysis *and* coincidence with geographic features. One of these comprised individuals unambiguously assigned to the northern Amazonian clade in phylogenetic analysis, but which were also restricted to the eastern Napo area of endemism, east of the Putumayo river (brown markers in Figures 3, 4, "weakly resolved eastern Napo" – abbreviated in R code and Supplementary Figures as "GSNapo"). The second groups comprised individuals found near the coasts in Suriname and the Brazilian state of Amapá, east of the Essequibo river (pale blue markers in Figures 3, 4: "Suriname + Amapá" – abbreviated in R code and Supplementary Figures as "GSSR"). Another subgroup was defined on the basis of restriction to the Jaú area of endemism (pale yellow markers in Figures 3, 4: "unresolved Jaú" – abbreviated in R code and Supplementary Figures as "GSImeri"). Lastly, a fourth group of individuals included all other individuals in the lowland northern Amazon clade, restricted to the Guiana Shield (green markers in Figure 3, 4: "weakly resolved Guiana Shield" – abbreviated in R code and Supplementary Figures as "GS"), comprising individuals east of the Jaú group (above), and west of those in the Suriname + Amapá group. The primary geographic barriers in this region separating western and eastern Guiana Shield populations seems to be the Guiana Highlands, which is where tepuis are found, as well as the Essequibo river.

For these descriptive analyses, we focus on the aforementioned eighteen areas as units of comparison because focusing on broader populations delimited by cluster analyses would likely generate statistics biased by population sub-structure—i.e., lower than expected heterozygosities (Wahlund, 1928). Further, groups delimited by broader cluster assignments may be more reflective of ancestral populations, and therefore not indicative of presently restricted groups (ie, inappropriately moving migrants back to their source populations, Kuhner, 2006). Statistics are reported from analyses with Dataset 2 (Table 1), as this includes the largest number of putatively unlinked markers (the first SNP from each of 2,581 ddRAD loci), unless otherwise indicated.

To estimate a measure of genetic diversity across these sampling regions, we calculated the rarefied allelic richness per population (restricted to populations comprising > 5 individuals) using the allelic.richness function in the hierfstat R package (Goudet, 2005), after removing all sites with missing genotypes (Supplemental R Script).

We also calculated the inbreeding coefficient $F_{IS}$, defined as $(H_S – H_I)/H_S$, where $H_I$ is the mean expected heterozygosity per individual within subpopulations, and $H_S$ is the mean expected heterozygosity within random mating populations (Goudet, 2005). We generated 100,000 bootstrapped estimates of $F_{IS}$, sampling over loci per population, using the boot.ppfis function in hierfstat (Goudet, 2005). For recent hybrid individuals, F1s should be more outbred (relative heterozygosity) than their parental genotypes. We tested the hypothesis that our sample of the introgressed western Napo population includes F1 individuals by estimating the inbreeding coefficient for a simulated F1 population, comprised of the progenitor lineages discussed in the main text. We generated a simulated F1 population using the hybridize function in adegenet (Jombart, 2008), and then estimated it's inbreeding coefficient as described above to compare to empirical estimates from source populations.

To perform a preliminary assessment of the potential for evolutionary processes deviating from the assumptions of Hardy-Weinberg equilibrium, we applied the hw.test function in the pegas R package (Paradis, 2010) with 1000 Monte Carlo permutations of alleles to compute an exact p value for each locus within each population. To assess the assumption of linkage intrinsic to most model-based analyses in this study, we computed the Standardized Index of Association $\bar{r}_d$ (Agapow and Burt, 2001; Brown et al., 1980) within populations using the poppr summary function in the poppr R package (Kamvar et al., 2014), and estimated p values with 1000 permutations. We estimated pairwise Weir and Cockerham's (Weir and Cockerham, 1984) $F_{st}$ among all 18 areas, and evaluated significance using 1000 bootstrapped datasets to estimate 95% confidence intervals using the 'assigner' R package (Gosselin et al., 2016).

We estimated the potential for sex-biased dispersal in *Pseudopipra* using sex assignments derived from voucher specimen records (135/232 samples) and the sexbias.test function in hierfstat (Goudet, 2005; Goudet et al., 2002), with mAIc test. We ran this test on all eight SNP datasets, and assessed significance with 1000 permutations.

Lastly, we quantified differentiation among two hierarchical strata recapitulating 1) deep coalescent structure (6 groups as identified by SVDquartets (~K5 from STRUCTURE + putative introgressed Napo hybrids as a separate group), and 2) populations identified in phylogenetic analyses which coincide with geographic barriers (18 groups), with analysis of molecular variance (AMOVA) (Excoffier et al., 1992). We used the poppr.amova wrapper function in the poppr R package (Kamvar et al., 2014) to perform AMOVA on adegenet genind objects, set to use the ade4 implementation of AMOVA with 1000 permutations to assess significance. For AMOVA calculations we used dataset 1, to minimize within individual variance. Code to perform all of these analyses is provided in the supplementary R scripts.

*Results*

Average missing data in SNP datasets D1-D4 (which required a SNP to be present in at least 80% of individuals), was generally low: mean: ~9.6%, SD: 5.4% and ranged from ~2.4% (Jaú subgroup of the Guiana Shield clade) to a maximum of ~20.5% (Costa Rica, though this was somewhat of an outlier – 75% of these areas had less than 13% missing data overall). Despite the fact that our sampling among areas delimited by geographic boundaries had high variance relative to the mean (mean: 12.94 [1 - 70], SD: 16.95, CoV: 1.31), the rarefied estimates of allele count in each of 13 areas (with > 5 individuals, and after filtering out all sites with missing genotypes) were similar. For dataset 2 (2581 unlinked SNPs): mean number of alleles: 297.76, SD: 6.44, CoV: 0.022. The greatest allelic diversity was observed in the Jaú (n=14, 305.75 alleles) and introgressed western Napo population (n=10, 303.8 alleles). The lowest allelic richness was found in the Rio de Janeiro (n=10, 287.15) and Bahia (n=6, 285.48 alleles) Atlantic Forest populations, followed closely by Panamanian populations (n=5, 292 alleles). These results are generally consistent across SNP datasets and with our EEMS analysis (Supplementary Figure 12). Summary below:

| **Population** | **Alleles** |
|---|---|
| Atlantic Forest (Bahia) | 285.45 |
| Atlantic Forest (Rio) | 287.15 |
| Central America (Panama) | 292.00 |
| South Andean Peru (South) | 294.00 |
| Eastern Inambari endemic | 297.42 |
| Xingu endemic | 297.52 |
| Weakly resolved Guiana Shield (western) | 299.46 |
| Weakly resolved Suriname + Amapá | 299.56 |
| Inambari endemic (western) | 302.47 |
| Weakly resolved eastern Napo | 302.49 |
| Tapajós endemic | 303.75 |
| Western Napo introgressed lineage | 303.80 |
| Unresolved Jaú | 305.76 |

Using dataset 2, we detected a number of populations to have positive mean inbreeding coefficients ($F_{IS} > 0.1$, Supplementary Figure 9), with lower 95% confidence intervals > 0. Panamanian, Costa Rican, South Andean (North clade), Rondônia, and Espírito Santo populations had 95% confidence intervals which overlapped zero, and thus cannot be confidently inferred to have positive or negative $F_{IS}$. However, the simulated F1 population for the western Napo had significantly negative $F_{IS}$, as expected for outbred F1 hybrids. This pattern implies that the contemporary western Napo population, with a signature of significant introgression and which was detected to have a positive $F_{IS}$ (lower 95% > 0.1), is not likely to include F1 individuals. Indeed, the confidence intervals for eastern Napo, Jaú, Inambari and western Napo populations, are mostly overlapping, with similar means (mean of mean estimates ~0.17, SD of mean estimates ~0.02, Supplementary Figure 9). Notably, the consistently positive level of inbreeding (median ~0.16, 95% CI [0.11-0.21]) among most *Pseudopipra* populations may be attributable to a highly polygynous lek mating system (Smith, 1979; Stopher et al., 2012; Waser et al., 1986).

After correcting for multiple tests with the Benjamin & Hochberg correction, exact tests of Hardy-Weinberg equilibrium using dataset 2 suggested most loci in most populations were in equilibrium. However, a small number of loci in the western Guiana group (19 loci), Suriname+Amapá (8 loci) and Tapajós (3 loci) areas were identified as being out of Hardy-Weinberg equilibrium. The role of selection and non-neutral processes in driving population genetic differentiation in *Pseudopipra* should be investigated in more detail in future studies. Estimates of $\bar{r}_d$ within these populations indicated that there was also no strong evidence of linkage among loci within populations after correcting for multiple tests, except for the Tapajós area, in which weak linage was detected ($\bar{r}_d$: 0.005956, p= 0.001). Application of the mAIc test (Goudet, 2005; Goudet et al., 2002) indicated that there was no evidence of strong sex biased dispersal (p > 0.5 across all SNP datasets).

Lastly, an AMOVA detected significant population differentiation at all evaluated levels, including between coalescent groups (well supported clades from SVDquartets) (~32%) and between samples within coalescent units (~5%) ($p < 0.001$ for all). Re-running the same AMOVA with evolutionary distances estimated with RAxML branch lengths (instead of the default allelic distance) indicated the same pattern, but with more of the variance explained by coalescent and population level strata (41.3% and 12.6% respectively). Both AMOVA analyses detected a significant proportion of the variance attributable to within sample variance (62% and 46% respectively).

In general, our analyses speak to some broad observations about the sensitivity of various methods to detecting population genetic structure in ddRAD data. Methods based on clustering of SNP data were the least sensitive (prone to lumping) to population structure, whereas phylogenetic analyses were the most sensitive to population structure (prone to splitting). Thus, different phylogenetic and population genetic cluster analyses have detected different levels of hierarchical structure in the data.

#### Isolation by distance and the effect of geography

The evolutionary history of *Pseudopipra* within the Amazon basin appears to be deeply connected to the South American landscape, adding additional support to a rich body of literature endorsing this hypothesis (Brumfield, 2012; Cracraft and Prum, 1988). For virtually all evaluated cases, we find significant effects of geographic barriers on structuring genetic variation within this species complex, including the Amazon River and many associated tributaries (Table 2, Supplementary Table 2b, Figure 7). Further afield, the "dry-diagonal" Cerrado belt appears to have strongly isolated Atlantic Forest lineages from their southeastern Amazonian Xingu relatives, as do the Andes exhibit a disproportionate effect on divergence between Peruvian foothills populations and Central American lineages (with the caveat that our sampling in that area is sparse, so our power to infer spatial patterns is necessarily limited).

The establishment of the Amazonian river system has recently been questioned as a driver of species—level variation across key areas in the Neotropics (Oliveira et al., 2017; Santorelli et al., 2018). These studies used distributional data to infer the effects of key proposed barriers and concluded that while large rivers clearly limit some Amazonian species—the large number of exceptions to this "rule" point towards alternative speciation mechanisms as the norm, rather than as the exception. Indeed, rivers can plausibly function as contemporary species limits without being the source of such limits (Santorelli et al., 2018). In the case of *Pseudopipra,* river barriers have clearly contributed to contemporary patterns of genetic diversity, regardless of whether or not the formation of the Amazonian drainage system was the primary driver of generating that diversity. Importantly, studies which rely on distributional data alone are limited in that their statistical power is contingent on the accuracy of species and subspecies delimitation. In the biogeographic context of the Amazon, this is likely to be enormously underestimated for birds (Brumfield, 2012; Smith et al., 2014). This fundamental limitation in our knowledge of cryptic avian diversity is therefore likely to bias inferences derived from distributional data, which is based on mostly untested species limits. Indeed, most studies that use genetic data to investigate the effect of river or other physical barriers in structuring Neotropical avian diversity have inferred strong, though varying effects (e.g. Harvey and Brumfield, 2015; Moore et al., 2008; Naka and Brumfield, 2018; Silva et al., 2019).

A number of authors have also noted that the practice of identifying genetic clusters with model based approaches often fail to appropriately account for the effects of isolation by distance (Guillot et al., 2013), and various methods are in development to improve our ability to model such correlated phenomena (Botta et al., 2015; Bradburd et al., 2018; Bradburd et al., 2013; Petkova et al., 2015). STRUCTURE in particular has been highlighted as potentially suffering from over-estimating K as a consequence of spatial autocorrelation in widely distributed genetic data (Bradburd et al., 2018). Our STRUCTURE analysis appears to exhibit this behavior for the southern Amazon, with a genetic cline of admixture that falls on a longitudinal gradient across the southern Amazon and ends in the well differentiated Atlantic Forest Rio de Janeiro population. While it is plausible that isolation by distance, combined with physical barriers to gene flow, could generate a similar pattern (as implied by our phylogenetic analyses), it is important to keep this caveat in mind when interpreting STRUCTURE results. For example, STRUCTURE may suggest that a scenario of K2 with an admixture gradient between two populations is preferred, when K1 with an isolation by distance effect may be a better description and more biologically plausible model for the data (Bradburd et al., 2018). The degree to which this kind of spatial autocorrelation confounds admixture analyses remains an open and important area of inquiry. Our EEMS analysis attempts to circumvent this issue, assuming a more biologically realistic process of non-homogeneous but continuous IBD across a complex landscape.

#### Notes on congruent patterns with *Ceratopipra*

Within the manakins, several molecular phylogenies (Leite et al., 2021; Ohlson et al., 2013) have placed *Pseudopipra* as sister to the genus *Ceratopipra*, which includes five well-recognized species that are extensively codistribtuted with *Pseudopipra.* The geographic breakpoints among these *Ceratopipra* species are highly concordant with the breakpoints among the genetic clusters within the *Pseudopipra* complex. The distributions of *C. erythrocephala* and *rubrocapilla* are extensively codistributed with the Guianan Shield and Southern Amazonian clades of *Pseudopipra*. *C. rubrocapilla* has a range broadly overlapping the southern Amazon and the Atlantic Forest clades of *Pseudopipra*. *C. mentalis* is regionally codistributed with *Pseudopipra* in Central America but extends farther south into the Chocó and the western edges of Columbia and Ecuador. *C. mentalis* is also found at lower altitudes than the lower montane populations of *Pseudopipra* in Central America. *C. chloromeros* has a narrow distribution in the lower montane forests of the southern Peruvian and northern Bolivia Andes, while *Pseudpipira* has more extensive montane populations in the Andes from Peru to Colombia. *C. cornuta* is distributed in montane forests of tepuis in Venezuela and western Guyuna, at altitudes where *Pseudopipra* does not occur.

Considering these similarities and differences, the phylogenetic relationships among the differentiated lineages of *Ceratopipra* and *Pseudopipra* are at least partly congruent. For example, in *Ceratopipra,* the predominantly Central American *C. mentalis* is sister to a clade consisting of northern *C. erythrocephala* and southern *C. chloromeros* + *C. rubrocapilla* Amazonian lineages. However, the phylogenetic position of Guianan *cornuta* is not congruent with the deepest split in *Pseudopipra* representing a split in the Northern Andes.

Therefore, it seems probable that these taxa have components of their phylogeographic history in common and these patterns should be investigated in more detail.

#### Vocal variation

*Pseudopipra* vocalizations have 1-3 buzzy or tonal notes. We measured: 1) starting frequency, 2) ending frequency, 3) minimum frequency, 4) maximum frequency, 5) number of notes, and 6) duration of the entire vocalization (see Supplementary Figure 13). To obtain a conservative estimate of the number of individuals sampled, we took measurements of one vocalization from each recording. When there were multiple recordings by the same recordist on the same day and location, only one of the recordings was measured. Some recordists raised the possibility that the tonal notes, particularly the "tonk" in vocal type 1, may be a mechanical sound, but further research is required to determine which sounds are vocalizations and which are mechanical sonations. We performed principal components analysis (PCA) and logistic regression on the vocal measurements to test for significant differences between the vocal types and to reduce the dimensionality of the data for comparison to results from analysis of genetic data. The PCA analysis was performed using the princomp function and the logistic regression was performed using the glm function, both in the stats R package (R Core Team, 2018). The geographic distribution of each vocal type was assessed using latitude and longitude coordinates included in the metadata of each recording. When no coordinates were available, we determined latitude and longitude based on the description of the locality.

Our analysis identified 14 qualitatively distinct vocalization types that can be easily diagnosed by ear, or by visual inspection of sonograms (which we arbitrarily labeled numerically). Eleven types were identified as male advertisement songs on the basis of expert opinion in the context of recordist notes. Three additional types were classified as territorial calls, suggesting that most populations have a repertoire size of at least two vocalizations. We note that the behavioral context of these vocalizations was not always clear, so our assessment reflects a preliminary hypothesis of homology within these two categories.

##### *Male advertisement vocalizations*

All male advertisement vocalizations appear to be unique to a single monophyletic lineage (see Figure 8, Taxonomic Summary), except in the cases of vocalization types 2 (Clade F, C/E, and C6) and type 1 (A1 + B1) which appear to be shared plesiomorphically across non-sister lineages (see main text and discussion below). Type 4 (Clade A2), 3 (Clade D), and 7 (Clade C1) all appear restricted to single monophyletic lineages. In our initial assessment, we identified vocalization type 6, which has a geographic range which coincides with both the southern Tapajós clade and the Atlantic Forest clade. However, upon closer consideration, we found a subtle difference between the Atlantic Forest (6a) and Tapajós (6b) regions. Type 6a recordings, which are restricted to the Atlantic Forest, have a relatively constant pitch for both the buzz and tonal notes, whereas the type 6b recordings, which are restricted to the Tapajós regions, have a descending pitch for both notes. The sharing of a similar vocal phenotype (6a and 6b) clearly links the Atlantic forest populations to the Southern Amazon, in congruence with our genetic data (Clade C7), even though these southern Tapajós individuals have a genetic affinity to other more northern Tapajós individuals.

Lastly, we also identified several male advertisement vocalization types with restricted, non-overlapping geographic ranges in regions for which we lack genetic samples. Vocal type 8 was restricted to Subtropical forests of the eastern slope of the Andes from western Venezuela to Morona-Santiago, Ecuador. Vocal type 9 was restricted to Subtropical forests of western Cauca, Colombia south to Esmeraldas, Ecuador. Vocal types 11a and 11b were restricted to Subtropical forests of Magdalena Valley, Antioquia to Huila, Colombia. It is likely they fall within the range of variation delimited by the Central America + North Andean Peru clade.

*Call vocalizations*

Our analysis identified three distinct but similar call vocalization types that can be diagnosed by ear, or by visual inspection of sonograms. Vocalization type 5 was primarily recorded in the northern Amazon basin, coinciding with the broad ranging Northern Amazon + Guiana Shield lineage (Clade D). From our initial classification, one recording from the Atlantic Forest (XC427315) was assigned to type 5, suggesting this type may be shared between the Guiana Shield clade and the Atlantic Forest clade. If true, this pattern would imply that the southern Amazon clade may have two call types (13 and 5) – as this was a singular record, we choose to wait for additional confirmation before making this recommendation. As with vocalization type 1, call vocalization type 10 appears to be shared pleisiomorphically across deeply divergent non-sister Andean clades (A1 + B1). Lastly, call vocalization type 13 was found to be shared plesiomorphically across diverse lowland southern Amazonian forms (Clade C).

*Phylogenetic reconstruction of vocal patterns*

After assigning vocal types to genetic lineages, we generated a *preliminary* reconstruction of vocal type evolution (coded as discrete characters) under maximum parsimony, with the castor R package (Louca and Doebeli, 2017). For this analysis, we generated a lineage tree by hand to reflect the best supported phylogenetic hypothesis (though not considering branch lengths). As mentioned previously, given the behavioral context of vocalizations in manakins, which evolve under strong sexual selection (Prum, 1992, 1994), it is not clear how well our classification into lekking advertisements and call type categories represent homologous characters. While diagnostic for lineages, which is critical for identifying and recognizing species, the labels we have assigned to vocal types may represent either shared, derived, or convergent characters, even when the "same" vocal type we identify is shared across non-sister lineages (e.g, vocal type 2 for *pygmaea*, *discolor*, and *separabilis*). Without a better understanding of vocal character evolution, we present these reconstructions as preliminary and as a starting point for future studies.

In sum, and as depicted in Supplementary Figure 16, the *Pseudopipra* MRCA is reconstructed to have an Andean vocal type in both cases (male advertisement type 1, and call type 10). As noted above, and in the main text, vocal type 2 is also reconstructed as being shared plesiomorphically across a number of lowland Amazonian regions, with four vocal types (6a, 6b, 7, and 3) derived from this type. In the northern Andes and Central America, five male advertisement types (11a+11b, 11b, 9, 8, and 4) appear to be derived from Andean type 1. Despite the caveats listed above, we find this analysis to be compelling because the patterns we detect under parsimony are consistent with the narrative we develop in the main text about the likely Andean origins of the clade, both for call type vocalizations and male advertisement vocalizations.

### Supplementary Figures and Tables

**Supplementary Table 1. Specimen data table (separate xlsx)**

**Supplementary Table 2a. Results from Mantel tests – patristic distance**

Supplementary Table 2a.

| Region | locality code | mantel r | two-tailed p | lower 2.5% limit | upper 97.5% limit | log.d | perm | significant (p < 0.001) |
|---|---|---|---|---|---|---|---|---|
| Full dataset | | 0.748 | 0.0001 | 0.729 | 0.763 | F | 10000 | * |
| Central America - Costa Rica | CACR | 0.459 | 0.2551 | 0.017 | 1.000 | F | 10000 | - |
| Central America - Panama | CAPA | 0.694 | 0.1019 | 0.021 | 0.921 | F | 10000 | - |
| North Andean – San Martín | CAMA | NA | NA | NA | NA | F | 10000 | - |
| South Andean Peru (North) | CPN | -0.261 | 1 | -0.261 | -0.261 | F | 10000 | - |
| South Andean Peru (South) | CPS | 0.816 | 0.0993 | 0.618 | 0.998 | F | 10000 | - |
| Weakly resolved Guiana Shield | GS | 0.099 | 0.0947 | 0.058 | 0.134 | F | 10000 | - |
| Unresolved Jaú | GSIMERI | 0.096 | 0.243 | 0.043 | 0.166 | F | 10000 | - |
| Weakly resolved eastern Napo | GSNAPO | 0.313 | 0.0513 | 0.177 | 0.560 | F | 10000 | - |
| Weakly resolved Suriname + Amapá | GSSR | 0.069 | 0.2483 | 0.032 | 0.121 | F | 10000 | - |
| Western Napo introgressed lineage | PH | 0.508 | 0.017 | 0.340 | 0.754 | F | 10000 | - |
| Western Inambari endemic | INAMBARI | 0.608 | 0.001 | 0.382 | 0.848 | F | 10000 | * |
| Eastern Inambari endemic | INAMBARIE | 0.004 | 0.981 | -0.196 | 0.244 | F | 10000 | - |
| Rondônia endemic | RONDONIA | NA | NA | NA | NA | F | 10000 | - |
| Tapajós endemic | TAPAJOS | 0.250 | 0.0148 | 0.182 | 0.313 | F | 10000 | - |
| Xingu endemic | XINGU | 0.079 | 0.8572 | -0.170 | 0.275 | F | 10000 | - |
| Atlantic Forest – Bahia | AFBAHIA | 0.484 | 0.0173 | 0.173 | 0.804 | F | 10000 | - |
| Atlantic Forest – Espírito Santo | AFES | NA | NA | NA | NA | F | 10000 | - |
| Atlantic Forest – Rio | AFRIO | 0.701 | 0.0007 | 0.615 | 0.782 | F | 10000 | * |

| Region | locality code | mantel r | two-tailed p | lower 2.5% limit | upper 97.5% limit | log.d | perm | significant (p < 0.001) |
|---|---|---|---|---|---|---|---|---|
| Full dataset | | 0.391 | 0.0001 | 0.379 | 0.401 | T | 10000 | * |
| Central America - Costa Rica | CACR | 0.459 | 0.253 | 0.017 | 1.000 | T | 10000 | - |
| Central America - Panama | CAPA | 0.519 | 0.2532 | 0.021 | 0.919 | T | 10000 | - |
| North Andean – San Martín | CAMA | NA | NA | NA | NA | T | 10000 | - |
| South Andean Peru (North) | CPN | -0.261 | 1 | -0.261 | -0.261 | T | 10000 | - |
| South Andean Peru (South) | CPS | 0.810 | 0.032 | 0.669 | 0.998 | T | 10000 | - |
| Weakly resolved Guiana Shield | GS | 0.091 | 0.0022 | 0.063 | 0.119 | T | 10000 | - |
| Unresolved Jaú | GSIMERI | 0.536 | 0.0001 | 0.319 | 0.684 | T | 10000 | * |
| Weakly resolved eastern Napo | GSNAPO | 0.313 | 0.0479 | 0.161 | 0.574 | T | 10000 | - |
| Weakly resolved Suriname + Amapá | GSSR | 0.065 | 0.1054 | 0.026 | 0.117 | T | 10000 | - |
| Western Napo introgressed lineage | PH | 0.248 | 0.091 | 0.102 | 0.407 | T | 10000 | - |
| Western Inambari endemic | INAMBARI | 0.714 | 0.0008 | 0.499 | 0.952 | T | 10000 | * |
| Eastern Inambari endemic | INAMBARIE | -0.004 | 0.9801 | -0.304 | 0.213 | T | 10000 | - |
| Rondônia endemic | RONDONIA | NA | NA | NA | NA | T | 10000 | - |
| Tapajós endemic | TAPAJOS | 0.247 | 0.0011 | 0.115 | 0.353 | T | 10000 | - |
| Xingu endemic | XINGU | 0.079 | 0.854 | -0.112 | 0.275 | T | 10000 | - |
| Atlantic Forest – Bahia | AFBAHIA | 0.481 | 0.0693 | 0.298 | 0.885 | T | 10000 | - |
| Atlantic Forest – Espírito Santo | AFES | NA | NA | NA | NA | T | 10000 | - |
| Atlantic Forest – Rio | AFRIO | 0.747 | 0.0012 | 0.676 | 0.912 | T | 10000 | - |

**Supplementary Table 2b. Results from partial Mantel tests – patristic distance**

Supplementary Table 2b.

| Approximate Barrier | Comparison (populations) | partial mantel r | two-tailed p | lower 2.5% limit | upper 97.5% limit | log.d | perm | significant (p < 0.001) |
|---|---|---|---|---|---|---|---|---|
| Cordillera de Talamanca | Costa Rica vs Panama | -0.1586 | 0.2972 | -0.2668 | 0.1548 | F | 10000 | - |
| Andes (1) | Central America vs Marañón | -0.5912 | 0.0967 | -0.7292 | -0.4338 | F | 10000 | - |
| Andes (2) | Central America vs (Marañón + South Andean Peru) | 0.0410 | 0.4030 | -0.2477 | 0.1369 | F | 10000 | - |
| Andes (3) | Central America vs (Everything else) | -0.2671 | 0.0001 | -0.2948 | -0.2293 | F | 10000 | * |
| Ucayali river | South Andean Peru vs Inambari | -0.9065 | 0.0001 | -0.9403 | -0.8721 | F | 10000 | * |
| Eastern Marañón + Hauallaga Rivers | Introgressed western Napo vs Inambari | -0.8963 | 0.0001 | -0.9166 | -0.8776 | F | 10000 | * |
| Putumayo river | Introgressed western Napo vs eastern Napo | -0.7442 | 0.0001 | -0.8258 | -0.7020 | F | 10000 | * |
| Purus river | Western Inambari vs eastern Inambari | -0.6034 | 0.0001 | -0.7205 | -0.0439 | F | 10000 | * |
| Madeira river | Eastern Inambari vs Rondo̒nia | -0.5498 | 0.0001 | -0.6974 | -0.1289 | F | 10000 | * |
| Tapajós river | Rondo̒nia vs Tapajós | -0.1751 | 0.0441 | -0.2328 | -0.1138 | F | 10000 | - |
| Xingu river | Tapajós vs Xingu | -0.3176 | 0.0004 | -0.3835 | -0.2400 | F | 10000 | * |
| Cerrado (1) | All pooled pops vs pooled Atlantic Forest | -0.1834 | 0.0008 | -0.2286 | -0.1122 | F | 10000 | * |
| Cerrado (2) | Xingu vs Bahia | -0.5647 | 0.0001 | -0.6092 | -0.4341 | F | 10000 | * |
| Japurá river | Eastern Napo vs Jaú | -0.1507 | 0.0146 | -0.2969 | -0.1086 | F | 10000 | - |
| Negro river | Jaú vs central Guiana Shield | -0.0530 | 0.3107 | -0.0880 | -0.0120 | F | 10000 | - |
| Essequibo river | central Guiana Shield vs eastern Guiana shield | -0.3502 | 0.0001 | -0.3913 | 0.0600 | F | 10000 | * |
| Amazonas river | All pooled lowland N vs all pooled lowland S | -0.9118 | 0.0001 | -0.9182 | -0.9071 | F | 10000 | * |

| Approximate Barrier | Comparison (populations) | partial mantel r | two-tailed p | lower 2.5% limit | upper 97.5% limit | log.d | perm | significant (p < 0.001) |
|---|---|---|---|---|---|---|---|---|
| Cordillera de Talamanca | Costa Rica vs Panama | -0.7445 | 0.0001 | -0.8021 | -0.6927 | T | 10000 | * |
| Andes (1) | Central America vs Marañón | -0.9955 | 0.0001 | -0.9974 | -0.9633 | T | 10000 | * |
| Andes (2) | Central America vs (Marañón + South Andean Peru) | -0.8292 | 0.0001 | -0.9416 | -0.7805 | T | 10000 | * |
| Andes (3) | Central America vs (Everything else) | -0.4636 | 0.0001 | -0.4952 | -0.4196 | T | 10000 | * |
| Ucayali river | South Andean Peru vs Inambari | -0.9096 | 0.0001 | -0.9382 | -0.8828 | T | 10000 | * |
| Eastern Marañón + Hauallaga Rivers | Introgressed western Napo vs Inambari | -0.8800 | 0.0001 | -0.9005 | -0.8622 | T | 10000 | * |
| Putumayo river | Introgressed western Napo vs eastern Napo | -0.9000 | 0.0001 | -0.9407 | -0.8842 | T | 10000 | * |
| Purus river | Western Inambari vs eastern Inambari | -0.8332 | 0.0001 | -0.8629 | -0.8110 | T | 10000 | * |
| Madeira river | Eastern Inambari vs Rondo̒nia | -0.7513 | 0.0001 | -0.8139 | -0.6498 | T | 10000 | * |
| Tapajós river | Rondo̒nia vs Tapajós | -0.3161 | 0.0001 | -0.3879 | -0.2109 | T | 10000 | * |
| Xingu river | Tapajós vs Xingu | -0.7034 | 0.0001 | -0.7543 | -0.6546 | T | 10000 | * |
| Cerrado (1) | All pooled pops vs pooled Atlantic Forest | -0.7213 | 0.0001 | -0.7450 | -0.7020 | T | 10000 | * |
| Cerrado (2) | Xingu vs Bahia | -0.9339 | 0.0001 | -0.9517 | 0.0700 | T | 10000 | * |
| Japurá river | Eastern Napo vs Jaú | -0.3993 | 0.0001 | -0.4896 | -0.3293 | T | 10000 | * |
| Negro river | Jaú vs central Guiana Shield | -0.0360 | 0.4825 | -0.0690 | 0.0010 | T | 10000 | - |
| Essequibo river | central Guiana Shield vs eastern Guiana shield | -0.4704 | 0.0001 | -0.5026 | -0.4437 | T | 10000 | * |
| Amazonas river | All pooled lowland N vs all pooled lowland S | -0.9150 | 0.0001 | -0.9205 | -0.9106 | T | 10000 | * |

**Supplementary Table 2c. Results from Mantel tests – coancestry**

Supplementary Table 2c.

| **Region** | locality code | **mantel r** | **two-tailed p** | **lower 2.5% limit** | **upper 97.5% limit** | **log.d** | **perm** | **significant (p < 0.001)** |
|---|---|---|---|---|---|---|---|---|
| Full dataset | | 0.3735 | 0.0001 | 0.3598 | 0.3879 | F | 10000 | * |
| Central America - Costa Rica | CACR | -0.4803 | 0.7496 | -0.8288 | 0.0004 | F | 10000 | - |
| Central America - Panama | CAPA | 0.3246 | 0.4483 | -0.0001 | 0.9743 | F | 10000 | - |
| North Andean – San Martín | CAMA | NA | NA | NA | NA | F | 10000 | - |
| South Andean Peru (North) | CPN | 0.8087 | 0.6701 | 0.8087 | 0.8087 | F | 10000 | - |
| South Andean Peru (South) | CPS | 0.4880 | 0.0987 | -0.6721 | 0.9726 | F | 10000 | - |
| Weakly resolved Guiana Shield | GS | 0.0691 | 0.0010 | 0.0561 | 0.0862 | F | 10000 | - |
| Unresolved Jaú | GSIMERI | 0.1475 | 0.1410 | 0.0506 | 0.2401 | F | 10000 | - |
| Weakly resolved eastern Napo | GSNAPO | 0.0996 | 0.4049 | -0.0868 | 0.2369 | F | 10000 | - |
| Weakly resolved Suriname + Amapá | GSSR | 0.0987 | 0.0269 | 0.0592 | 0.1262 | F | 10000 | - |
| Western Napo introgressed lineage | PH | 0.2729 | 0.0802 | 0.1492 | 0.4670 | F | 10000 | - |
| Western Inambari endemic | INAMBARI | 0.4788 | 0.0047 | 0.2834 | 0.6791 | F | 10000 | - |
| Eastern Inambari endemic | INAMBARIE | 0.0160 | 0.9285 | -0.2173 | 0.2406 | F | 10000 | - |
| Rondônia endemic | RONDONIA | NA | NA | NA | NA | F | 10000 | - |
| Tapajós endemic | TAPAJOS | 0.0999 | 0.0415 | 0.0639 | 0.1461 | F | 10000 | - |
| Xingu endemic | XINGU | 0.1910 | 0.5621 | -0.3141 | 0.5524 | F | 10000 | - |
| Atlantic Forest – Bahia | AFBAHIA | -0.0081 | 1.0000 | -0.2826 | 0.3544 | F | 10000 | - |
| Atlantic Forest – Espírito Santo | AFES | NA | NA | NA | NA | F | 10000 | - |
| Atlantic Forest – Rio | AFRIO | 0.6050 | 0.0005 | 0.5585 | 0.7021 | F | 10000 | * |

| **Region** | locality code | **mantel r** | **two-tailed p** | **lower 2.5% limit** | **upper 97.5% limit** | **log.d** | **perm** | **significant (p < 0.001)** |
|---|---|---|---|---|---|---|---|---|
| Full dataset | | 0.3262 | 0.0001 | 0.3133 | 0.3378 | T | 10000 | * |
| Central America - Costa Rica | CACR | -0.4803 | 0.7437 | -0.8288 | 0.0004 | T | 10000 | - |
| Central America - Panama | CAPA | 0.7348 | 0.1027 | 0.2647 | 0.9485 | T | 10000 | - |
| North Andean – San Martín | CAMA | NA | NA | NA | NA | T | 10000 | - |
| South Andean Peru (North) | CPN | 0.8087 | 0.6704 | 0.8087 | 0.8087 | T | 10000 | - |
| South Andean Peru (South) | CPS | 0.5794 | 0.1639 | -0.6546 | 0.9644 | T | 10000 | - |
| Weakly resolved Guiana Shield | GS | 0.1601 | 0.0001 | 0.1282 | 0.1836 | T | 10000 | * |
| Unresolved Jaú | GSIMERI | 0.4496 | 0.0005 | 0.2383 | 0.5880 | T | 10000 | * |
| Weakly resolved eastern Napo | GSNAPO | 0.0996 | 0.4055 | -0.0839 | 0.2374 | T | 10000 | - |
| Weakly resolved Suriname + Amapá | GSSR | 0.1860 | 0.0001 | 0.1408 | 0.2226 | T | 10000 | * |
| Western Napo introgressed lineage | PH | 0.2832 | 0.0566 | 0.1543 | 0.4336 | T | 10000 | - |
| Western Inambari endemic | INAMBARI | 0.4351 | 0.0283 | 0.1358 | 0.8800 | T | 10000 | * |
| Eastern Inambari endemic | INAMBARIE | 0.2755 | 0.0736 | 0.0514 | 0.4345 | T | 10000 | - |
| Rondônia endemic | RONDONIA | NA | NA | NA | NA | T | 10000 | - |
| Tapajós endemic | TAPAJOS | 0.3406 | 0.0003 | 0.1647 | 0.4277 | T | 10000 | * |
| Xingu endemic | XINGU | 0.1910 | 0.5648 | -0.3034 | 0.5639 | T | 10000 | - |
| Atlantic Forest – Bahia | AFBAHIA | -0.0217 | 0.9040 | -0.3916 | 0.4473 | T | 10000 | - |
| Atlantic Forest – Espírito Santo | AFES | NA | NA | NA | NA | T | 10000 | - |
| Atlantic Forest – Rio | AFRIO | 0.5455 | 0.0006 | 0.4761 | 0.6276 | T | 10000 | * |

**Supplementary Table 2d. Results from partial Mantel tests – coancestry**

Supplementary Table 2d.

| Approximate Barrier | Comparison (populations) | partial mantel r | two-tailed p | lower 2.5% limit | upper 97.5% limit | log.d | perm | significant (p < 0.001) |
|---|---|---|---|---|---|---|---|---|
| Cordillera de Talamanca | Costa Rica vs Panama | -0.5524 | 0.0073 | -0.6754 | 0.0921 | F | 10000 | - |
| Andes (1) | Central America vs Marañón | -0.4747 | 0.0967 | -0.6474 | -0.3129 | F | 10000 | - |
| Andes (2) | Central America vs (Marañón + South Andean Peru) | 0.0858 | 0.0449 | -0.1128 | 0.1744 | F | 10000 | - |
| Andes (3) | Central America vs (Everything else) | 1.0000 | 0.0514 | 0.0299 | 0.0466 | F | 10000 | - |
| Ucayali river | South Andean Peru vs Inambari | -0.6470 | 0.0003 | -0.7328 | -0.5724 | F | 10000 | * |
| Eastern Marañón + Hauallaga Rivers | Introgressed western Napo vs Inambari | -0.7325 | 0.0002 | -0.7892 | -0.6733 | F | 10000 | * |
| Putumayo river | Introgressed western Napo vs eastern Napo | -0.5847 | 0.0001 | -0.6282 | -0.5427 | F | 10000 | * |
| Purus river | Western Inambari vs eastern Inambari | -0.2668 | 0.0016 | -0.3713 | -0.1015 | F | 10000 | - |
| Madeira river | Eastern Inambari vs Rondo̐nia | -0.2351 | 0.2674 | -0.4326 | -0.0358 | F | 10000 | * |
| Tapajós river | Rondo̐nia vs Tapajós | -0.1295 | 0.0310 | -0.4121 | -0.0761 | F | 10000 | - |
| Xingu river | Tapajós vs Xingu | -0.0328 | 0.5455 | -0.0953 | 0.0045 | F | 10000 | - |
| Cerrado (1) | All pooled pops vs pooled Atlantic Forest | 0.1522 | 0.0088 | 0.1258 | 0.1782 | F | 10000 | - |
| Cerrado (2) | Xingu vs Bahia | -0.6525 | 0.0009 | -0.7186 | -0.5138 | F | 10000 | * |
| Japurá river | Eastern Napo vs Jaú | -0.2413 | 0.0001 | -0.3151 | -0.1326 | F | 10000 | * |
| Negro river | Jaú vs central Guiana Shield | 0.0029 | 0.8820 | -0.0108 | 0.0218 | F | 10000 | - |
| Essequibo river | central Guiana Shield vs eastern Guiana shield | -0.1383 | 0.0001 | -0.1815 | -0.1164 | F | 10000 | * |
| Amazonas river | All pooled lowland N vs all pooled lowland S | -0.8101 | 0.0001 | -0.8289 | -0.7992 | F | 10000 | * |

| Approximate Barrier | Comparison (populations) | partial mantel r | two-tailed p | lower 2.5% limit | upper 97.5% limit | log.d | perm | significant (p < 0.001) |
|---|---|---|---|---|---|---|---|---|
| Cordillera de Talamanca | Costa Rica vs Panama | -0.3805 | 0.0139 | -0.6218 | 0.0465 | T | 10000 | - |
| Andes (1) | Central America vs Marañón | -0.9876 | 0.0001 | -0.9962 | -0.9080 | T | 10000 | * |
| Andes (2) | Central America vs (Marañón + South Andean Peru) | -0.7704 | 0.0001 | -0.9038 | -0.7109 | T | 10000 | * |
| Andes (3) | Central America vs (Everything else) | -0.0855 | 0.0001 | -0.1011 | -0.0740 | T | 10000 | * |
| Ucayali river | South Andean Peru vs Inambari | -0.6449 | 0.0001 | -0.7262 | -0.5683 | T | 10000 | * |
| Eastern Marañón + Hauallaga Rivers | Introgressed western Napo vs Inambari | -0.7017 | 0.0001 | -0.7649 | -0.6387 | T | 10000 | * |
| Putumayo river | Introgressed western Napo vs eastern Napo | -0.7874 | 0.0001 | -0.8147 | -0.7670 | T | 10000 | * |
| Purus river | Western Inambari vs eastern Inambari | 0.9000 | 0.0001 | -0.7288 | -0.5743 | T | 10000 | * |
| Madeira river | Eastern Inambari vs Rondo̐nia | -0.6487 | 0.0167 | -0.7388 | -0.5370 | T | 10000 | - |
| Tapajós river | Rondo̐nia vs Tapajós | -0.1451 | 0.0038 | -0.3839 | -0.0801 | T | 10000 | - |
| Xingu river | Tapajós vs Xingu | -0.1200 | 0.0017 | -0.2963 | -0.0685 | T | 10000 | - |
| Cerrado (1) | All pooled pops vs pooled Atlantic Forest | -0.2015 | 0.0007 | -0.2347 | -0.1710 | T | 10000 | * |
| Cerrado (2) | Xingu vs Bahia | -0.7871 | 0.0001 | -0.8905 | -0.7102 | T | 10000 | * |
| Japurá river | Eastern Napo vs Jaú | -0.4626 | 0.0001 | -0.5232 | -0.4126 | T | 10000 | * |
| Negro river | Jaú vs central Guiana Shield | 0.0322 | 0.9000 | 0.0217 | 0.0482 | T | 10000 | - |
| Essequibo river | central Guiana Shield vs eastern Guiana shield | -0.2212 | 0.0001 | -0.2821 | -0.1912 | T | 10000 | * |
| Amazonas river | All pooled lowland N vs all pooled lowland S | -0.8197 | 0.0001 | -0.8358 | -0.8092 | T | 10000 | * |

**Supplementary Table 3: Vocal measures (separate xlsx)**

**Supplementary Table 4: Vocal recording metadata (separate xlsx)**

**Supplementary Table 5: G-PhoCS parameters**

**Supplementary Table 5a, θ: effective population size**

| | Western Napo | Eastern Napo | Inambari | MRCA Western Napo, Inambari | MRCA Inambari, Eastern Napo |
|---|---|---|---|---|---|
| median | 4950840 | 2807467.5 | 2797062.5 | 5356300 | 1429540 |
| 95% HPD Interval LOW | 4527457.5 | 2498760 | 2566870 | 4658055 | 1352200 |
| 95% HPD Interval HIGH | 5378937.5 | 3132990 | 3044770 | 6047630 | 1512647.5 |

**Supplementary Table 5b, $\tau$: splitting time in generations**

| | MRCA Western Napo, Inambari | MRCA Inambari, Eastern Napo |
|---|---|---|
| median | 484600 | 1025420 |
| 95% HPD Interval LOW | 442610 | 954030 |
| 95% HPD Interval HIGH | 532120 | 1099450 |

**Supplementary Table 5c, m : migration rate (migrants per generation)**

| | Eastern Napo to Inambari | Inambari to Eastern Napo | Eastern Napo to Western Napo | Western Napo to Eastern Napo | Inambari to Western Napo | Western Napo to Inambari |
|---|---|---|---|---|---|---|
| median | 0.0663 | 0.4196 | 1.7508 | 0.3671 | 0.0000 | 0.0000 |
| 95% HPD Interval LOW | 0.0000 | 0.2670 | 1.0470 | 0.0000 | 0.0000 | 0.0000 |
| 95% HPD Interval HIGH | 0.1763 | 0.6109 | 2.5872 | 0.8184 | 0.0056 | 0.0028 |

**Supplementary Figure 1. GBIF occurrence records**
In this plot, we show all GBIF occurrence records in October 2018 (in black) with our genetic sampling localities in red. The BirdLife approximate genus range map is shown in green, and our modifications to this map are shown in. This figure is provided primarily to illustrate how the BirdLife range map is inaccurate in the western Amazon, in Loreto, Peru, where our analyses detect an introgressed hybrid lineage, as well as across the ranges of Andan taxa, which were revised after discussion with Andrés M. Cuervo (*personal communication*). We emphasize that the geographic limit boundaries depicted in map figures are approximate.

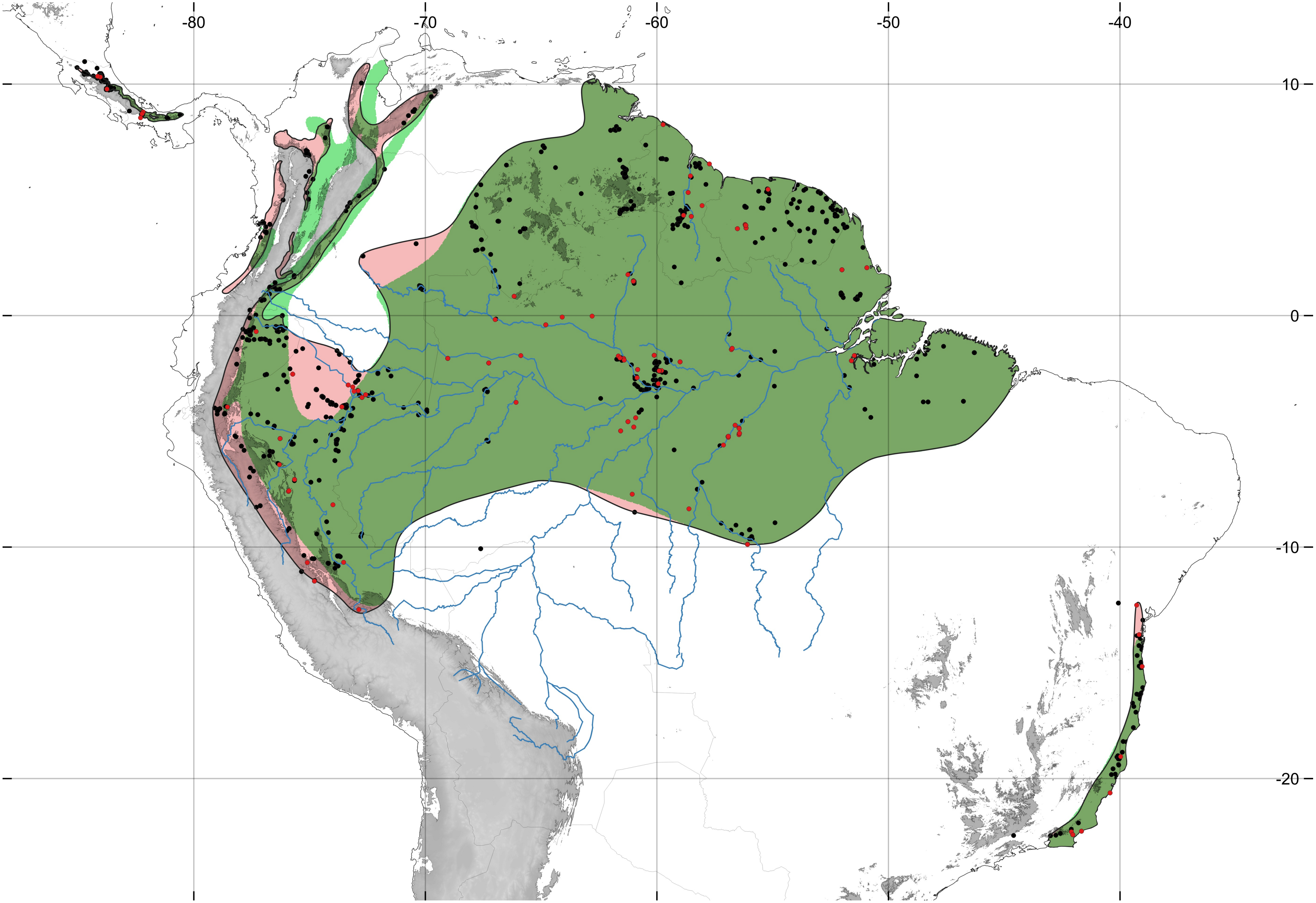

-80
-70
-60
-50
-40
10
0
-10
-20

**Supplementary Figure 2a-h. K-means clustering of SNP datasets**
For each figure a-h, the PCoA projection of SNP data is plotted on the top panel, with minimum convex hulls (minimum implied range) and plotting symbols indicating an optimized K-means clustering solution based on BIC score. PCoA explained 13-17% of the variance in the SNP data on the first two axes, and K-means clustering assignments derived from each dataset recovers similar population assignments (see main text for descriptions). Plotting symbols and colored convex hulls reflect cluster assignment.

### Supplementary Figure 2a

**dataset1 PCoA k−means K = 5**

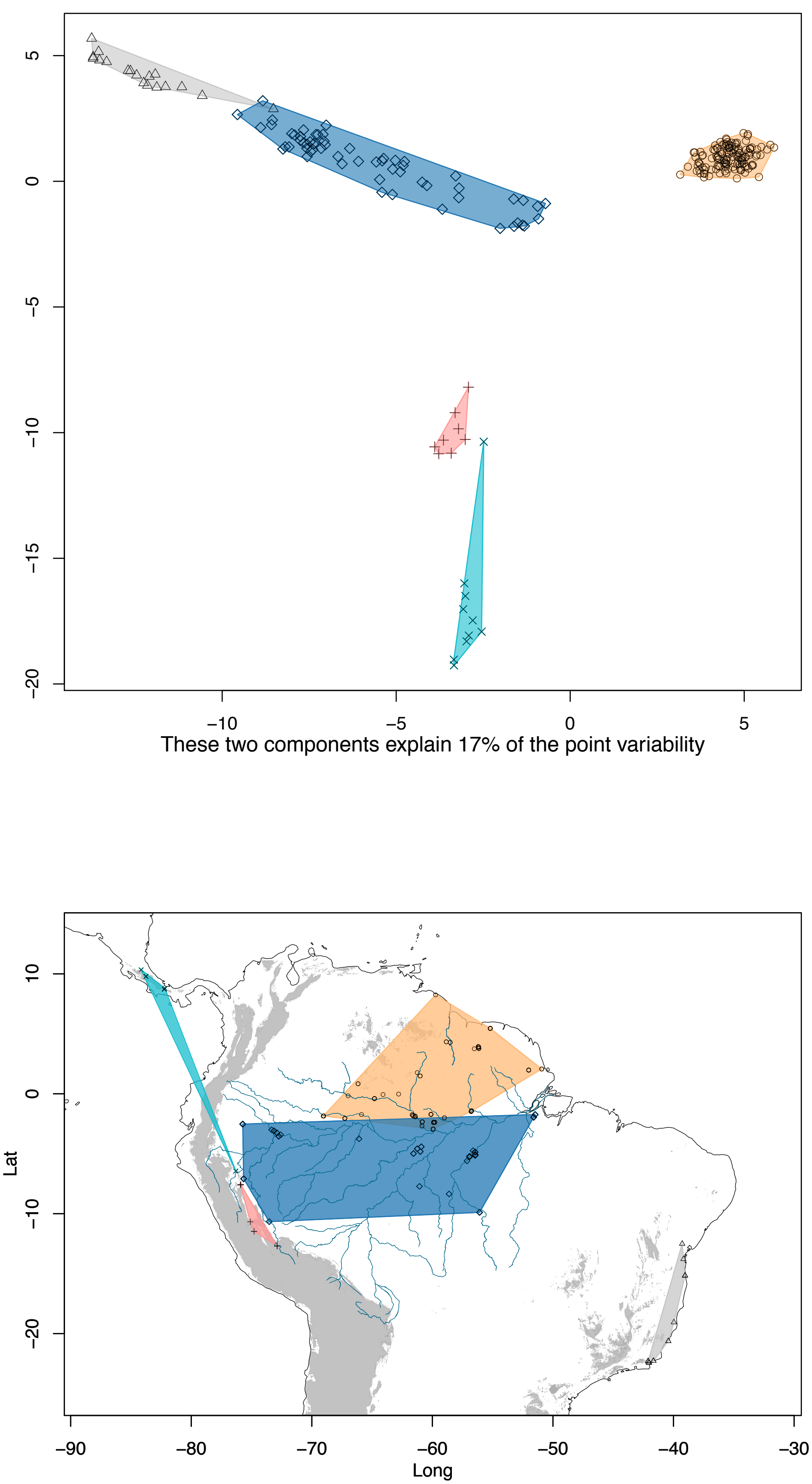

#### Supplementary Figure 2b

**dataset1.nomiss PCoA k−means K = 6**

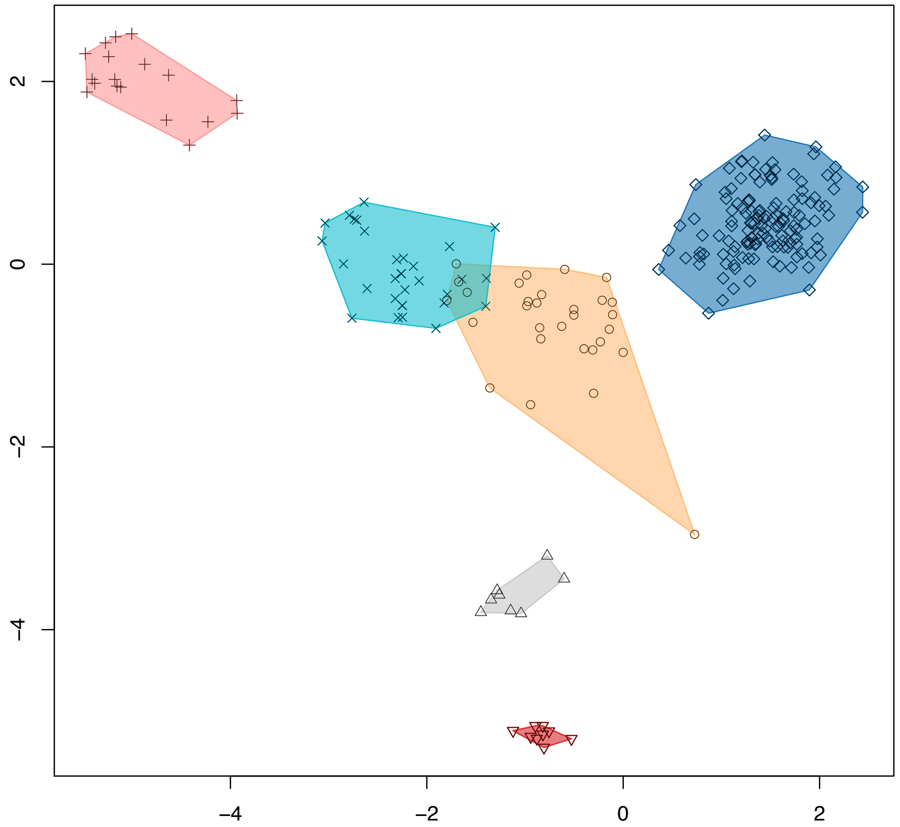

These two components explain 22% of the point variability

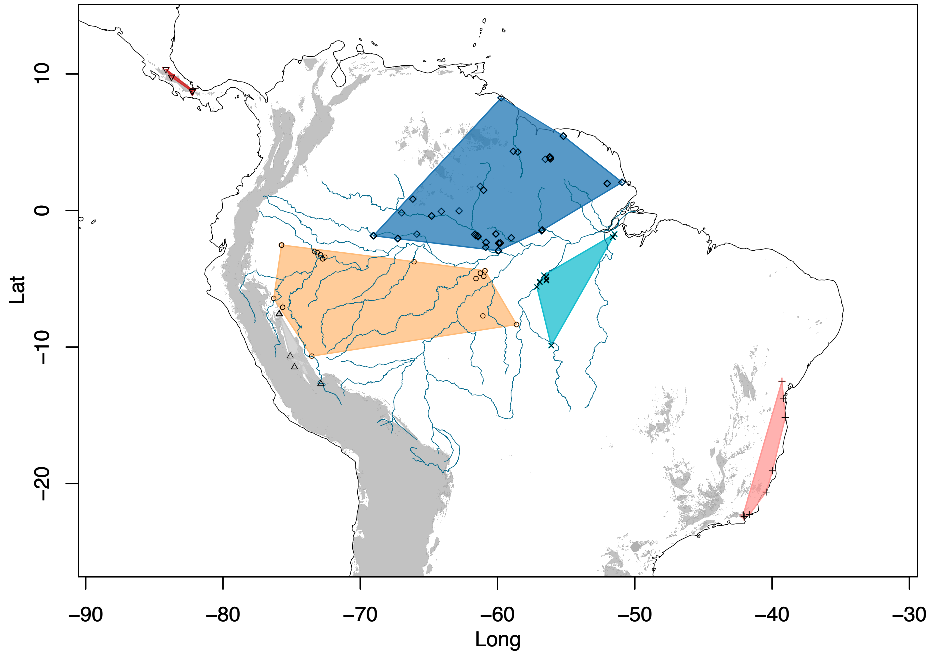

#### Supplementary Figure 2c

**dataset2 PCoA k–means K = 5**

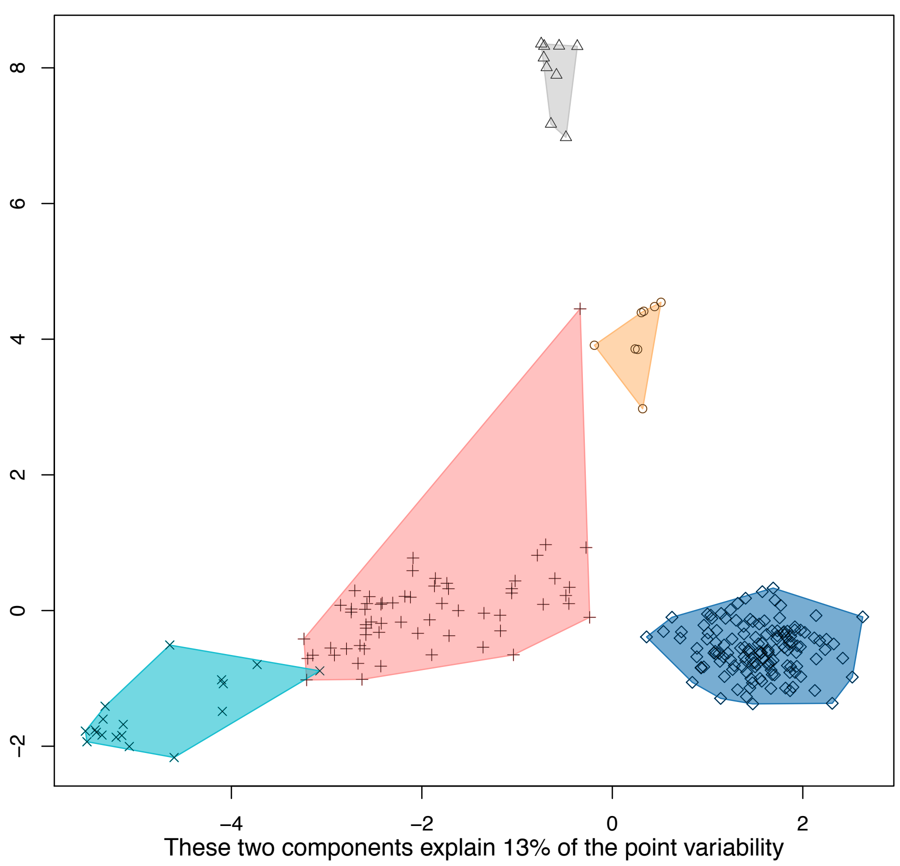

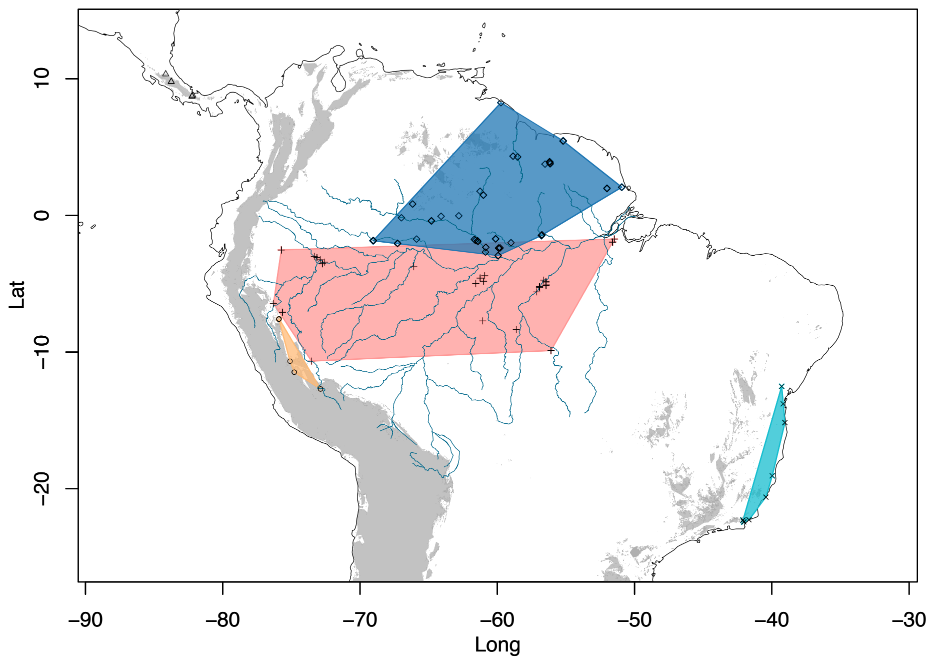

### Supplementary Figure 2d

**dataset2.nomiss PCoA k−means K = 8**

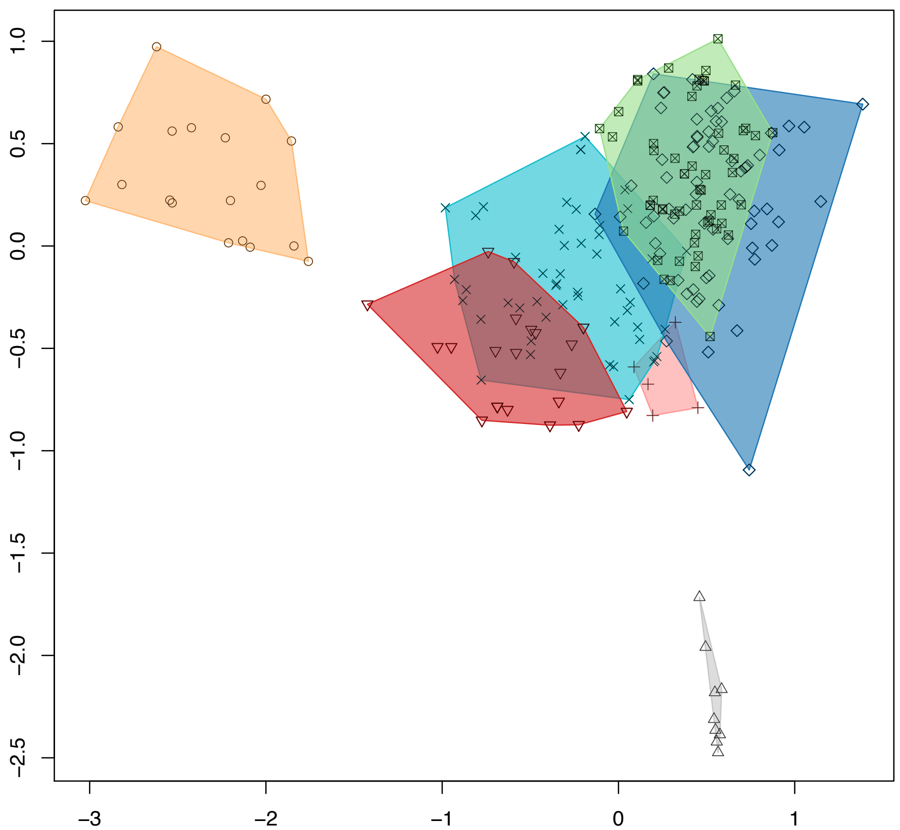

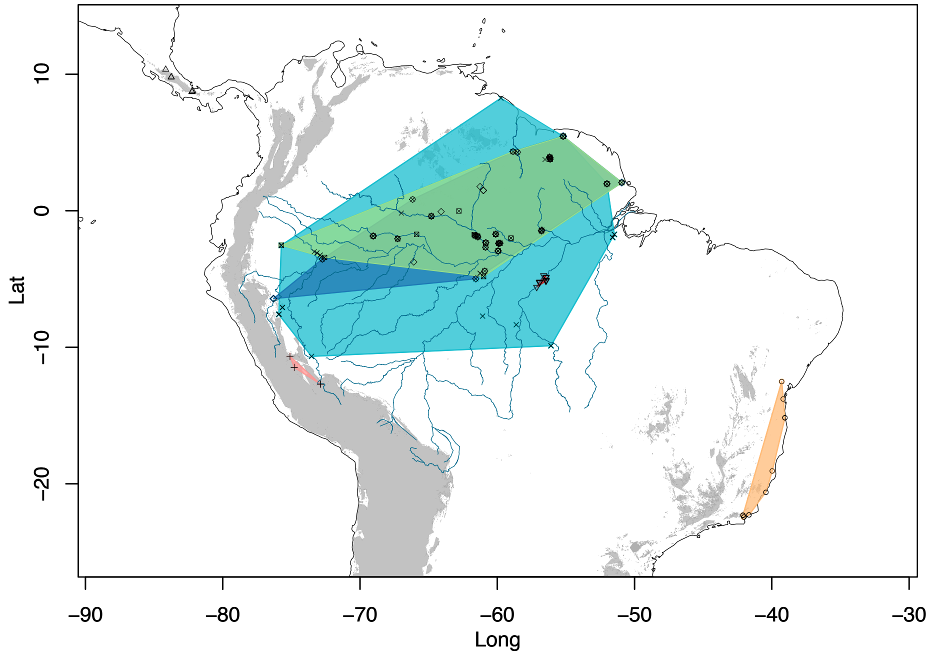

### Supplementary Figure 2e

**dataset3 PCoA k−means K = 5**

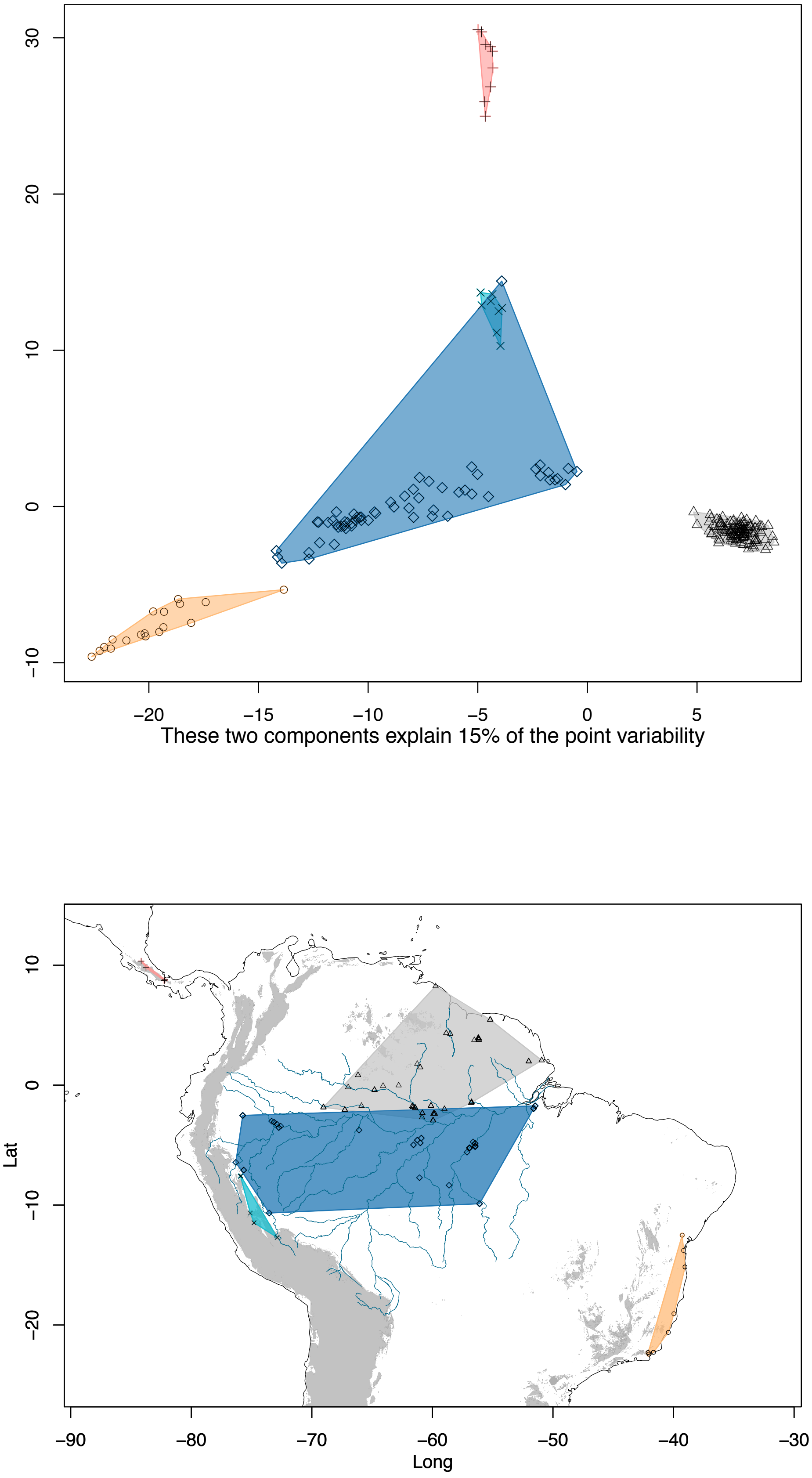

#### Supplementary Figure 2f

**dataset3.nomiss PCoA k−means K = 5**

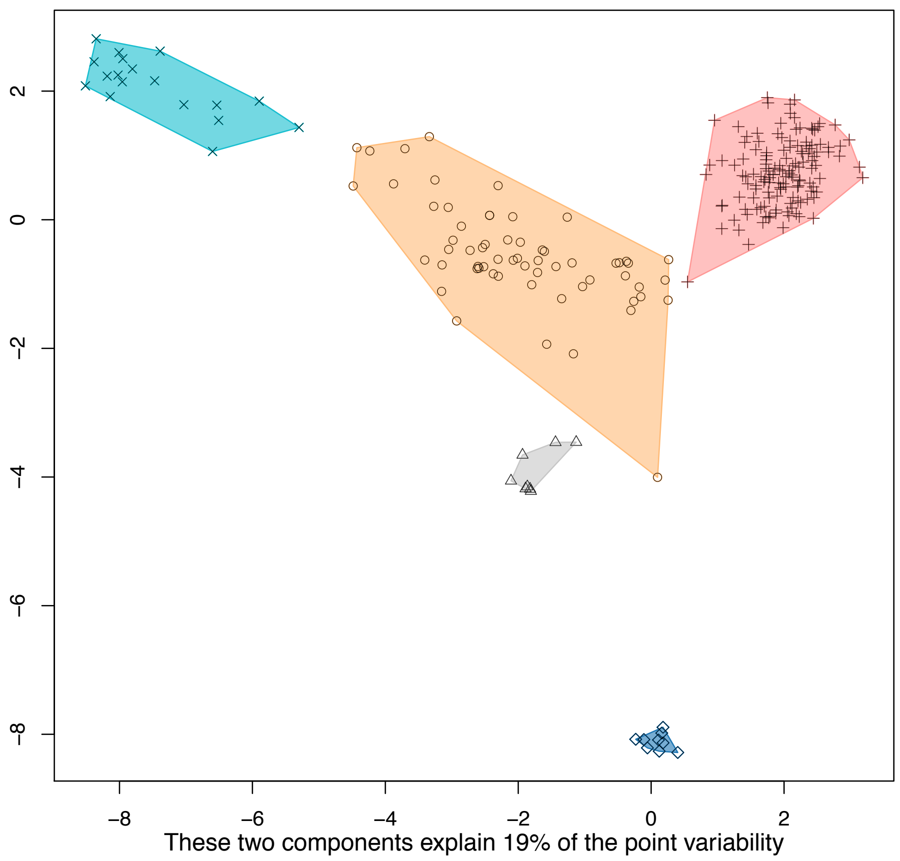

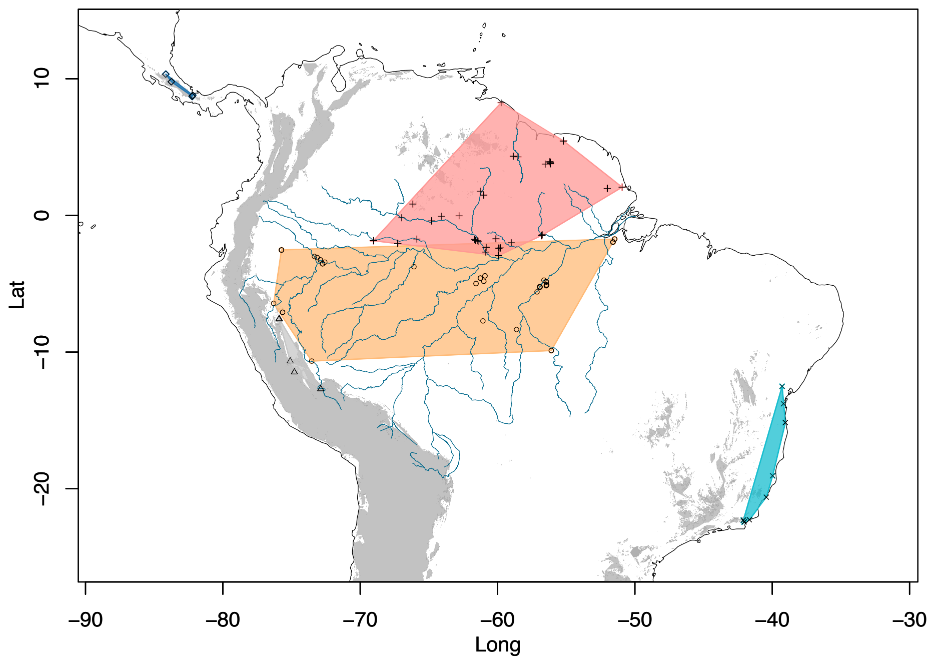

### Supplementary Figure 2g

**dataset4 PCoA k–means K = 5**

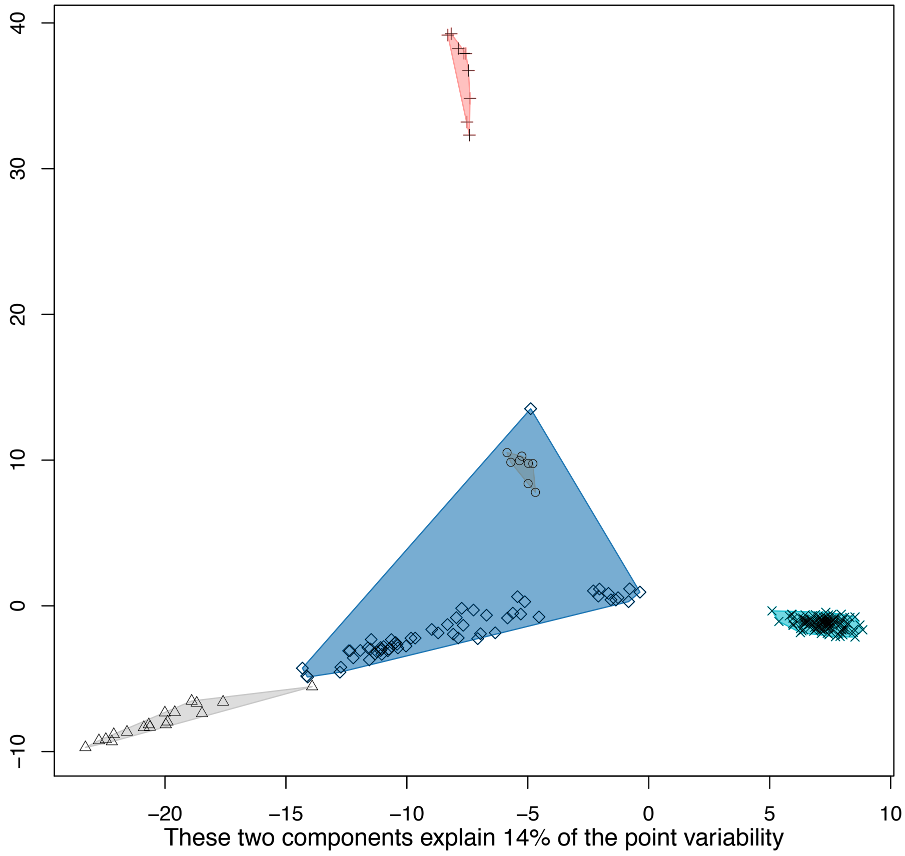

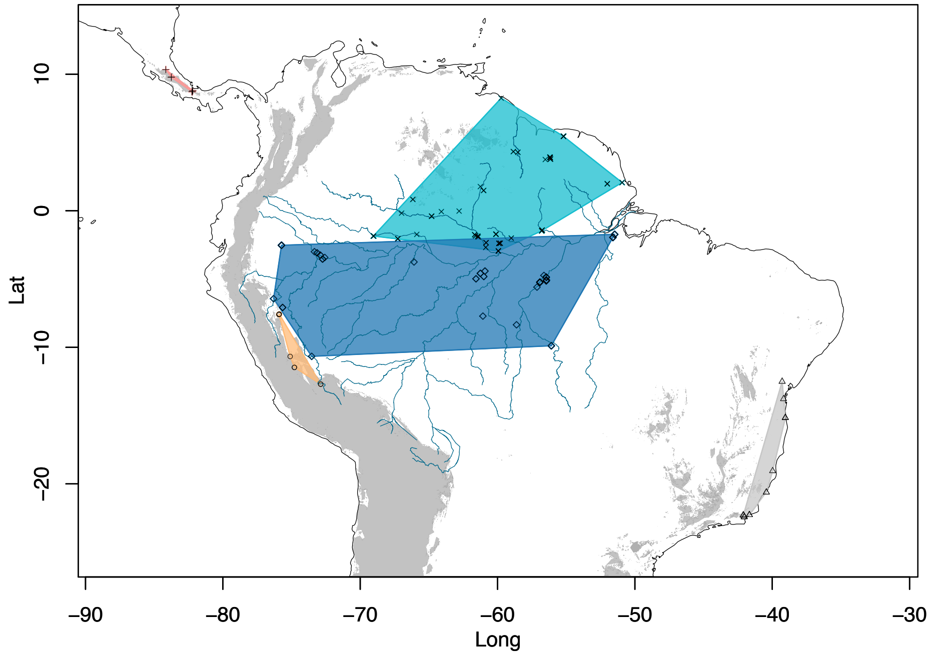

#### Supplementary Figure 2h

**dataset4.nomiss PCoA k−means K = 5**

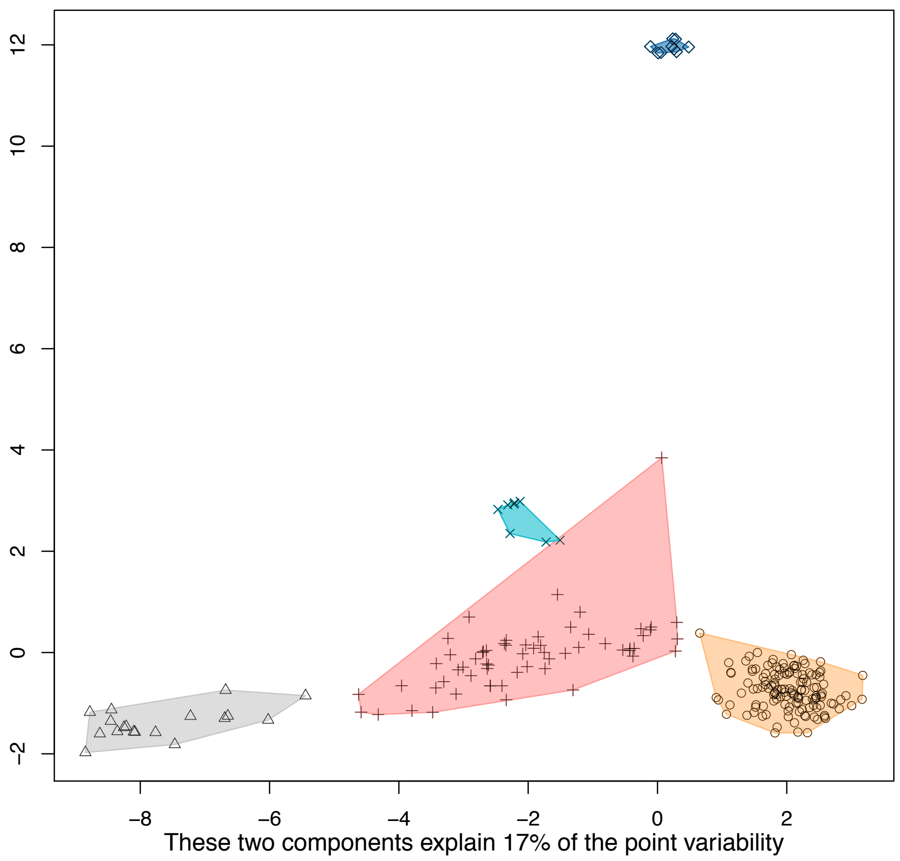

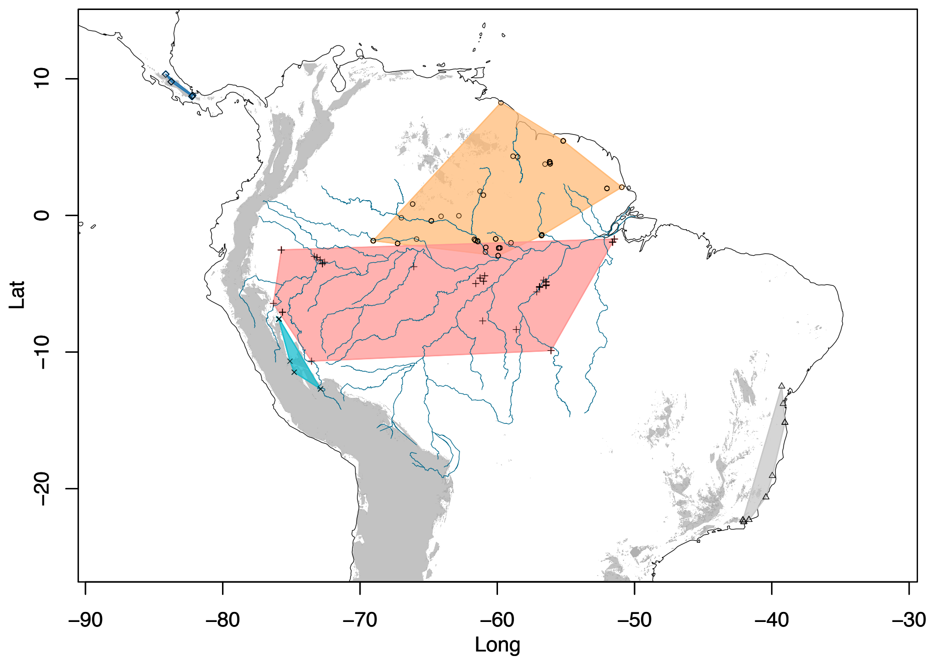

**Supplementary Figure 3 – log likelihoods of STRUCTURE runs**
Summarized log likelihood values across STRUCTURE runs for each value of K, with a plateau starting at K~5, and variance across runs increasing dramatically after K10

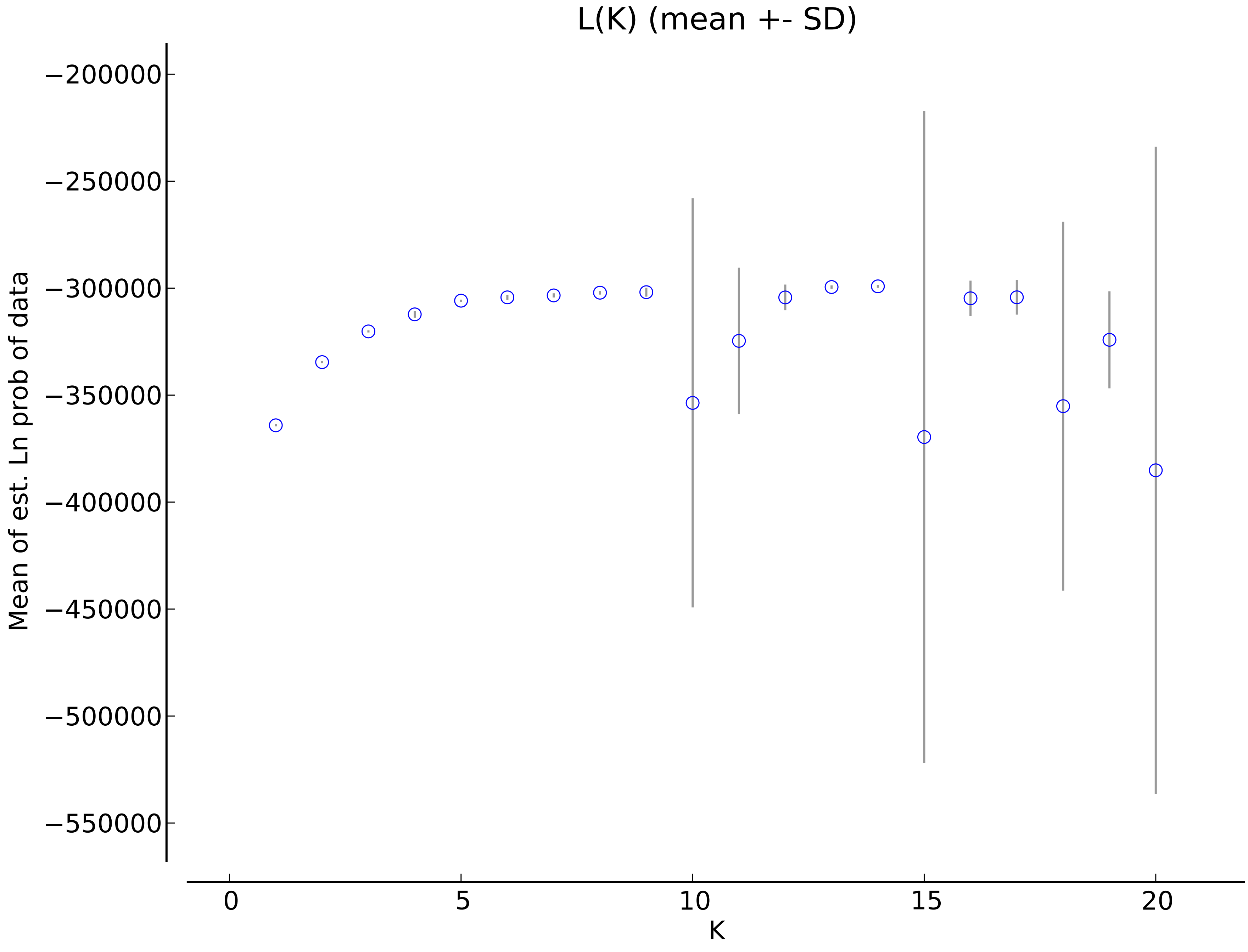
L(K) (mean +- SD)
Mean of est. Ln prob of data
−200000
−250000
−300000
−350000
−400000
−450000
−500000
−550000
0
5
10
15
20
K

**Supplementary Figure 4. STRUCTURE output for k2-10**

Full STRUCTURE output for dataset D1, depicting population assignments and admixture for K2-10. The likelihood of each evaluated number of K clusters from 1:20 plateaued at K = 5, with the standard deviation across runs increasing rapidly after this point (Supplementary Figure 3). See results text for descriptions of these analyses. At K2, the first partition divides the dataset into broad northern and southern Amazonian groups, with all Andean and Central American samples assigned to predominantly southern Amazonian genetic provenance, with some northern admixture. Western Napo individuals are detected as an approximately even mixture of northern and southern Amazonian genomes. At K5, the five identified clusters broadly correspond to wide biogeographic Amazonian regions which encompass multiple areas of endemism (see text). For each barplot, colors are sampled randomly from a 20 color viridis color palette for each run (i.e., they are not synonymous across values of K, see supplementary R script). The tree to the left corresponds to the RAxML result using the 50% haplotype dataset, with tip labels and colors indicating group membership to one of eighteen population-areas. Colored tip labels correspond to clade label colors in Figures 3 and 5. At higher values of K, the broad-scale population assignments inferred at K5 are similar, however additional admixture components are inferred for most groups. The introgressed western Napo lineage is eventually placed into its own cluster at K9-10.

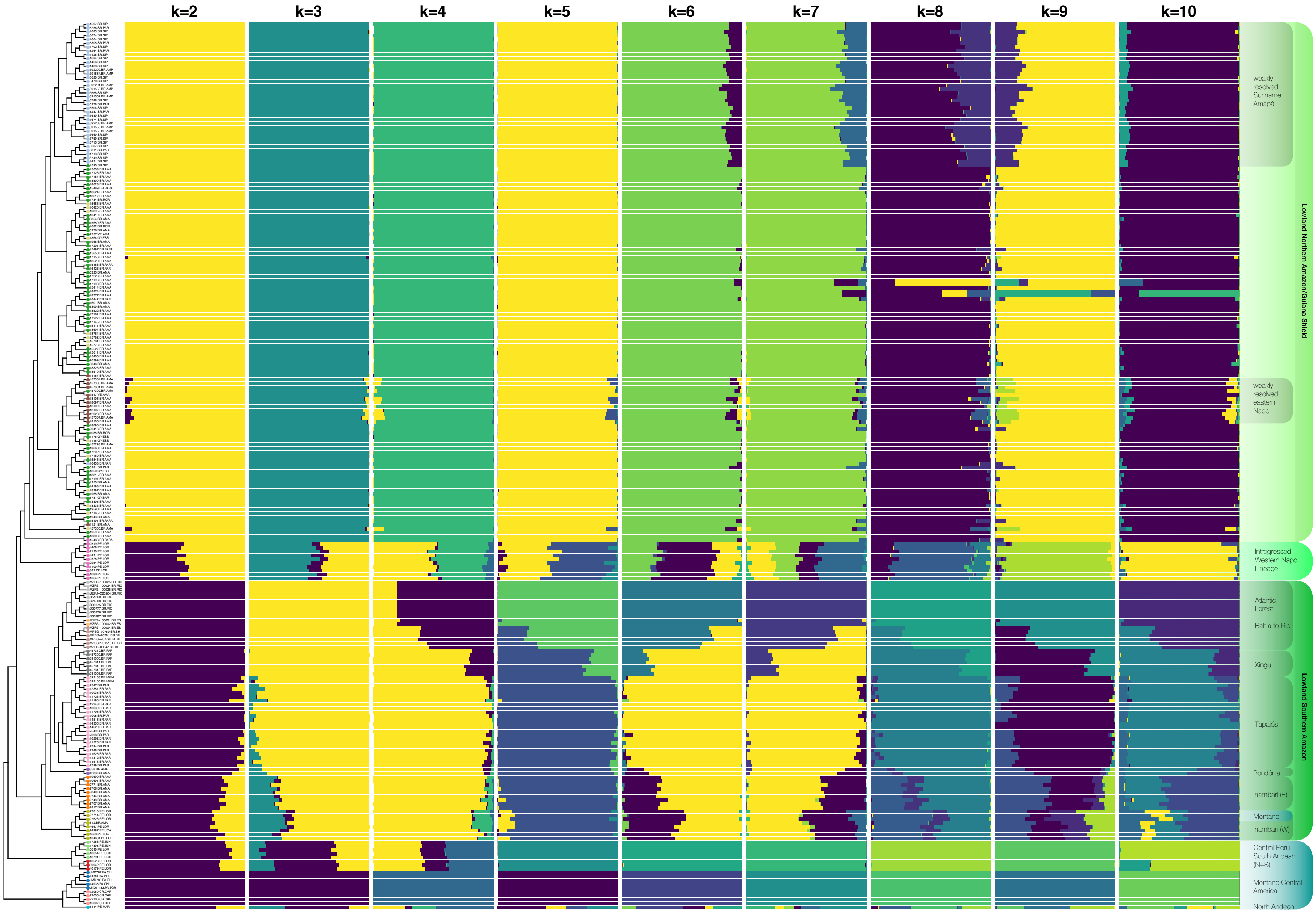
k=2
k=3
k=4
k=5
k=6
k=7
k=8
k=9
k=10
weakly resolved Suriname, Amapá
weakly resolved eastern Napo
Lowland Northern Amazon/Guiana Shield
Introgressed Western Napo Lineage
Atlantic Forest
Bahia to Rio
Xingu
Tapajós
Rondônia
Inambari (E)
Montane
Inambari (W)
Lowland Southern Amazon
Central Peru South Andean (N+S)
Montane Central America
North Andean

**Supplementary Figure 5. fineRADstructure population assignment dendrogram**
Clustering dendrogram generated from fineRADstructure population assignment. Note: this is not a phylogenetic hypothesis, but rather, a clustering based on genomic similarity which considers data from the full coancestry matrix. Tip labels correspond to population codes used internally for R scripts and other analyses. Each of these codes has a 1:1 correspondence with the labeled localities in Figures 3 and 4. CAMA: North Andean – San Martín, Peru, CACR: Central America - Costa Rica, CAPA: Central America – Panama, CPS:, South Andean Peru (South), CPN: South Andean Peru (North), INAMBARI: Western Inambari endemic, INAMBARIE: Eastern Inambari endemic , RONDONIA: Rondônia endemic, TAPAJOS: Tapajós endemic, XINGU: Xingu endemic, AFBAHIA: Atlantic Forest – Bahia, AFES: Atlantic Forest – Espírito Santo, AFRIO: Atlantic Forest – Rio, PH (putative hybrid): Western Napo introgressed lineage, GSIMERI: unresolved Jaú, GSNAPO: weakly resolved eastern Napo, GS: weakly resolved Guiana Shield, GSSR: weakly resolved Suriname + Amapá.

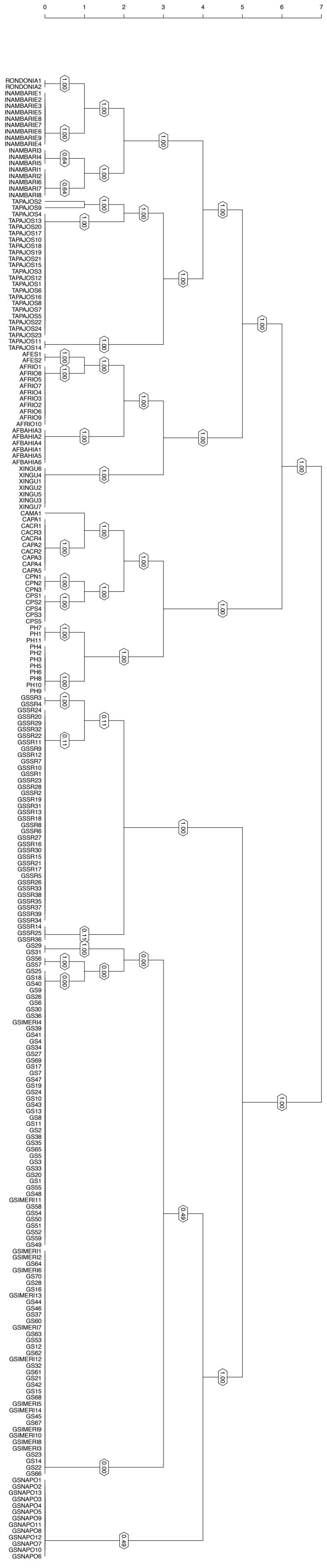
0
1
2
3
4
5
6
7
RONDONIA1
RONDONIA2
INAMBARIE1
INAMBARIE2
INAMBARIE3
INAMBARIE5
INAMBARIE8
INAMBARIE7
INAMBARIE6
INAMBARIE9
INAMBARIE4
INAMBARI3
INAMBARI4
INAMBARI5
INAMBARI1
INAMBARI2
INAMBARI6
INAMBARI7
INAMBARI8
TAPAJOS2
TAPAJOS9
TAPAJOS4
TAPAJOS13
TAPAJOS20
TAPAJOS17
TAPAJOS10
TAPAJOS18
TAPAJOS19
TAPAJOS21
TAPAJOS15
TAPAJOS3
TAPAJOS12
TAPAJOS1
TAPAJOS6
TAPAJOS16
TAPAJOS8
TAPAJOS7
TAPAJOS5
TAPAJOS22
TAPAJOS24
TAPAJOS23
TAPAJOS11
TAPAJOS14
AFES1
AFES2
AFRIO1
AFRIO8
AFRIO5
AFRIO7
AFRIO4
AFRIO3
AFRIO2
AFRIO6
AFRIO9
AFRIO10
AFBAHIA3
AFBAHIA2
AFBAHIA4
AFBAHIA1
AFBAHIA5
AFBAHIA6
XINGU6
XINGU4
XINGU1
XINGU2
XINGU5
XINGU3
XINGU7
CAMA1
CAPA1
CACR1
CACR3
CACR4
CAPA2
CACR2
CAPA3
CAPA4
CAPA5
CPN1
CPN2
CPN3
CPS1
CPS2
CPS4
CPS3
CPS5
PH7
PH1
PH11
PH4
PH2
PH3
PH5
PH6
PH8
PH10
PH9
GSSR3
GSSR4
GSSR24
GSSR20
GSSR29
GSSR32
GSSR22
GSSR11
GSSR9
GSSR12
GSSR7
GSSR10
GSSR1
GSSR23
GSSR28
GSSR2
GSSR19
GSSR31
GSSR13
GSSR18
GSSR8
GSSR6
GSSR27
GSSR16
GSSR30
GSSR15
GSSR21
GSSR17
GSSR5
GSSR26
GSSR33
GSSR38
GSSR35
GSSR37
GSSR39
GSSR34
GSSR14
GSSR25
GSSR36
GS29
GS31
GS56
GS57
GS25
GS18
GS40
GS9
GS26
GS6
GS30
GS36
GSIMERI4
GS39
GS41
GS4
GS34
GS27
GS69
GS17
GS7
GS47
GS19
GS24
GS10
GS43
GS13
GS8
GS11
GS2
GS38
GS35
GS65
GS5
GS3
GS33
GS20
GS1
GS55
GS48
GSIMERI11
GS58
GS54
GS50
GS51
GS52
GS59
GS49
GSIMERI1
GSIMERI2
GS64
GSIMERI6
GS70
GS28
GS16
GSIMERI13
GS44
GS46
GS37
GS60
GSIMERI7
GS63
GS53
GS12
GS62
GSIMERI12
GS32
GS61
GS21
GS42
GS15
GS68
GSIMERI5
GSIMERI14
GS45
GS67
GSIMERI9
GSIMERI10
GSIMERI8
GSIMERI3
GS23
GS14
GS22
GS66
GSNAPO1
GSNAPO2
GSNAPO13
GSNAPO3
GSNAPO4
GSNAPO5
GSNAPO9
GSNAPO11
GSNAPO8
GSNAPO12
GSNAPO7
GSNAPO10
GSNAPO6

**Supplementary Figure 6. RadPainter coancestry matrix**
The raw coancestry matrix from the full haplotyle dataset, output from the fineRADstructure program. See Supplementary Figure 5 caption for descriptions of locality codes, matching those in Figure 3.

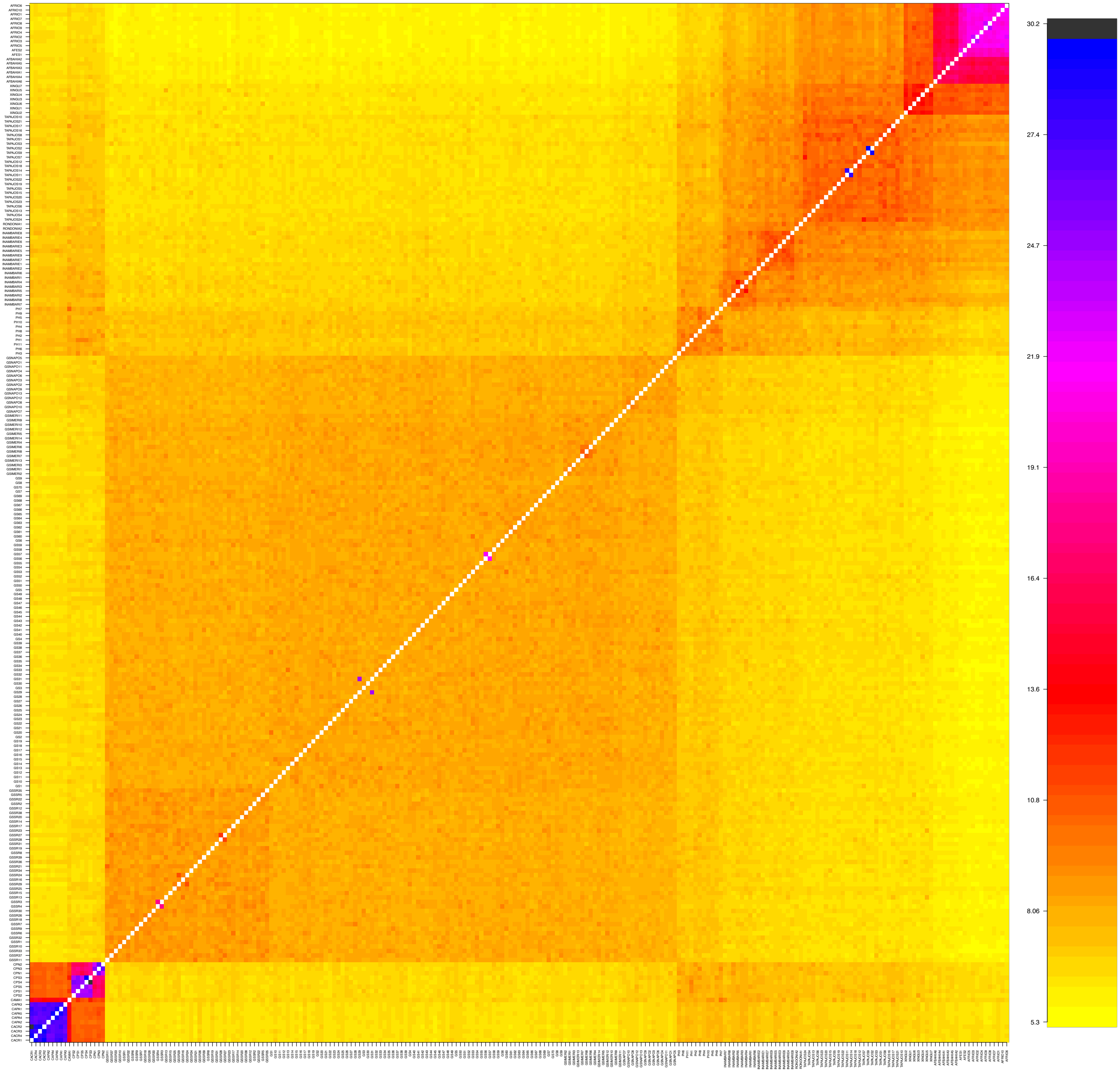
30.2
27.4
24.7
21.9
19.1
16.4
13.6
10.8
8.06
5.3

**Supplementary Figure 7. RadPainter coancestry matrix**
The coancestry matrix from the full haplotype dataset, with values averaged within each of 18 focal population-areas. See Supplementary Figure 5 caption for descriptions of localities, matching those in Figure 3.

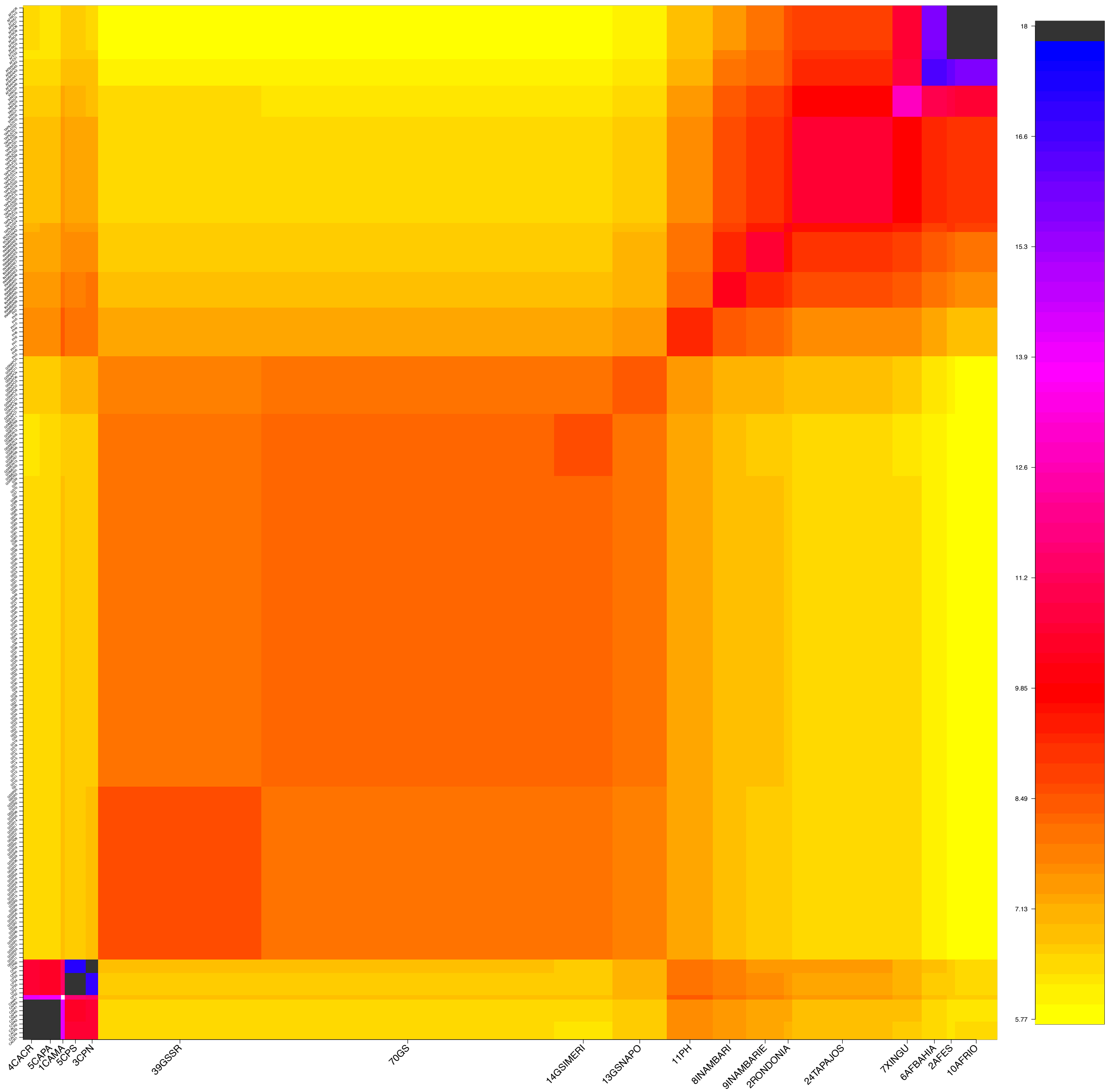
4CACR
5CAPA
1CAMA
5CPS
3CPN
39GSSR
70GS
14GSIMERI
13GSNAPO
11PH
8INAMBARI
9INAMBARIE
2RONDONIA
24TAPAJOS
7XINGU
6AFBAHIA
2AFES
10AFRIO
18
16.6
15.3
13.9
12.6
11.2
9.85
8.49
7.13
5.77

**Supplementary Figure 8a-h. K-means clustering of the coancestry matrix**
The Lawson et al. (2012) 'normalized PCA' approach provided with the fineRADstructure software captured 89% of the variance in the genetic data on the first four component axes (axis 1: 51.4%, axis 2, 24.9%, axis 3: 7.77%, axis 4: 4.94%). Thus, the coancestry matrix reflects substantially more information than standard PCoA/PCA of SNP data (Supplementary Figure 2a-h). K-means phenetic clustering of the coancestry matrix more finely partitions the genetic data and explains a much greater proportion of the overall genetic variance than K-means clustering of the raw SNP data. Top panel: normalized PCA projection of all individuals on the first two component axes, which capture ~76% of the point variability. Plotting symbols and colored convex hulls reflect cluster assignment. Figure 8a indicates membership to each of eighteen focal population-areas (i.e., not K-means assignments, see Supplemental Appendix text for justification) and are shown as minimum convex hulls in coancestry PC space (top) as well as projected onto a map (bottom, also shown in Figure 4). Figure 8b shows the K-means clustering solution of the coancestry matrix when K is fixed to 18 (i.e., not based on BIC scores). This clustering produces a similar set of groups very similar to Figure 8a. In Figure 8c, we show the K-means clustering solution of the coancestry matrix, with a BIC minimum plateau of ~8. This set of groups is concordant with hierarchical strata determined in earlier analyses but is also further partitioned relative to standard PC analyses on our SNP data. This clustering solution identified 1) Central America (Clade A2 in Figure 3), 2) South Andean Peru + San Martín (North Andean Peru); (Clade B + Clade A1 in Figure 3), 3) western Napo (Clade E in Figure 3), 4) Inambari + Rondônia (Clades C1, C3, and C4 in Figure 3), 5) Tapajós + Xingu (Clades C5 + C6 in Figure 3), 6) eastern Napo, Jaú, western Guiana shield, 7) Suriname + Amapá (Clade D in Figure 3) and 8) the Atlantic Forest (Clade C7), as separate groups which best explain the variance in the data. Notably, this solution is compatible with our phylogenetic hypothesis, except for the clustering of our single San Martín sample with geographically proximate Peruvian populations, rather than Central American populations (see discussion). This solution generally also recapitulates subspecies boundaries (Figure 2).

### Supplementary Figure 8a

**coancestry matrix, 18 phylo−regions PCoA k−means K = NA**

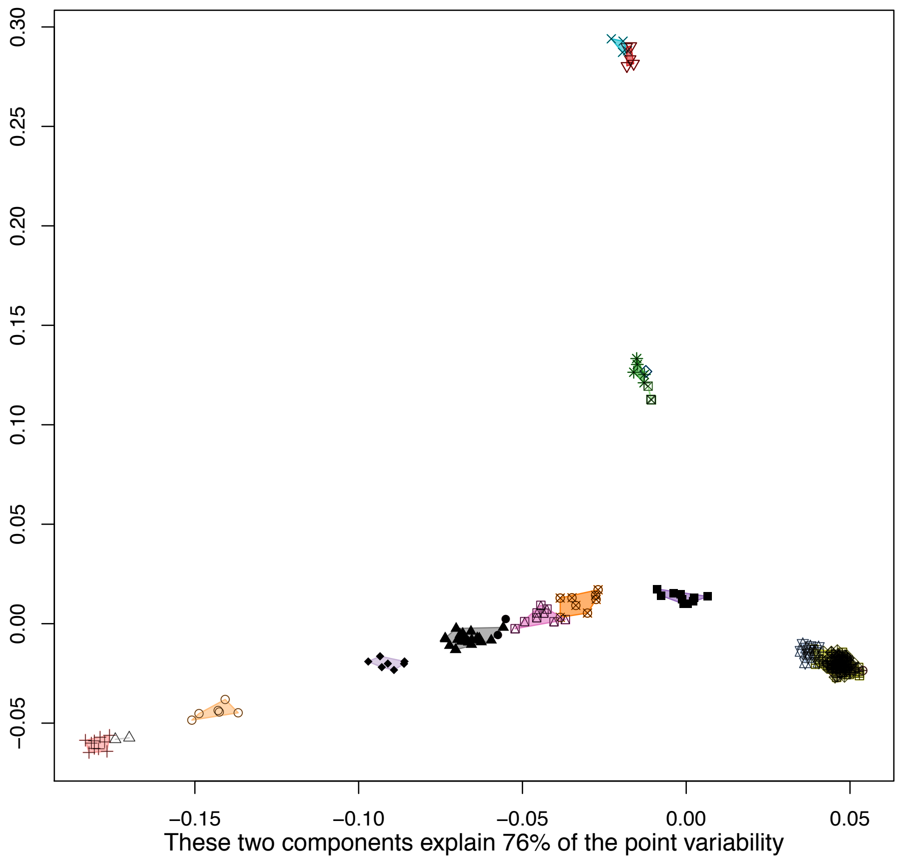

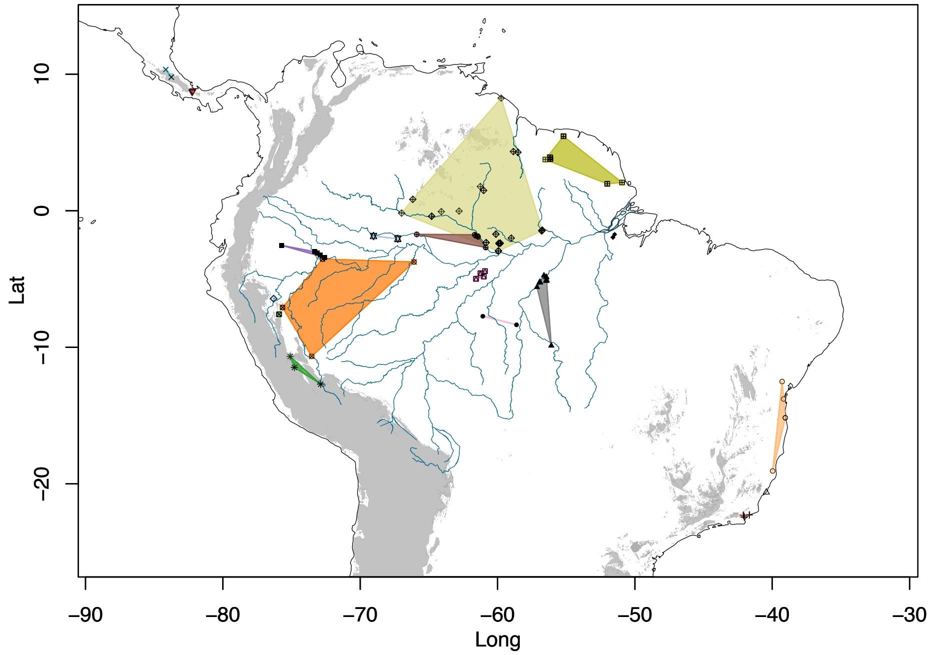

### Supplementary Figure 8b

**coancestry matrix, k=18 kmeans solution PCoA k−means K = 18**

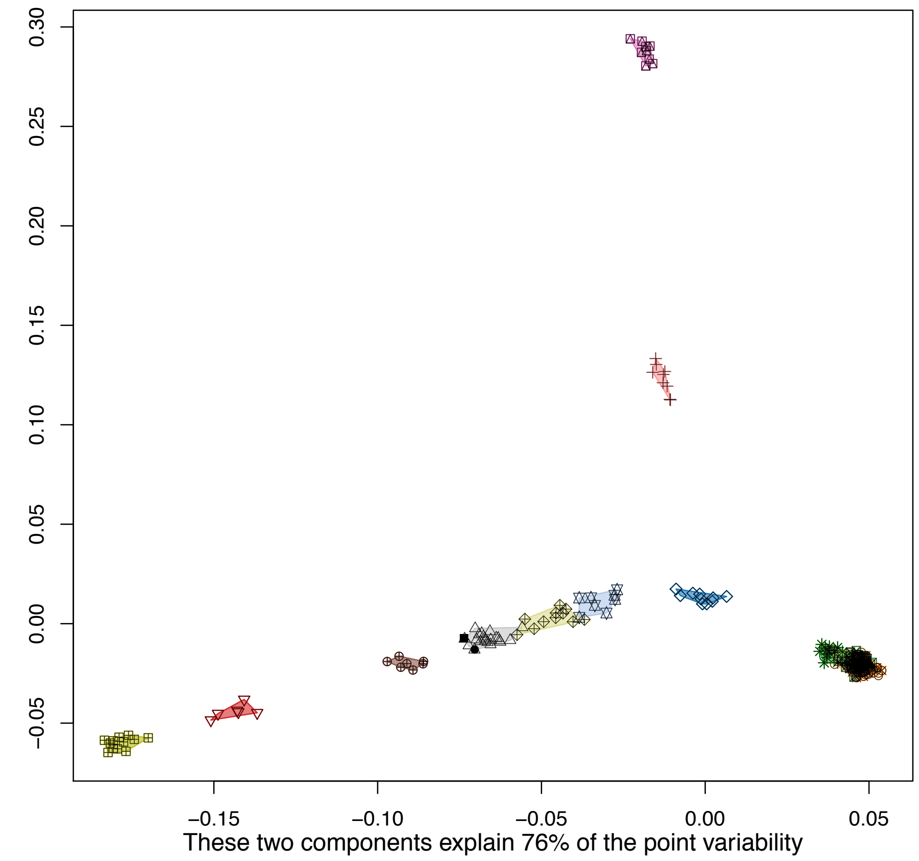

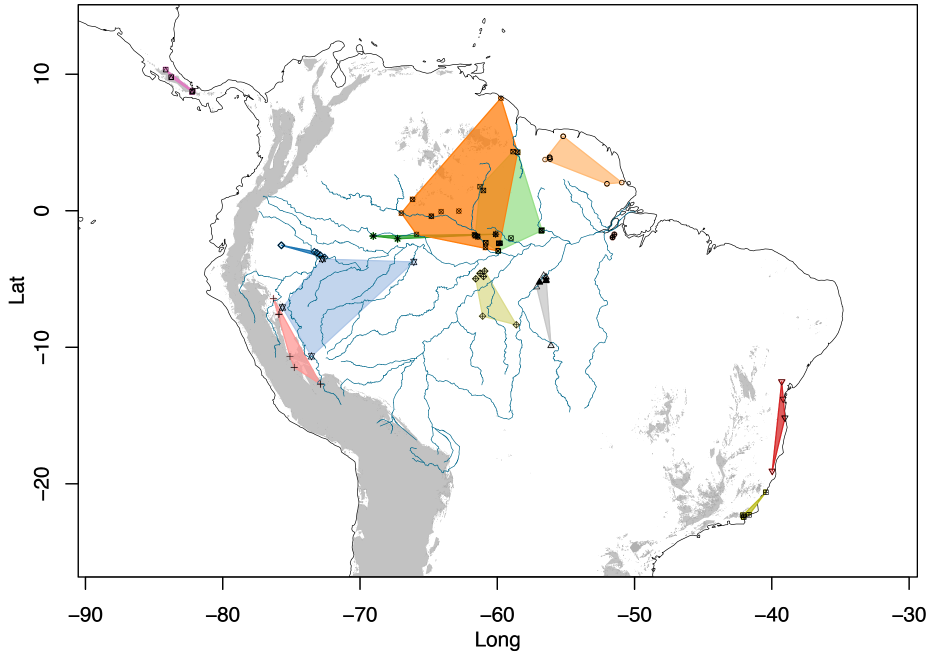

#### Supplementary Figure 8c

**coancestry matrix, BIC min K=8 kmeans solution PCoA k−means K = 8**

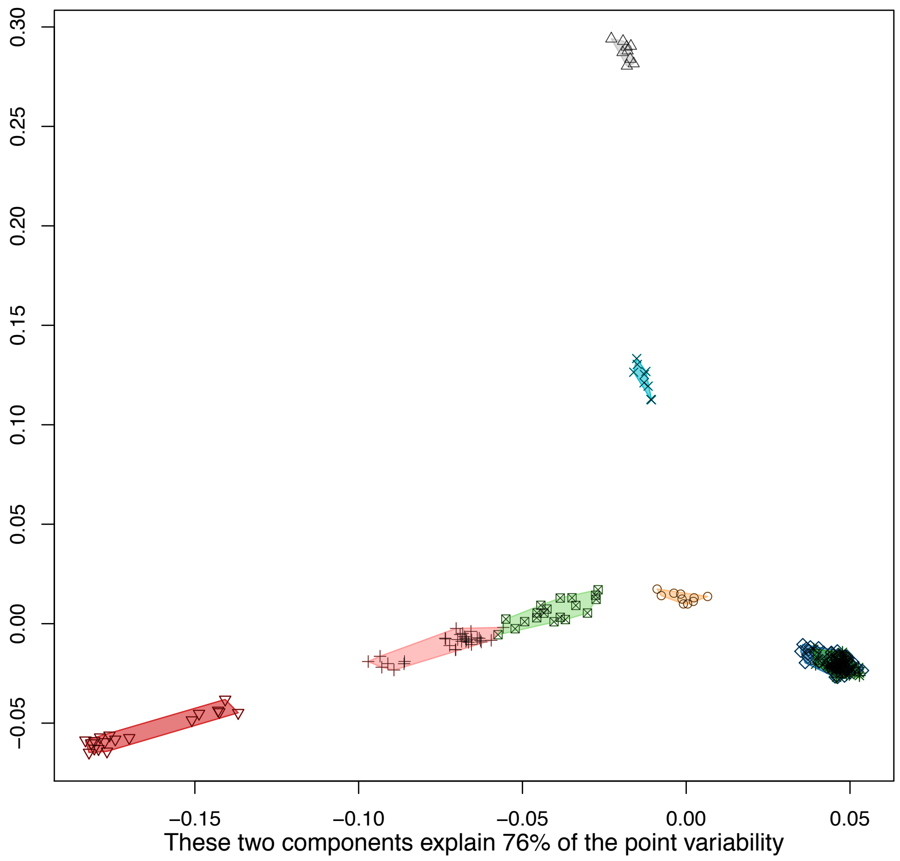

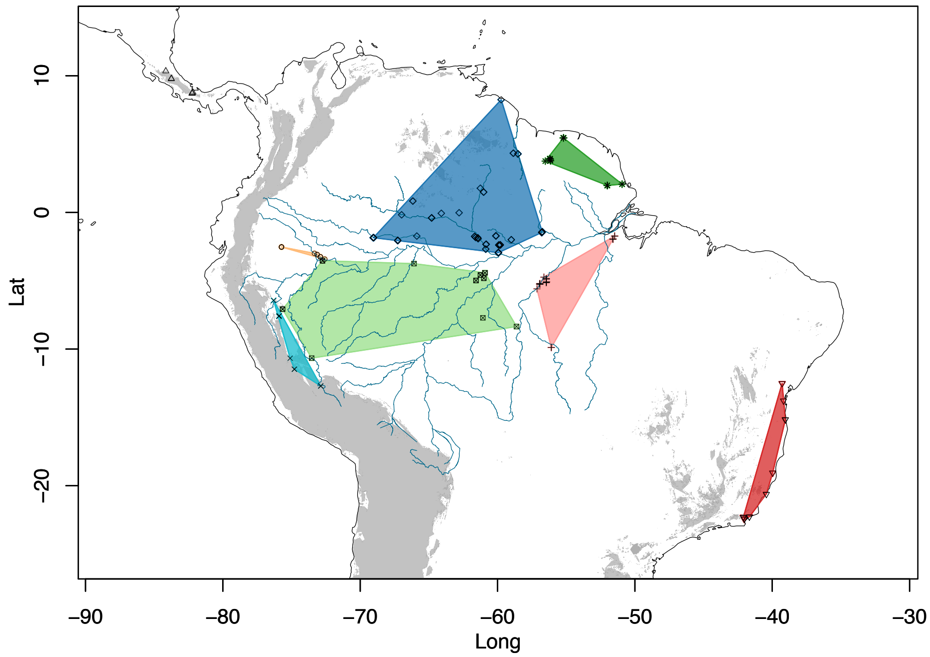

**Supplementary Figure 9. Estimates of Inbreeding coefficients $F_{IS}$**

Most populations were detected to be significantly inbred on the basis of $F_{IS}$ ($F_{IS} > 1$, with lower 95% confidence intervals > 0. Panamanian, Costa Rican, South Andean (North clade), Rondônia, and Espírito Santo clades had 95% confidence intervals which overlapped zero, and thus cannot be confidently inferred to have positive or negative $F_{IS}$. The simulated F1 (SIMF1) population, however, did have significantly negative $F_{IS}$, as predicted. This pattern suggests that the introgressed western Napo population, which was detected to have a significantly positive $F_{IS}$, is not likely to include F1 individuals. The confidence intervals for eastern Napo, Jaú, Inambari and western Napo popularions are generally overlapping, with similar means (mean of mean estimates ~0.17, SD of mean estimates ~0.02). Locality codes are the same as those described in the caption for Supplementary Figure 5.

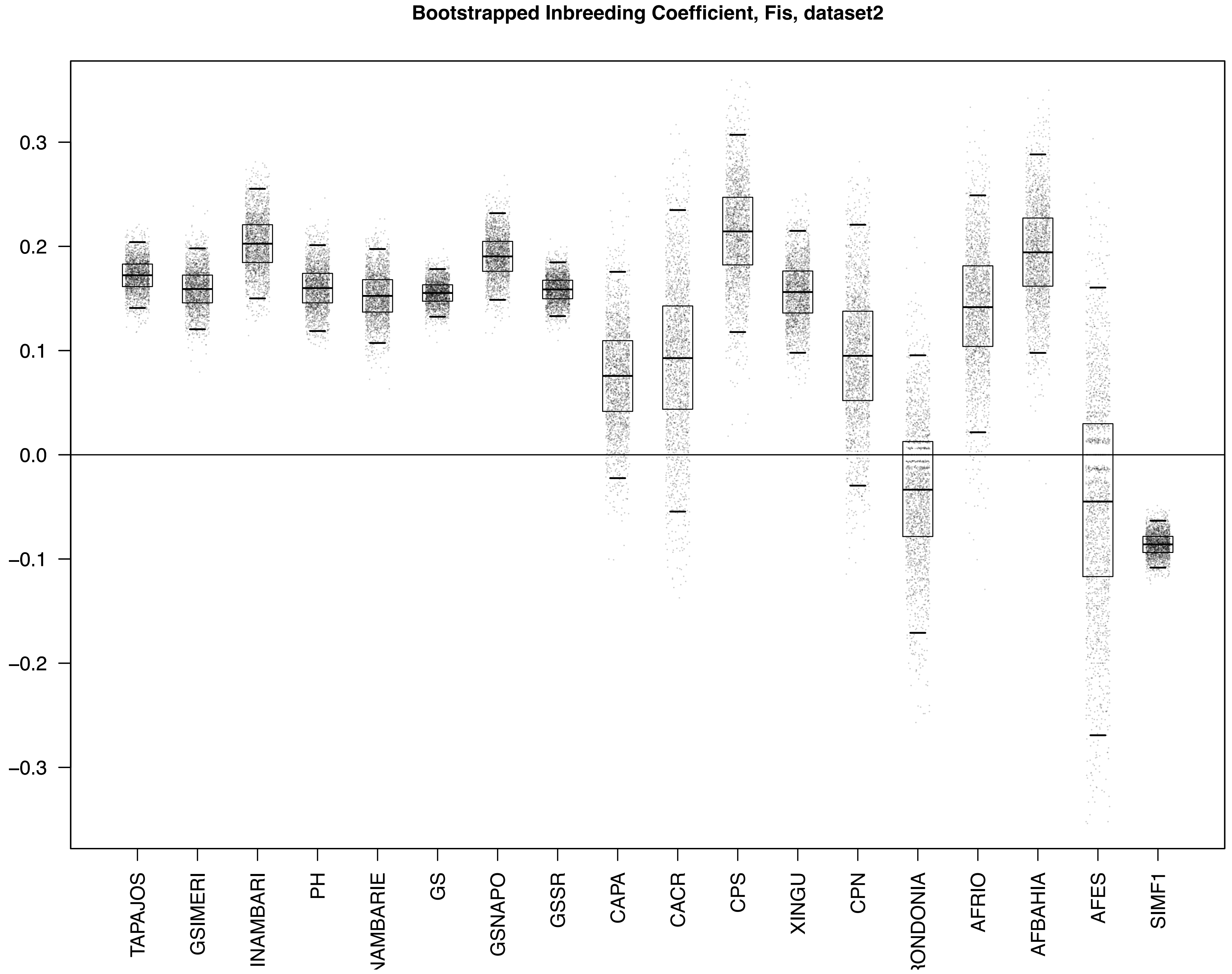

Bootstrapped Inbreeding Coefficient, Fis, dataset2
0.3
0.2
0.1
0.0
−0.1
−0.2
−0.3
TAPAJOS
GSIMERI
INAMBARI
PH
GS
GSNAPO
GSSR
CAPA
CACR
CPS
XINGU
CPN
AFRIO
AFBAHIA
AFES
SIMF1

**Supplementary Figure 10. Population pairwise $F_{st}$**

We estimated pairwise Weir and Cockerham's (Weir and Cockerham, 1984) $F_{st}$ among all 18 focal areas, and evaluated significance using 1000 bootstrapped datasets to estimate 95% confidence intervals using the 'assigner' R package (Gosselin et al., 2016). Here, we show these results plotted as heatmaps for each SNP dataset. See main text for additional discussion. Locality codes are the same as those in the caption for Supplementary Figure 5.

### Supplementary Figure 10a
### SNP dataset 1

| | AFBAHIA | AFES | AFRIO | CACR | CAMA | CAPA | CPN | CPS | GS | GSIMERI | GSNAPO | GSSR | INAMBARI | INAMBARIE | PH | RONDONIA | TAPAJOS | XINGU |
|---|---|---|---|---|---|---|---|---|---|---|---|---|---|---|---|---|---|---|
| AFBAHIA | | 0.09<br>0.15 | 0.18<br>0.23 | 0.61<br>0.66 | 0.45<br>0.51 | 0.64<br>0.68 | 0.52<br>0.58 | 0.54<br>0.59 | 0.26<br>0.29 | 0.28<br>0.32 | 0.27<br>0.31 | 0.27<br>0.30 | 0.24<br>0.28 | 0.21<br>0.26 | 0.25<br>0.29 | 0.22<br>0.28 | 0.18<br>0.21 | 0.16<br>0.20 |
| AFES | 0.12 | | 0.13<br>0.19 | 0.77<br>0.82 | 0.61<br>0.67 | 0.79<br>0.83 | 0.66<br>0.71 | 0.64<br>0.69 | 0.28<br>0.33 | 0.29<br>0.34 | 0.28<br>0.33 | 0.29<br>0.33 | 0.27<br>0.33 | 0.25<br>0.30 | 0.28<br>0.33 | 0.30<br>0.37 | 0.21<br>0.26 | 0.21<br>0.26 |
| AFRIO | 0.21 | 0.16 | | 0.70<br>0.74 | 0.62<br>0.67 | 0.71<br>0.75 | 0.63<br>0.68 | 0.63<br>0.68 | 0.33<br>0.36 | 0.38<br>0.41 | 0.37<br>0.41 | 0.34<br>0.38 | 0.36<br>0.41 | 0.34<br>0.38 | 0.36<br>0.40 | 0.39<br>0.46 | 0.27<br>0.31 | 0.29<br>0.34 |
| CACR | 0.64 | 0.80 | 0.72 | | 0.68<br>0.75 | 0.04<br>0.13 | 0.71<br>0.76 | 0.66<br>0.71 | 0.33<br>0.38 | 0.37<br>0.41 | 0.36<br>0.41 | 0.35<br>0.39 | 0.42<br>0.47 | 0.42<br>0.47 | 0.36<br>0.41 | 0.59<br>0.65 | 0.39<br>0.44 | 0.50<br>0.55 |
| CAMA | 0.48 | 0.64 | 0.65 | 0.72 | | 0.71<br>0.78 | 0.56<br>0.64 | 0.54<br>0.62 | 0.18<br>0.24 | 0.19<br>0.25 | 0.18<br>0.23 | 0.19<br>0.25 | 0.20<br>0.27 | 0.21<br>0.28 | 0.16<br>0.23 | 0.22<br>0.32 | 0.24<br>0.30 | 0.31<br>0.38 |
| CAPA | 0.66 | 0.81 | 0.74 | 0.09 | 0.74 | | 0.73<br>0.78 | 0.68<br>0.73 | 0.34<br>0.38 | 0.39<br>0.43 | 0.39<br>0.43 | 0.36<br>0.40 | 0.46<br>0.51 | 0.45<br>0.50 | 0.39<br>0.44 | 0.64<br>0.69 | 0.41<br>0.46 | 0.54<br>0.59 |
| CPN | 0.55 | 0.68 | 0.65 | 0.73 | 0.60 | 0.76 | | 0.34<br>0.42 | 0.27<br>0.32 | 0.29<br>0.34 | 0.29<br>0.33 | 0.28<br>0.32 | 0.32<br>0.37 | 0.32<br>0.38 | 0.28<br>0.33 | 0.46<br>0.52 | 0.31<br>0.36 | 0.41<br>0.46 |
| CPS | 0.56 | 0.67 | 0.65 | 0.68 | 0.58 | 0.71 | 0.38 | | 0.29<br>0.33 | 0.32<br>0.37 | 0.32<br>0.36 | 0.30<br>0.34 | 0.36<br>0.41 | 0.35<br>0.40 | 0.31<br>0.36 | 0.49<br>0.54 | 0.33<br>0.37 | 0.43<br>0.49 |
| GS | 0.28 | 0.31 | 0.35 | 0.35 | 0.21 | 0.36 | 0.29 | 0.31 | | 0.00<br>0.01 | 0.01<br>0.02 | 0.02<br>0.02 | 0.12<br>0.15 | 0.13<br>0.16 | 0.08<br>0.10 | 0.11<br>0.15 | 0.15<br>0.18 | 0.19<br>0.22 |
| GSIMERI | 0.30 | 0.32 | 0.40 | 0.39 | 0.22 | 0.41 | 0.32 | 0.34 | 0.01* | | 0.01<br>0.02 | 0.02<br>0.03 | 0.13<br>0.16 | 0.14<br>0.17 | 0.08<br>0.11 | 0.12<br>0.16 | 0.16<br>0.19 | 0.20<br>0.23 |
| GSNAPO | 0.29 | 0.31 | 0.39 | 0.38 | 0.20 | 0.41 | 0.31 | 0.34 | 0.01 | 0.02 | | 0.03<br>0.04 | 0.12<br>0.14 | 0.12<br>0.15 | 0.07<br>0.09 | 0.10<br>0.14 | 0.15<br>0.18 | 0.19<br>0.22 |
| GSSR | 0.29 | 0.31 | 0.36 | 0.37 | 0.22 | 0.38 | 0.30 | 0.32 | 0.02 | 0.03 | 0.03 | | 0.13<br>0.15 | 0.14<br>0.17 | 0.08<br>0.11 | 0.12<br>0.16 | 0.17<br>0.19 | 0.20<br>0.23 |
| INAMBARI | 0.26 | 0.30 | 0.39 | 0.45 | 0.23 | 0.49 | 0.35 | 0.38 | 0.13 | 0.14 | 0.13 | 0.14 | | 0.05<br>0.07 | 0.06<br>0.08 | 0.03<br>0.08 | 0.08<br>0.10 | 0.13<br>0.16 |
| INAMBARIE | 0.24 | 0.28 | 0.36 | 0.44 | 0.25 | 0.47 | 0.35 | 0.38 | 0.14 | 0.15 | 0.14 | 0.15 | 0.06 | | 0.08<br>0.10 | 0.01<br>0.05 | 0.06<br>0.08 | 0.12<br>0.15 |
| PH | 0.27 | 0.31 | 0.38 | 0.39 | 0.20 | 0.42 | 0.31 | 0.33 | 0.09 | 0.09 | 0.08 | 0.09 | 0.07 | 0.09 | | 0.06<br>0.10 | 0.10<br>0.13 | 0.15<br>0.18 |
| RONDONIA | 0.25 | 0.34 | 0.42 | 0.62 | 0.27 | 0.67 | 0.49 | 0.52 | 0.13 | 0.14 | 0.12 | 0.14 | 0.05 | 0.03 | 0.08 | | 0.01<br>0.05 | 0.08<br>0.13 |
| TAPAJOS | 0.19 | 0.23 | 0.29 | 0.42 | 0.27 | 0.44 | 0.33 | 0.35 | 0.17 | 0.18 | 0.16 | 0.18 | 0.09 | 0.07 | 0.12 | 0.03 | | 0.06<br>0.09 |
| XINGU | 0.18 | 0.24 | 0.31 | 0.53 | 0.34 | 0.57 | 0.43 | 0.46 | 0.20 | 0.22 | 0.20 | 0.21 | 0.14 | 0.13 | 0.16 | 0.10 | 0.08 | |

FST: 0.8, 0.6, 0.4, 0.2

### Supplementary Figure 10b
### SNP dataset 1m

| | AFBAHIA | AFES | AFRIO | CACR | CAMA | CAPA | CPN | CPS | GS | GSIMERI | GSNAPO | GSSR | INAMBARI | INAMBARIE | PH | RONDONIA | TAPAJOS | XINGU |
|---|---|---|---|---|---|---|---|---|---|---|---|---|---|---|---|---|---|---|
| AFBAHIA | | 0.01<br>0.14 | 0.10<br>0.25 | 0.51<br>0.66 | 0.38<br>0.56 | 0.53<br>0.68 | 0.46<br>0.61 | 0.47<br>0.61 | 0.24<br>0.38 | 0.26<br>0.40 | 0.26<br>0.40 | 0.25<br>0.40 | 0.25<br>0.40 | 0.19<br>0.33 | 0.24<br>0.39 | 0.16<br>0.33 | 0.17<br>0.30 | 0.11<br>0.25 |
| AFES | 0.08 | | 0.03<br>0.21 | 0.66<br>0.81 | 0.49<br>0.70 | 0.69<br>0.83 | 0.60<br>0.75 | 0.59<br>0.74 | 0.29<br>0.43 | 0.30<br>0.45 | 0.31<br>0.45 | 0.31<br>0.46 | 0.32<br>0.47 | 0.25<br>0.41 | 0.28<br>0.45 | 0.22<br>0.43 | 0.21<br>0.37 | 0.19<br>0.37 |
| AFRIO | 0.18 | 0.13 | | 0.59<br>0.75 | 0.57<br>0.72 | 0.61<br>0.76 | 0.59<br>0.72 | 0.58<br>0.71 | 0.32<br>0.46 | 0.36<br>0.51 | 0.37<br>0.51 | 0.34<br>0.48 | 0.38<br>0.53 | 0.32<br>0.47 | 0.35<br>0.51 | 0.31<br>0.51 | 0.27<br>0.41 | 0.26<br>0.43 |
| CACR | 0.59 | 0.74 | 0.68 | | 0.55<br>0.78 | 0.00<br>0.14 | 0.63<br>0.80 | 0.56<br>0.74 | 0.27<br>0.42 | 0.31<br>0.46 | 0.32<br>0.47 | 0.29<br>0.44 | 0.36<br>0.54 | 0.33<br>0.49 | 0.31<br>0.48 | 0.45<br>0.64 | 0.31<br>0.47 | 0.44<br>0.59 |
| CAMA | 0.48 | 0.61 | 0.65 | 0.67 | | 0.54<br>0.78 | 0.56<br>0.78 | 0.50<br>0.72 | 0.08<br>0.26 | 0.07<br>0.27 | 0.09<br>0.29 | 0.06<br>0.26 | 0.11<br>0.34 | 0.12<br>0.34 | 0.06<br>0.28 | 0.08<br>0.39 | 0.18<br>0.38 | 0.25<br>0.45 |
| CAPA | 0.61 | 0.77 | 0.69 | 0.07* | 0.67 | | 0.63<br>0.80 | 0.58<br>0.74 | 0.27<br>0.42 | 0.32<br>0.47 | 0.33<br>0.48 | 0.29<br>0.45 | 0.37<br>0.54 | 0.33<br>0.51 | 0.32<br>0.49 | 0.50<br>0.68 | 0.32<br>0.48 | 0.46<br>0.61 |
| CPN | 0.54 | 0.68 | 0.65 | 0.72 | 0.68 | 0.73 | | 0.21<br>0.45 | 0.23<br>0.36 | 0.27<br>0.41 | 0.27<br>0.41 | 0.25<br>0.38 | 0.26<br>0.40 | 0.25<br>0.40 | 0.22<br>0.38 | 0.38<br>0.55 | 0.24<br>0.39 | 0.33<br>0.50 |
| CPS | 0.55 | 0.67 | 0.65 | 0.66 | 0.62 | 0.66 | 0.33 | | 0.25<br>0.38 | 0.28<br>0.42 | 0.29<br>0.43 | 0.26<br>0.40 | 0.30<br>0.45 | 0.29<br>0.43 | 0.23<br>0.38 | 0.38<br>0.58 | 0.26<br>0.39 | 0.36<br>0.51 |
| GS | 0.31 | 0.37 | 0.39 | 0.35 | 0.17 | 0.35 | 0.30 | 0.32 | | 0.00<br>0.02 | 0.00<br>0.02 | 0.01<br>0.02 | 0.09<br>0.19 | 0.09<br>0.16 | 0.06<br>0.14 | 0.06<br>0.18 | 0.14<br>0.23 | 0.16<br>0.27 |
| GSIMERI | 0.33 | 0.38 | 0.44 | 0.39 | 0.17 | 0.40 | 0.34 | 0.35 | 0.01* | | 0.00<br>0.03 | 0.01<br>0.03 | 0.10<br>0.22 | 0.09<br>0.18 | 0.07<br>0.15 | 0.07<br>0.19 | 0.14<br>0.23 | 0.18<br>0.29 |
| GSNAPO | 0.34 | 0.39 | 0.44 | 0.40 | 0.19 | 0.41 | 0.34 | 0.36 | 0.01* | 0.01* | | 0.02<br>0.05 | 0.10<br>0.22 | 0.09<br>0.18 | 0.05<br>0.14 | 0.06<br>0.18 | 0.14<br>0.23 | 0.17<br>0.28 |
| GSSR | 0.33 | 0.38 | 0.42 | 0.37 | 0.16 | 0.37 | 0.32 | 0.33 | 0.01 | 0.02 | 0.03 | | 0.10<br>0.21 | 0.11<br>0.19 | 0.08<br>0.16 | 0.08<br>0.21 | 0.16<br>0.25 | 0.18<br>0.30 |
| INAMBARI | 0.32 | 0.40 | 0.46 | 0.45 | 0.23 | 0.46 | 0.33 | 0.38 | 0.14 | 0.16 | 0.16 | 0.16 | | 0.03<br>0.13 | 0.04<br>0.12 | 0.04<br>0.20 | 0.07<br>0.18 | 0.12<br>0.26 |
| INAMBARIE | 0.26 | 0.33 | 0.40 | 0.41 | 0.23 | 0.42 | 0.33 | 0.37 | 0.12 | 0.13 | 0.13 | 0.14 | 0.07 | | 0.06<br>0.14 | 0.00<br>0.10 | 0.05<br>0.15 | 0.10<br>0.21 |
| PH | 0.32 | 0.37 | 0.44 | 0.40 | 0.18 | 0.41 | 0.30 | 0.31 | 0.10 | 0.11 | 0.09 | 0.12 | 0.08 | 0.10 | | 0.00<br>0.12 | 0.09<br>0.17 | 0.13<br>0.25 |
| RONDONIA | 0.25 | 0.33 | 0.42 | 0.56 | 0.24 | 0.60 | 0.47 | 0.49 | 0.12 | 0.13 | 0.12 | 0.14 | 0.12 | 0.04* | 0.05* | | 0.01<br>0.12 | 0.04<br>0.16 |
| TAPAJOS | 0.23 | 0.29 | 0.34 | 0.39 | 0.28 | 0.40 | 0.32 | 0.32 | 0.18 | 0.18 | 0.18 | 0.20 | 0.12 | 0.10 | 0.13 | 0.06 | | 0.06<br>0.12 |
| XINGU | 0.18 | 0.28 | 0.34 | 0.52 | 0.36 | 0.54 | 0.42 | 0.44 | 0.22 | 0.23 | 0.23 | 0.23 | 0.19 | 0.16 | 0.19 | 0.10 | 0.09 | |

FST
0.6
0.4
0.2

### Supplementary Figure 10c
### SNP dataset 2

| | AFBAHIA | AFES | AFRIO | CACR | CAMA | CAPA | CPN | CPS | GS | GSIMERI | GSNAPO | GSSR | INAMBARI | INAMBARIE | PH | RONDONIA | TAPAJOS | XINGU |
|---|---|---|---|---|---|---|---|---|---|---|---|---|---|---|---|---|---|---|
| AFBAHIA | | 0.05<br>0.21 | 0.20<br>0.31 | 0.53<br>0.63 | 0.47<br>0.62 | 0.52<br>0.63 | 0.50<br>0.60 | 0.50<br>0.60 | 0.18<br>0.26 | 0.19<br>0.27 | 0.18<br>0.27 | 0.19<br>0.26 | 0.17<br>0.26 | 0.17<br>0.27 | 0.15<br>0.24 | 0.21<br>0.35 | 0.13<br>0.21 | 0.13<br>0.22 |
| AFES | 0.13 | | 0.02<br>0.17 | 0.63<br>0.74 | 0.61<br>0.74 | 0.61<br>0.72 | 0.58<br>0.68 | 0.53<br>0.65 | 0.20<br>0.31 | 0.21<br>0.33 | 0.20<br>0.31 | 0.20<br>0.31 | 0.19<br>0.31 | 0.20<br>0.33 | 0.17<br>0.28 | 0.28<br>0.43 | 0.15<br>0.26 | 0.14<br>0.27 |
| AFRIO | 0.26 | 0.09 | | 0.64<br>0.73 | 0.63<br>0.74 | 0.63<br>0.72 | 0.59<br>0.69 | 0.58<br>0.68 | 0.26<br>0.34 | 0.31<br>0.40 | 0.29<br>0.39 | 0.26<br>0.35 | 0.30<br>0.39 | 0.30<br>0.40 | 0.27<br>0.36 | 0.38<br>0.51 | 0.23<br>0.31 | 0.28<br>0.39 |
| CACR | 0.59 | 0.69 | 0.69 | | 0.56<br>0.71 | 0.01<br>0.11 | 0.60<br>0.71 | 0.56<br>0.66 | 0.34<br>0.44 | 0.36<br>0.47 | 0.34<br>0.45 | 0.34<br>0.44 | 0.38<br>0.48 | 0.37<br>0.48 | 0.31<br>0.42 | 0.50<br>0.62 | 0.36<br>0.46 | 0.45<br>0.55 |
| CAMA | 0.55 | 0.68 | 0.69 | 0.64 | | 0.56<br>0.71 | 0.48<br>0.64 | 0.47<br>0.62 | 0.29<br>0.42 | 0.29<br>0.43 | 0.26<br>0.41 | 0.28<br>0.42 | 0.24<br>0.39 | 0.29<br>0.43 | 0.21<br>0.36 | 0.30<br>0.49 | 0.28<br>0.42 | 0.33<br>0.48 |
| CAPA | 0.58 | 0.67 | 0.68 | 0.06 | 0.64 | | 0.58<br>0.69 | 0.55<br>0.65 | 0.32<br>0.43 | 0.35<br>0.45 | 0.33<br>0.43 | 0.32<br>0.42 | 0.38<br>0.47 | 0.37<br>0.46 | 0.30<br>0.40 | 0.50<br>0.62 | 0.35<br>0.45 | 0.44<br>0.54 |
| CPN | 0.55 | 0.63 | 0.64 | 0.66 | 0.57 | 0.64 | | 0.31<br>0.44 | 0.28<br>0.39 | 0.30<br>0.40 | 0.28<br>0.38 | 0.28<br>0.38 | 0.31<br>0.42 | 0.33<br>0.45 | 0.26<br>0.37 | 0.40<br>0.53 | 0.30<br>0.40 | 0.36<br>0.48 |
| CPS | 0.55 | 0.59 | 0.63 | 0.61 | 0.55 | 0.60 | 0.38 | | 0.29<br>0.40 | 0.32<br>0.42 | 0.31<br>0.41 | 0.31<br>0.40 | 0.34<br>0.44 | 0.36<br>0.47 | 0.29<br>0.39 | 0.40<br>0.53 | 0.33<br>0.43 | 0.39<br>0.49 |
| GS | 0.22 | 0.26 | 0.30 | 0.39 | 0.36 | 0.37 | 0.33 | 0.35 | | 0.00<br>0.01 | 0.01<br>0.02 | 0.01<br>0.02 | 0.11<br>0.16 | 0.09<br>0.14 | 0.06<br>0.10 | 0.09<br>0.17 | 0.11<br>0.15 | 0.13<br>0.19 |
| GSIMERI | 0.23 | 0.27 | 0.35 | 0.42 | 0.37 | 0.40 | 0.35 | 0.37 | 0.00* | | 0.01<br>0.03 | 0.01<br>0.03 | 0.10<br>0.15 | 0.09<br>0.15 | 0.05<br>0.10 | 0.09<br>0.18 | 0.10<br>0.15 | 0.12<br>0.19 |
| GSNAPO | 0.22 | 0.26 | 0.34 | 0.40 | 0.34 | 0.38 | 0.34 | 0.36 | 0.01 | 0.02 | | 0.02<br>0.04 | 0.09<br>0.14 | 0.08<br>0.13 | 0.04<br>0.08 | 0.06<br>0.13 | 0.09<br>0.14 | 0.11<br>0.18 |
| GSSR | 0.22 | 0.26 | 0.30 | 0.39 | 0.36 | 0.38 | 0.33 | 0.35 | 0.01 | 0.02 | 0.03 | | 0.11<br>0.16 | 0.10<br>0.16 | 0.06<br>0.10 | 0.09<br>0.17 | 0.11<br>0.16 | 0.14<br>0.20 |
| INAMBARI | 0.22 | 0.25 | 0.35 | 0.43 | 0.32 | 0.43 | 0.37 | 0.39 | 0.13 | 0.13 | 0.11 | 0.14 | | 0.04<br>0.10 | 0.04<br>0.08 | 0.03<br>0.11 | 0.07<br>0.12 | 0.10<br>0.15 |
| INAMBARIE | 0.22 | 0.27 | 0.35 | 0.43 | 0.36 | 0.42 | 0.39 | 0.42 | 0.12 | 0.12 | 0.10 | 0.13 | 0.07 | | 0.04<br>0.10 | 0.03<br>0.12 | 0.05<br>0.09 | 0.09<br>0.17 |
| PH | 0.19 | 0.22 | 0.32 | 0.37 | 0.29 | 0.35 | 0.32 | 0.34 | 0.08 | 0.07 | 0.06 | 0.08 | 0.06 | 0.07 | | 0.03<br>0.12 | 0.06<br>0.11 | 0.09<br>0.16 |
| RONDONIA | 0.28 | 0.36 | 0.45 | 0.56 | 0.40 | 0.57 | 0.46 | 0.47 | 0.13 | 0.13 | 0.10 | 0.13 | 0.07 | 0.07 | 0.07 | | 0.01<br>0.09 | 0.06<br>0.17 |
| TAPAJOS | 0.16 | 0.21 | 0.28 | 0.41 | 0.35 | 0.40 | 0.35 | 0.38 | 0.13 | 0.13 | 0.11 | 0.13 | 0.09 | 0.07 | 0.09 | 0.05 | | 0.04<br>0.09 |
| XINGU | 0.18 | 0.21 | 0.34 | 0.50 | 0.41 | 0.50 | 0.42 | 0.45 | 0.16 | 0.16 | 0.15 | 0.17 | 0.13 | 0.13 | 0.12 | 0.12 | 0.06 | |

FST
0.6
0.4
0.2

### Supplementary Figure 10d
### SNP dataset 2m

| | AFBAHIA | AFES | AFRIO | CACR | CAMA | CAPA | CPN | CPS | GS | GSIMERI | GSNAPO | GSSR | INAMBARI | INAMBARIE | PH | RONDONIA | TAPAJOS | XINGU |
|---|---|---|---|---|---|---|---|---|---|---|---|---|---|---|---|---|---|---|
| AFBAHIA | | 0.00 / 0.76 | 0.22 / 0.57 | 0.58 / 0.85 | 0.61 / 0.91 | 0.53 / 0.82 | 0.44 / 0.83 | 0.52 / 0.83 | 0.16 / 0.62 | 0.19 / 0.61 | 0.19 / 0.60 | 0.20 / 0.63 | 0.15 / 0.61 | 0.16 / 0.64 | 0.10 / 0.58 | 0.18 / 0.71 | 0.13 / 0.47 | 0.05 / 0.56 |
| AFES | 0.50* | | 0.00 / 0.56 | 0.62 / 0.89 | 0.75 / 0.95 | 0.57 / 0.86 | 0.57 / 0.90 | 0.55 / 0.85 | 0.31 / 0.72 | 0.30 / 0.68 | 0.29 / 0.68 | 0.32 / 0.74 | 0.26 / 0.69 | 0.31 / 0.76 | 0.23 / 0.67 | 0.36 / 0.79 | 0.21 / 0.59 | 0.19 / 0.67 |
| AFRIO | 0.44 | 0.29* | | 0.60 / 0.83 | 0.58 / 0.88 | 0.55 / 0.80 | 0.50 / 0.78 | 0.56 / 0.81 | 0.28 / 0.64 | 0.30 / 0.63 | 0.30 / 0.61 | 0.30 / 0.67 | 0.30 / 0.63 | 0.29 / 0.65 | 0.24 / 0.60 | 0.33 / 0.66 | 0.25 / 0.55 | 0.26 / 0.63 |
| CACR | 0.75 | 0.78 | 0.73 | | 0.32 / 0.78 | 0.00 / 0.09 | 0.43 / 0.79 | 0.49 / 0.78 | 0.25 / 0.64 | 0.20 / 0.59 | 0.22 / 0.62 | 0.26 / 0.65 | 0.27 / 0.64 | 0.28 / 0.67 | 0.21 / 0.62 | 0.27 / 0.71 | 0.24 / 0.59 | 0.29 / 0.69 |
| CAMA | 0.80 | 0.87 | 0.75 | 0.60 | | 0.28 / 0.74 | 0.29 / 0.86 | 0.37 / 0.79 | 0.12 / 0.67 | 0.01 / 0.57 | 0.03 / 0.59 | 0.09 / 0.66 | 0.11 / 0.62 | 0.16 / 0.70 | 0.03 / 0.62 | 0.11 / 0.73 | 0.15 / 0.58 | 0.24 / 0.70 |
| CAPA | 0.71 | 0.74 | 0.70 | 0.04* | 0.54 | | 0.36 / 0.73 | 0.44 / 0.75 | 0.23 / 0.59 | 0.18 / 0.52 | 0.21 / 0.56 | 0.22 / 0.61 | 0.23 / 0.59 | 0.25 / 0.60 | 0.16 / 0.56 | 0.21 / 0.65 | 0.21 / 0.54 | 0.28 / 0.62 |
| CPN | 0.68 | 0.76 | 0.65 | 0.64 | 0.63 | 0.58 | | 0.07 / 0.59 | 0.07 / 0.41 | 0.05 / 0.39 | 0.06 / 0.37 | 0.07 / 0.44 | 0.08 / 0.40 | 0.10 / 0.46 | 0.00 / 0.35 | 0.00 / 0.41 | 0.07 / 0.37 | 0.15 / 0.46 |
| CPS | 0.70 | 0.73 | 0.70 | 0.66 | 0.63 | 0.63 | 0.36 | | 0.22 / 0.66 | 0.18 / 0.61 | 0.16 / 0.61 | 0.21 / 0.67 | 0.23 / 0.61 | 0.23 / 0.68 | 0.15 / 0.62 | 0.15 / 0.66 | 0.20 / 0.59 | 0.26 / 0.67 |
| GS | 0.44 | 0.57 | 0.49 | 0.48 | 0.46 | 0.43 | 0.25 | 0.48 | | 0.00 / 0.05 | 0.00 / 0.05 | 0.00 / 0.02 | 0.03 / 0.28 | 0.00 / 0.15 | 0.02 / 0.15 | 0.00 / 0.18 | 0.10 / 0.32 | 0.08 / 0.27 |
| GSIMERI | 0.43 | 0.52 | 0.49 | 0.42 | 0.36 | 0.37 | 0.23 | 0.43 | 0.03* | | 0.00 / 0.05 | 0.01 / 0.06 | 0.03 / 0.24 | 0.03 / 0.12 | 0.02 / 0.14 | 0.00 / 0.21 | 0.10 / 0.29 | 0.10 / 0.25 |
| GSNAPO | 0.43 | 0.52 | 0.48 | 0.44 | 0.37 | 0.40 | 0.20 | 0.42 | 0.02* | 0.02* | | 0.00 / 0.05 | 0.05 / 0.22 | 0.01 / 0.15 | 0.01 / 0.11 | 0.00 / 0.12 | 0.09 / 0.28 | 0.09 / 0.23 |
| GSSR | 0.46 | 0.58 | 0.51 | 0.49 | 0.46 | 0.44 | 0.25 | 0.47 | 0.01* | 0.03 | 0.02* | | 0.04 / 0.27 | 0.02 / 0.17 | 0.02 / 0.15 | 0.01 / 0.19 | 0.09 / 0.31 | 0.09 / 0.29 |
| INAMBARI | 0.42 | 0.52 | 0.48 | 0.48 | 0.41 | 0.43 | 0.25 | 0.46 | 0.14 | 0.15 | 0.13 | 0.14 | | 0.02 / 0.21 | 0.00 / 0.18 | 0.00 / 0.12 | 0.04 / 0.27 | 0.02 / 0.24 |
| INAMBARIE | 0.45 | 0.58 | 0.50 | 0.51 | 0.49 | 0.45 | 0.27 | 0.49 | 0.08* | 0.08 | 0.07 | 0.09 | 0.10 | | 0.00 / 0.07 | 0.00 / 0.13 | 0.04 / 0.24 | 0.03 / 0.20 |
| PH | 0.38 | 0.50 | 0.45 | 0.45 | 0.40 | 0.39 | 0.17* | 0.42 | 0.08 | 0.08 | 0.06 | 0.08 | 0.07* | 0.01* | | 0.00 / 0.01 | 0.02 / 0.21 | 0.00 / 0.16 |
| RONDONIA | 0.51 | 0.62 | 0.52 | 0.53 | 0.48 | 0.47 | 0.18* | 0.45 | 0.09* | 0.09* | 0.06* | 0.11 | 0.02* | 0.04* | 0.00* | | 0.00 / 0.17 | 0.00 / 0.12 |
| TAPAJOS | 0.32 | 0.44 | 0.41 | 0.44 | 0.40 | 0.40 | 0.22 | 0.43 | 0.21 | 0.19 | 0.18 | 0.21 | 0.16 | 0.13 | 0.10 | 0.04* | | 0.02 / 0.22 |
| XINGU | 0.32 | 0.47 | 0.47 | 0.53 | 0.52 | 0.48 | 0.31 | 0.50 | 0.17 | 0.18 | 0.17 | 0.19 | 0.13 | 0.11 | 0.07* | 0.01* | 0.11 | |

FST
0.8
0.6
0.4
0.2
0.0

### Supplementary Figure 10e
#### SNP dataset 3

| | AFBAHIA | AFES | AFRIO | CACR | CAMA | CAPA | CPN | CPS | GS | GSIMERI | GSNAPO | GSSR | INAMBARI | INAMBARIE | PH | RONDONIA | TAPAJOS | XINGU |
|---|---|---|---|---|---|---|---|---|---|---|---|---|---|---|---|---|---|---|
| AFBAHIA | | 0.12<br>0.16 | 0.19<br>0.23 | 0.60<br>0.63 | 0.44<br>0.48 | 0.63<br>0.66 | 0.51<br>0.54 | 0.52<br>0.55 | 0.25<br>0.27 | 0.27<br>0.29 | 0.26<br>0.28 | 0.26<br>0.28 | 0.23<br>0.25 | 0.21<br>0.23 | 0.23<br>0.26 | 0.24<br>0.28 | 0.17<br>0.19 | 0.16<br>0.18 |
| AFES | 0.14 | | 0.14<br>0.18 | 0.76<br>0.79 | 0.57<br>0.62 | 0.78<br>0.81 | 0.64<br>0.67 | 0.62<br>0.66 | 0.27<br>0.30 | 0.29<br>0.32 | 0.28<br>0.31 | 0.28<br>0.31 | 0.26<br>0.29 | 0.25<br>0.28 | 0.27<br>0.30 | 0.31<br>0.36 | 0.21<br>0.23 | 0.21<br>0.25 |
| AFRIO | 0.21 | 0.16 | | 0.69<br>0.72 | 0.60<br>0.64 | 0.71<br>0.73 | 0.63<br>0.65 | 0.62<br>0.65 | 0.32<br>0.35 | 0.37<br>0.40 | 0.36<br>0.39 | 0.34<br>0.36 | 0.36<br>0.39 | 0.33<br>0.36 | 0.35<br>0.38 | 0.42<br>0.45 | 0.27<br>0.30 | 0.30<br>0.32 |
| CACR | 0.62 | 0.78 | 0.70 | | 0.68<br>0.73 | 0.04<br>0.09 | 0.72<br>0.75 | 0.67<br>0.70 | 0.33<br>0.35 | 0.37<br>0.39 | 0.36<br>0.38 | 0.34<br>0.37 | 0.40<br>0.43 | 0.40<br>0.43 | 0.36<br>0.39 | 0.58<br>0.61 | 0.37<br>0.40 | 0.49<br>0.52 |
| CAMA | 0.46 | 0.60 | 0.62 | 0.71 | | 0.71<br>0.75 | 0.56<br>0.61 | 0.55<br>0.59 | 0.17<br>0.21 | 0.19<br>0.22 | 0.17<br>0.20 | 0.18<br>0.22 | 0.17<br>0.22 | 0.20<br>0.24 | 0.15<br>0.19 | 0.20<br>0.26 | 0.21<br>0.25 | 0.28<br>0.32 |
| CAPA | 0.65 | 0.79 | 0.72 | 0.06 | 0.73 | | 0.74<br>0.77 | 0.69<br>0.72 | 0.34<br>0.36 | 0.39<br>0.41 | 0.38<br>0.41 | 0.35<br>0.38 | 0.44<br>0.47 | 0.43<br>0.46 | 0.38<br>0.41 | 0.63<br>0.66 | 0.39<br>0.42 | 0.52<br>0.55 |
| CPN | 0.53 | 0.65 | 0.64 | 0.74 | 0.59 | 0.75 | | 0.34<br>0.38 | 0.26<br>0.28 | 0.28<br>0.31 | 0.27<br>0.30 | 0.26<br>0.29 | 0.30<br>0.33 | 0.30<br>0.33 | 0.26<br>0.29 | 0.44<br>0.48 | 0.29<br>0.32 | 0.39<br>0.43 |
| CPS | 0.54 | 0.64 | 0.63 | 0.69 | 0.57 | 0.71 | 0.36 | | 0.27<br>0.30 | 0.31<br>0.33 | 0.30<br>0.33 | 0.28<br>0.31 | 0.33<br>0.36 | 0.33<br>0.36 | 0.29<br>0.32 | 0.45<br>0.49 | 0.31<br>0.34 | 0.42<br>0.45 |
| GS | 0.26 | 0.29 | 0.33 | 0.34 | 0.19 | 0.35 | 0.27 | 0.29 | | 0.00<br>0.01 | 0.01<br>0.01 | 0.01<br>0.02 | 0.11<br>0.13 | 0.12<br>0.13 | 0.07<br>0.08 | 0.11<br>0.13 | 0.14<br>0.15 | 0.18<br>0.20 |
| GSIMERI | 0.28 | 0.30 | 0.38 | 0.38 | 0.21 | 0.40 | 0.30 | 0.32 | 0.00* | | 0.01<br>0.02 | 0.02<br>0.03 | 0.12<br>0.14 | 0.13<br>0.14 | 0.08<br>0.09 | 0.12<br>0.14 | 0.15<br>0.17 | 0.20<br>0.22 |
| GSNAPO | 0.27 | 0.30 | 0.38 | 0.37 | 0.19 | 0.39 | 0.29 | 0.31 | 0.01 | 0.02 | | 0.03<br>0.03 | 0.10<br>0.12 | 0.11<br>0.13 | 0.06<br>0.07 | 0.10<br>0.12 | 0.14<br>0.15 | 0.18<br>0.20 |
| GSSR | 0.27 | 0.29 | 0.35 | 0.35 | 0.20 | 0.37 | 0.28 | 0.30 | 0.02 | 0.02 | 0.03 | | 0.12<br>0.14 | 0.13<br>0.14 | 0.08<br>0.09 | 0.12<br>0.15 | 0.15<br>0.17 | 0.19<br>0.21 |
| INAMBARI | 0.24 | 0.28 | 0.37 | 0.42 | 0.19 | 0.45 | 0.31 | 0.35 | 0.12 | 0.13 | 0.11 | 0.13 | | 0.04<br>0.06 | 0.06<br>0.07 | 0.04<br>0.06 | 0.07<br>0.09 | 0.13<br>0.15 |
| INAMBARIE | 0.22 | 0.26 | 0.34 | 0.41 | 0.22 | 0.44 | 0.32 | 0.34 | 0.12 | 0.14 | 0.12 | 0.14 | 0.05 | | 0.07<br>0.08 | 0.02<br>0.04 | 0.05<br>0.07 | 0.12<br>0.14 |
| PH | 0.25 | 0.29 | 0.36 | 0.37 | 0.17 | 0.40 | 0.28 | 0.30 | 0.07 | 0.08 | 0.07 | 0.08 | 0.06 | 0.08 | | 0.06<br>0.09 | 0.10<br>0.11 | 0.15<br>0.17 |
| RONDONIA | 0.26 | 0.33 | 0.43 | 0.60 | 0.23 | 0.64 | 0.46 | 0.47 | 0.12 | 0.13 | 0.11 | 0.13 | 0.05 | 0.03 | 0.08 | | 0.03<br>0.05 | 0.10<br>0.12 |
| TAPAJOS | 0.18 | 0.22 | 0.29 | 0.39 | 0.23 | 0.40 | 0.31 | 0.32 | 0.15 | 0.16 | 0.14 | 0.16 | 0.08 | 0.06 | 0.10 | 0.04 | | 0.07<br>0.08 |
| XINGU | 0.17 | 0.23 | 0.31 | 0.50 | 0.30 | 0.53 | 0.41 | 0.43 | 0.19 | 0.21 | 0.19 | 0.20 | 0.14 | 0.13 | 0.16 | 0.11 | 0.07 | |

FST
0.6
0.4
0.2

### Supplementary Figure 10f
### SNP dataset 3m

| | AFBAHIA | AFES | AFRIO | CACR | CAMA | CAPA | CPN | CPS | GS | GSIMERI | GSNAPO | GSSR | INAMBARI | INAMBARIE | PH | RONDONIA | TAPAJOS | XINGU |
|---|---|---|---|---|---|---|---|---|---|---|---|---|---|---|---|---|---|---|
| AFBAHIA | | 0.05<br>0.14 | 0.14<br>0.24 | 0.55<br>0.65 | 0.38<br>0.52 | 0.58<br>0.67 | 0.46<br>0.56 | 0.46<br>0.56 | 0.24<br>0.33 | 0.27<br>0.36 | 0.27<br>0.35 | 0.25<br>0.34 | 0.25<br>0.33 | 0.21<br>0.32 | 0.24<br>0.33 | 0.19<br>0.31 | 0.17<br>0.25 | 0.12<br>0.19 |
| AFES | 0.09 | | 0.05<br>0.17 | 0.71<br>0.81 | 0.48<br>0.64 | 0.74<br>0.82 | 0.60<br>0.70 | 0.55<br>0.66 | 0.28<br>0.37 | 0.31<br>0.40 | 0.30<br>0.39 | 0.29<br>0.39 | 0.27<br>0.38 | 0.24<br>0.35 | 0.28<br>0.37 | 0.23<br>0.37 | 0.21<br>0.29 | 0.18<br>0.28 |
| AFRIO | 0.19 | 0.11 | | 0.64<br>0.74 | 0.56<br>0.68 | 0.66<br>0.75 | 0.59<br>0.68 | 0.57<br>0.66 | 0.34<br>0.42 | 0.39<br>0.47 | 0.38<br>0.47 | 0.35<br>0.44 | 0.38<br>0.48 | 0.34<br>0.44 | 0.37<br>0.46 | 0.37<br>0.49 | 0.27<br>0.36 | 0.27<br>0.36 |
| CACR | 0.60 | 0.76 | 0.69 | | 0.66<br>0.80 | 0.01<br>0.14 | 0.72<br>0.82 | 0.64<br>0.74 | 0.27<br>0.36 | 0.33<br>0.42 | 0.31<br>0.42 | 0.30<br>0.39 | 0.36<br>0.47 | 0.36<br>0.46 | 0.32<br>0.42 | 0.51<br>0.63 | 0.32<br>0.41 | 0.45<br>0.55 |
| CAMA | 0.46 | 0.56 | 0.62 | 0.73 | | 0.66<br>0.81 | 0.59<br>0.74 | 0.47<br>0.64 | 0.12<br>0.24 | 0.14<br>0.25 | 0.14<br>0.26 | 0.12<br>0.25 | 0.12<br>0.26 | 0.15<br>0.28 | 0.10<br>0.23 | 0.10<br>0.30 | 0.16<br>0.28 | 0.21<br>0.35 |
| CAPA | 0.62 | 0.78 | 0.71 | 0.08 | 0.73 | | 0.73<br>0.82 | 0.64<br>0.75 | 0.28<br>0.37 | 0.34<br>0.44 | 0.32<br>0.42 | 0.31<br>0.40 | 0.38<br>0.48 | 0.38<br>0.48 | 0.34<br>0.44 | 0.54<br>0.66 | 0.33<br>0.43 | 0.47<br>0.57 |
| CPN | 0.51 | 0.65 | 0.64 | 0.77 | 0.67 | 0.78 | | 0.25<br>0.41 | 0.24<br>0.33 | 0.28<br>0.37 | 0.28<br>0.37 | 0.26<br>0.36 | 0.27<br>0.37 | 0.27<br>0.37 | 0.25<br>0.35 | 0.40<br>0.51 | 0.25<br>0.34 | 0.35<br>0.46 |
| CPS | 0.51 | 0.61 | 0.62 | 0.69 | 0.56 | 0.70 | 0.33 | | 0.26<br>0.34 | 0.30<br>0.39 | 0.29<br>0.37 | 0.27<br>0.37 | 0.29<br>0.39 | 0.29<br>0.39 | 0.25<br>0.34 | 0.38<br>0.50 | 0.25<br>0.34 | 0.38<br>0.47 |
| GS | 0.29 | 0.33 | 0.38 | 0.32 | 0.18 | 0.33 | 0.29 | 0.30 | | 0.00<br>0.01 | 0.01<br>0.02 | 0.01<br>0.02 | 0.09<br>0.15 | 0.10<br>0.16 | 0.07<br>0.11 | 0.07<br>0.15 | 0.12<br>0.17 | 0.16<br>0.23 |
| GSIMERI | 0.31 | 0.35 | 0.43 | 0.38 | 0.20 | 0.39 | 0.33 | 0.34 | 0.00* | | 0.01<br>0.02 | 0.02<br>0.03 | 0.11<br>0.18 | 0.12<br>0.18 | 0.07<br>0.12 | 0.08<br>0.16 | 0.13<br>0.18 | 0.18<br>0.25 |
| GSNAPO | 0.31 | 0.35 | 0.43 | 0.37 | 0.20 | 0.38 | 0.32 | 0.33 | 0.01 | 0.01 | | 0.02<br>0.05 | 0.10<br>0.16 | 0.11<br>0.17 | 0.05<br>0.10 | 0.07<br>0.14 | 0.12<br>0.17 | 0.18<br>0.25 |
| GSSR | 0.30 | 0.34 | 0.40 | 0.35 | 0.19 | 0.35 | 0.31 | 0.32 | 0.02 | 0.03 | 0.04 | | 0.11<br>0.17 | 0.12<br>0.18 | 0.08<br>0.13 | 0.09<br>0.17 | 0.14<br>0.19 | 0.17<br>0.24 |
| INAMBARI | 0.29 | 0.33 | 0.43 | 0.42 | 0.19 | 0.44 | 0.32 | 0.34 | 0.12 | 0.14 | 0.13 | 0.14 | | 0.04<br>0.10 | 0.04<br>0.09 | 0.04<br>0.12 | 0.06<br>0.12 | 0.12<br>0.20 |
| INAMBARIE | 0.26 | 0.29 | 0.39 | 0.41 | 0.21 | 0.43 | 0.33 | 0.34 | 0.13 | 0.15 | 0.14 | 0.15 | 0.07 | | 0.08<br>0.14 | 0.02<br>0.09 | 0.06<br>0.11 | 0.12<br>0.19 |
| PH | 0.29 | 0.33 | 0.42 | 0.37 | 0.17 | 0.39 | 0.30 | 0.30 | 0.09 | 0.09 | 0.07 | 0.11 | 0.07 | 0.11 | | 0.01<br>0.08 | 0.08<br>0.13 | 0.15<br>0.22 |
| RONDONIA | 0.25 | 0.30 | 0.43 | 0.57 | 0.21 | 0.60 | 0.46 | 0.44 | 0.11 | 0.12 | 0.11 | 0.13 | 0.08 | 0.05 | 0.05 | | 0.01<br>0.09 | 0.08<br>0.17 |
| TAPAJOS | 0.21 | 0.25 | 0.32 | 0.37 | 0.22 | 0.38 | 0.29 | 0.30 | 0.14 | 0.15 | 0.14 | 0.16 | 0.09 | 0.08 | 0.10 | 0.05 | | 0.06<br>0.11 |
| XINGU | 0.15 | 0.23 | 0.32 | 0.50 | 0.29 | 0.52 | 0.41 | 0.43 | 0.20 | 0.21 | 0.21 | 0.20 | 0.16 | 0.16 | 0.18 | 0.12 | 0.09 | |

FST
0.6
0.4
0.2

### Supplementary Figure 10g
### SNP dataset 4

| | AFBAHIA | AFES | AFRIO | CACR | CAMA | CAPA | CPN | CPS | GS | GSIMERI | GSNAPO | GSSR | INAMBARI | INAMBARIE | PH | RONDONIA | TAPAJOS | XINGU |
|---|---|---|---|---|---|---|---|---|---|---|---|---|---|---|---|---|---|---|
| AFBAHIA | | 0.11<br>0.15 | 0.19<br>0.22 | 0.57<br>0.59 | 0.47<br>0.50 | 0.58<br>0.60 | 0.50<br>0.52 | 0.46<br>0.49 | 0.22<br>0.24 | 0.24<br>0.26 | 0.23<br>0.25 | 0.23<br>0.25 | 0.19<br>0.21 | 0.18<br>0.20 | 0.19<br>0.21 | 0.23<br>0.25 | 0.14<br>0.16 | 0.15<br>0.17 |
| AFES | 0.13 | | 0.13<br>0.16 | 0.67<br>0.70 | 0.58<br>0.62 | 0.66<br>0.69 | 0.57<br>0.60 | 0.50<br>0.53 | 0.24<br>0.27 | 0.26<br>0.29 | 0.24<br>0.27 | 0.24<br>0.27 | 0.20<br>0.23 | 0.21<br>0.23 | 0.20<br>0.23 | 0.27<br>0.31 | 0.17<br>0.19 | 0.19<br>0.21 |
| AFRIO | 0.21 | 0.15 | | 0.67<br>0.69 | 0.63<br>0.66 | 0.67<br>0.69 | 0.61<br>0.64 | 0.57<br>0.59 | 0.29<br>0.32 | 0.34<br>0.36 | 0.33<br>0.35 | 0.31<br>0.33 | 0.31<br>0.34 | 0.30<br>0.32 | 0.30<br>0.32 | 0.40<br>0.43 | 0.24<br>0.26 | 0.28<br>0.31 |
| CACR | 0.58 | 0.69 | 0.68 | | 0.60<br>0.63 | 0.05<br>0.08 | 0.64<br>0.67 | 0.58<br>0.60 | 0.37<br>0.40 | 0.41<br>0.43 | 0.39<br>0.41 | 0.38<br>0.40 | 0.39<br>0.42 | 0.41<br>0.43 | 0.36<br>0.38 | 0.52<br>0.55 | 0.39<br>0.41 | 0.48<br>0.50 |
| CAMA | 0.49 | 0.60 | 0.64 | 0.62 | | 0.60<br>0.63 | 0.52<br>0.56 | 0.47<br>0.50 | 0.28<br>0.31 | 0.29<br>0.32 | 0.27<br>0.30 | 0.28<br>0.31 | 0.23<br>0.26 | 0.27<br>0.30 | 0.21<br>0.24 | 0.26<br>0.31 | 0.28<br>0.32 | 0.34<br>0.38 |
| CAPA | 0.59 | 0.68 | 0.68 | 0.06 | 0.61 | | 0.64<br>0.67 | 0.59<br>0.61 | 0.38<br>0.40 | 0.41<br>0.44 | 0.40<br>0.43 | 0.39<br>0.41 | 0.41<br>0.44 | 0.42<br>0.45 | 0.37<br>0.40 | 0.54<br>0.57 | 0.40<br>0.42 | 0.49<br>0.51 |
| CPN | 0.51 | 0.58 | 0.63 | 0.66 | 0.54 | 0.65 | | 0.34<br>0.37 | 0.31<br>0.34 | 0.33<br>0.36 | 0.32<br>0.34 | 0.32<br>0.34 | 0.31<br>0.34 | 0.33<br>0.36 | 0.28<br>0.31 | 0.42<br>0.44 | 0.33<br>0.35 | 0.40<br>0.43 |
| CPS | 0.48 | 0.52 | 0.58 | 0.59 | 0.48 | 0.60 | 0.36 | | 0.30<br>0.33 | 0.32<br>0.35 | 0.31<br>0.33 | 0.31<br>0.33 | 0.31<br>0.33 | 0.32<br>0.34 | 0.28<br>0.30 | 0.36<br>0.39 | 0.31<br>0.34 | 0.38<br>0.41 |
| GS | 0.23 | 0.26 | 0.30 | 0.38 | 0.29 | 0.39 | 0.33 | 0.32 | | 0.00<br>0.01 | 0.01<br>0.01 | 0.01<br>0.02 | 0.10<br>0.12 | 0.10<br>0.11 | 0.07<br>0.07 | 0.11<br>0.13 | 0.12<br>0.13 | 0.16<br>0.18 |
| GSIMERI | 0.25 | 0.27 | 0.35 | 0.42 | 0.31 | 0.43 | 0.35 | 0.34 | 0.00* | | 0.01<br>0.02 | 0.02<br>0.02 | 0.11<br>0.12 | 0.11<br>0.12 | 0.07<br>0.08 | 0.12<br>0.14 | 0.12<br>0.13 | 0.17<br>0.19 |
| GSNAPO | 0.24 | 0.26 | 0.34 | 0.40 | 0.28 | 0.42 | 0.33 | 0.32 | 0.01 | 0.02 | | 0.03<br>0.03 | 0.09<br>0.10 | 0.10<br>0.11 | 0.05<br>0.06 | 0.10<br>0.12 | 0.11<br>0.12 | 0.16<br>0.18 |
| GSSR | 0.24 | 0.26 | 0.32 | 0.39 | 0.30 | 0.40 | 0.33 | 0.32 | 0.01 | 0.02 | 0.03 | | 0.11<br>0.12 | 0.11<br>0.12 | 0.07<br>0.08 | 0.12<br>0.14 | 0.13<br>0.14 | 0.16<br>0.18 |
| INAMBARI | 0.20 | 0.21 | 0.32 | 0.40 | 0.25 | 0.42 | 0.32 | 0.32 | 0.11 | 0.11 | 0.10 | 0.12 | | 0.04<br>0.05 | 0.05<br>0.06 | 0.03<br>0.05 | 0.07<br>0.08 | 0.11<br>0.13 |
| INAMBARIE | 0.19 | 0.22 | 0.31 | 0.42 | 0.29 | 0.43 | 0.34 | 0.33 | 0.11 | 0.12 | 0.10 | 0.12 | 0.05 | | 0.06<br>0.07 | 0.03<br>0.05 | 0.05<br>0.06 | 0.10<br>0.12 |
| PH | 0.20 | 0.21 | 0.31 | 0.37 | 0.22 | 0.39 | 0.29 | 0.29 | 0.07 | 0.07 | 0.06 | 0.07 | 0.05 | 0.07 | | 0.05<br>0.07 | 0.08<br>0.09 | 0.12<br>0.14 |
| RONDONIA | 0.24 | 0.29 | 0.42 | 0.53 | 0.28 | 0.55 | 0.43 | 0.38 | 0.12 | 0.13 | 0.11 | 0.13 | 0.04 | 0.04 | 0.06 | | 0.03<br>0.05 | 0.09<br>0.12 |
| TAPAJOS | 0.15 | 0.18 | 0.25 | 0.40 | 0.30 | 0.41 | 0.34 | 0.33 | 0.12 | 0.13 | 0.12 | 0.13 | 0.07 | 0.05 | 0.09 | 0.04 | | 0.06<br>0.07 |
| XINGU | 0.16 | 0.20 | 0.29 | 0.49 | 0.36 | 0.50 | 0.41 | 0.40 | 0.17 | 0.18 | 0.17 | 0.17 | 0.12 | 0.11 | 0.13 | 0.11 | 0.07 | |

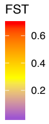

#### Supplementary Figure 10h
#### SNP dataset 4m

| | AFBAHIA | AFES | AFRIO | CACR | CAMA | CAPA | CPN | CPS | GS | GSIMERI | GSNAPO | GSSR | INAMBARI | INAMBARIE | PH | RONDONIA | TAPAJOS | XINGU |
|---|---|---|---|---|---|---|---|---|---|---|---|---|---|---|---|---|---|---|
| AFBAHIA | | 0.07<br>0.15 | 0.16<br>0.24 | 0.54<br>0.62 | 0.44<br>0.54 | 0.55<br>0.62 | 0.47<br>0.55 | 0.43<br>0.51 | 0.21<br>0.28 | 0.24<br>0.31 | 0.22<br>0.29 | 0.22<br>0.29 | 0.19<br>0.26 | 0.18<br>0.25 | 0.19<br>0.26 | 0.18<br>0.27 | 0.14<br>0.20 | 0.11<br>0.17 |
| AFES | 0.11 | | 0.07<br>0.18 | 0.64<br>0.72 | 0.55<br>0.67 | 0.64<br>0.73 | 0.55<br>0.63 | 0.44<br>0.53 | 0.24<br>0.33 | 0.27<br>0.36 | 0.25<br>0.33 | 0.25<br>0.34 | 0.20<br>0.29 | 0.19<br>0.28 | 0.19<br>0.29 | 0.21<br>0.32 | 0.15<br>0.24 | 0.15<br>0.24 |
| AFRIO | 0.20 | 0.12 | | 0.64<br>0.71 | 0.60<br>0.69 | 0.64<br>0.71 | 0.60<br>0.67 | 0.54<br>0.61 | 0.29<br>0.37 | 0.34<br>0.42 | 0.33<br>0.40 | 0.30<br>0.38 | 0.32<br>0.40 | 0.29<br>0.37 | 0.30<br>0.38 | 0.36<br>0.45 | 0.23<br>0.30 | 0.25<br>0.33 |
| CACR | 0.58 | 0.68 | 0.67 | | 0.58<br>0.69 | 0.02<br>0.09 | 0.65<br>0.72 | 0.55<br>0.63 | 0.34<br>0.42 | 0.39<br>0.47 | 0.36<br>0.45 | 0.35<br>0.44 | 0.37<br>0.46 | 0.38<br>0.47 | 0.34<br>0.42 | 0.47<br>0.56 | 0.36<br>0.43 | 0.45<br>0.53 |
| CAMA | 0.49 | 0.61 | 0.65 | 0.64 | | 0.59<br>0.69 | 0.55<br>0.66 | 0.43<br>0.53 | 0.24<br>0.35 | 0.26<br>0.38 | 0.24<br>0.35 | 0.25<br>0.36 | 0.20<br>0.31 | 0.23<br>0.35 | 0.18<br>0.29 | 0.18<br>0.33 | 0.23<br>0.34 | 0.28<br>0.39 |
| CAPA | 0.59 | 0.69 | 0.68 | 0.05 | 0.64 | | 0.64<br>0.72 | 0.57<br>0.65 | 0.35<br>0.43 | 0.39<br>0.48 | 0.37<br>0.46 | 0.36<br>0.45 | 0.38<br>0.47 | 0.40<br>0.48 | 0.35<br>0.44 | 0.49<br>0.58 | 0.36<br>0.45 | 0.46<br>0.54 |
| CPN | 0.51 | 0.59 | 0.63 | 0.68 | 0.61 | 0.69 | | 0.33<br>0.42 | 0.32<br>0.40 | 0.35<br>0.44 | 0.33<br>0.41 | 0.33<br>0.41 | 0.31<br>0.39 | 0.32<br>0.41 | 0.29<br>0.37 | 0.39<br>0.48 | 0.30<br>0.38 | 0.38<br>0.46 |
| CPS | 0.47 | 0.49 | 0.58 | 0.60 | 0.48 | 0.61 | 0.38 | | 0.31<br>0.39 | 0.34<br>0.41 | 0.31<br>0.39 | 0.33<br>0.40 | 0.30<br>0.37 | 0.31<br>0.39 | 0.27<br>0.35 | 0.32<br>0.41 | 0.29<br>0.37 | 0.36<br>0.44 |
| GS | 0.24 | 0.29 | 0.33 | 0.38 | 0.30 | 0.39 | 0.36 | 0.35 | | 0.00<br>0.01 | 0.01<br>0.02 | 0.01<br>0.02 | 0.08<br>0.13 | 0.09<br>0.13 | 0.06<br>0.09 | 0.08<br>0.14 | 0.10<br>0.14 | 0.15<br>0.20 |
| GSIMERI | 0.27 | 0.31 | 0.38 | 0.43 | 0.32 | 0.44 | 0.39 | 0.37 | 0.00* | | 0.01<br>0.02 | 0.02<br>0.03 | 0.09<br>0.15 | 0.10<br>0.15 | 0.06<br>0.10 | 0.09<br>0.15 | 0.10<br>0.14 | 0.16<br>0.22 |
| GSNAPO | 0.26 | 0.29 | 0.37 | 0.40 | 0.30 | 0.41 | 0.37 | 0.35 | 0.01 | 0.01 | | 0.02<br>0.04 | 0.08<br>0.13 | 0.09<br>0.14 | 0.04<br>0.07 | 0.07<br>0.12 | 0.10<br>0.13 | 0.15<br>0.20 |
| GSSR | 0.25 | 0.29 | 0.35 | 0.40 | 0.30 | 0.41 | 0.37 | 0.36 | 0.01 | 0.02 | 0.03 | | 0.10<br>0.14 | 0.10<br>0.15 | 0.07<br>0.11 | 0.09<br>0.16 | 0.11<br>0.15 | 0.15<br>0.21 |
| INAMBARI | 0.23 | 0.25 | 0.36 | 0.42 | 0.26 | 0.43 | 0.35 | 0.34 | 0.10 | 0.12 | 0.10 | 0.12 | | 0.04<br>0.08 | 0.04<br>0.07 | 0.03<br>0.09 | 0.05<br>0.10 | 0.10<br>0.16 |
| INAMBARIE | 0.21 | 0.24 | 0.33 | 0.43 | 0.29 | 0.44 | 0.37 | 0.35 | 0.11 | 0.12 | 0.11 | 0.13 | 0.05 | | 0.06<br>0.11 | 0.02<br>0.08 | 0.05<br>0.09 | 0.10<br>0.16 |
| PH | 0.22 | 0.24 | 0.34 | 0.38 | 0.24 | 0.40 | 0.33 | 0.31 | 0.08 | 0.08 | 0.06 | 0.09 | 0.05 | 0.08 | | 0.01<br>0.07 | 0.07<br>0.10 | 0.12<br>0.17 |
| RONDONIA | 0.23 | 0.27 | 0.40 | 0.52 | 0.26 | 0.54 | 0.43 | 0.36 | 0.11 | 0.12 | 0.10 | 0.12 | 0.06 | 0.05 | 0.04 | | 0.02<br>0.07 | 0.07<br>0.14 |
| TAPAJOS | 0.17 | 0.20 | 0.27 | 0.40 | 0.29 | 0.40 | 0.34 | 0.33 | 0.12 | 0.12 | 0.11 | 0.13 | 0.07 | 0.07 | 0.09 | 0.04 | | 0.06<br>0.09 |
| XINGU | 0.14 | 0.20 | 0.29 | 0.49 | 0.34 | 0.50 | 0.42 | 0.40 | 0.17 | 0.19 | 0.18 | 0.18 | 0.13 | 0.13 | 0.15 | 0.11 | 0.08 | |

FST: 0.6, 0.4, 0.2

**Supplementary Figure 11. EEMS model fit**

Regressing the observed dissimilarity between demes against the fitted dissimilarity between demes provides an indication of model fit (Petkova et al., 2015). For the present dataset, model fit (leftmost plot) was very high ($R^2$ = 0.865). Within demes (central plot, within demes that represent more than a single individual), model fit was somewhat less, but still high ($R^2$ = 0.579). Lastly, comparing observed dissimilarity between demes against great circle distance between demes suggested a strong signal of isolation by distance operating at the scale of the entire dataset ($R^2$ = 0.431). Overall, the EEMS model does a very good job of describing spatially structured variation in this dataset.

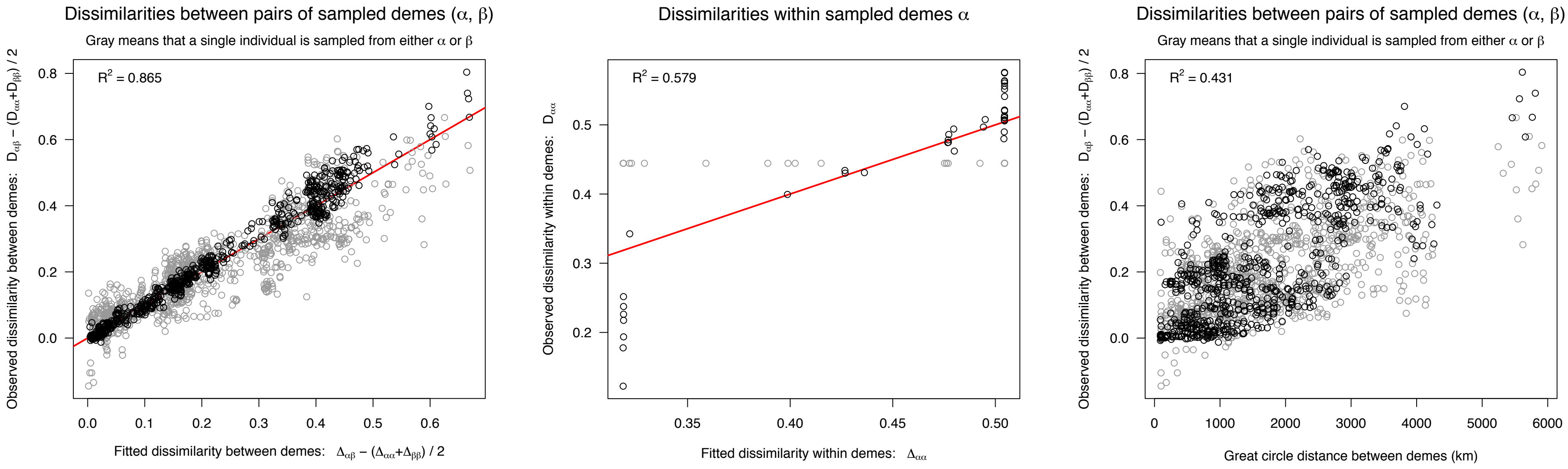
Dissimilarities between pairs of sampled demes (α, β)
Gray means that a single individual is sampled from either α or β
$R^2$ = 0.865
Observed dissimilarity between demes: $D_{\alpha\beta} - (D_{\alpha\alpha}+D_{\beta\beta}) / 2$
Fitted dissimilarity between demes: $\Delta_{\alpha\beta} - (\Delta_{\alpha\alpha}+\Delta_{\beta\beta}) / 2$
Dissimilarities within sampled demes α
$R^2$ = 0.579
Observed dissimilarity within demes: $D_{\alpha\alpha}$
Fitted dissimilarity within demes: $\Delta_{\alpha\alpha}$
Dissimilarities between pairs of sampled demes (α, β)
Gray means that a single individual is sampled from either α or β
$R^2$ = 0.431
Observed dissimilarity between demes: $D_{\alpha\beta} - (D_{\alpha\alpha}+D_{\beta\beta}) / 2$
Great circle distance between demes (km)

**Supplementary Figure 12. EEMS estimated genetic diversity**
Two broad clusters of relatively high genetic diversity (greatest heterozygosity) were detected in EEMS. The first is centered along the Amazon river and was relatively uniform within the northern Amazonian basin, reflecting relatively high diversity in the Guiana shield and lowland Amazonian populations. The second relatively high diversity group reflected the introgressed western Napo population. Areas of relatively low genetic diversity included the Atlantic Forest, the Peruvian Andes, and Central American lineages. These results are generally consistent with our estimates of allelic richness (Supplementary Appendix for details).

### Posterior mean diversity rates q (on the log10 scale)

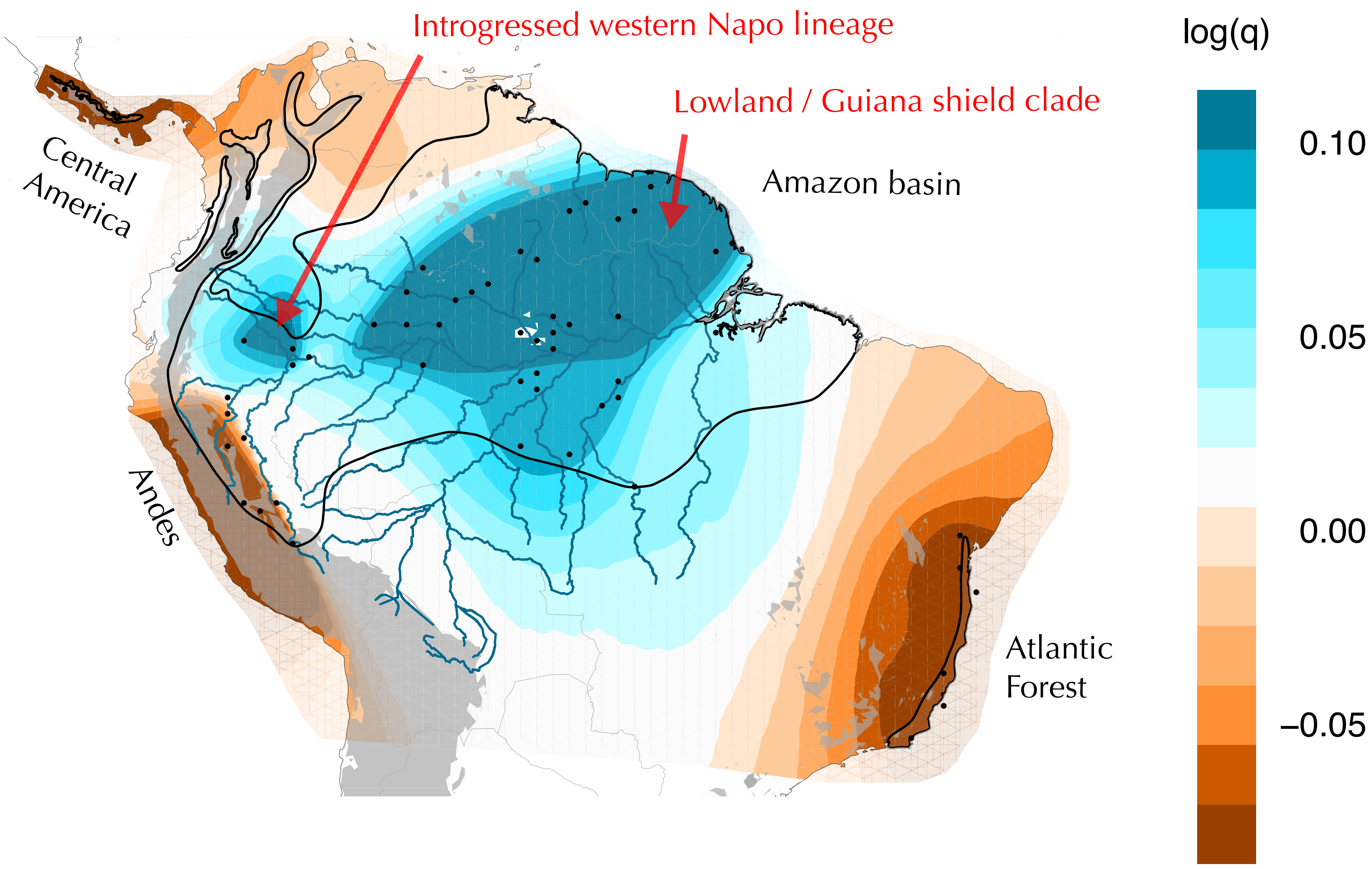

**Supplementary Figure 13. Quantification of vocal variation**
All *Pseudopipra* vocalizations have 1-3 buzzy and/or tonal notes. We measured: 1) starting frequency, 2) ending frequency, 3) minimum frequency, 4) maximum frequency, 5) number of notes, and 6) duration of the entire song. To obtain a conservative estimate of the number of individuals sampled, we took measurements of one song from each recording.

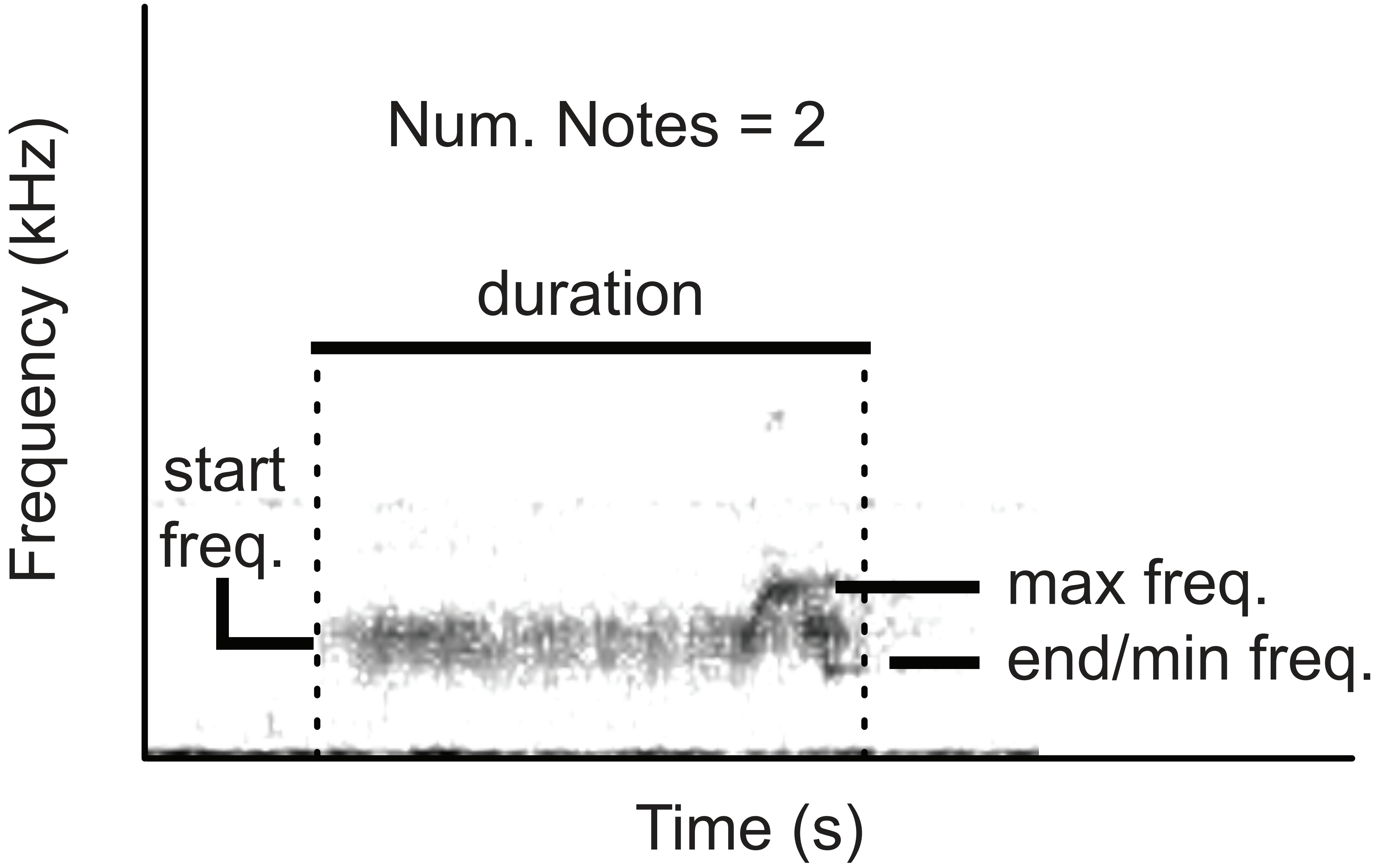
Num. Notes = 2
duration
start freq.
max freq.
end/min freq.
Frequency (kHz)
Time (s)

**Supplementary Figure 14. Expanded summary of vocalization phenotypes**
PCA and logistic regression on vocalizations (Supplementary Table 3) found significant differences among types ($p < 0.001$). The first three axes of a PCA explained ~90% of the variation in lekking vocalization characters, with PC1 (~64%) primarily explaining variation in note number and frequency. Panels A and D show vocalization records plotted into the first and second components of a principal components analysis. Panels B and E show vocalization data points plotted into the second and third principal components. Lastly, Panels C and F are reproduced from Figure 8 and 9 in the main text. All vocal types we identify can be quantitatively discriminated.

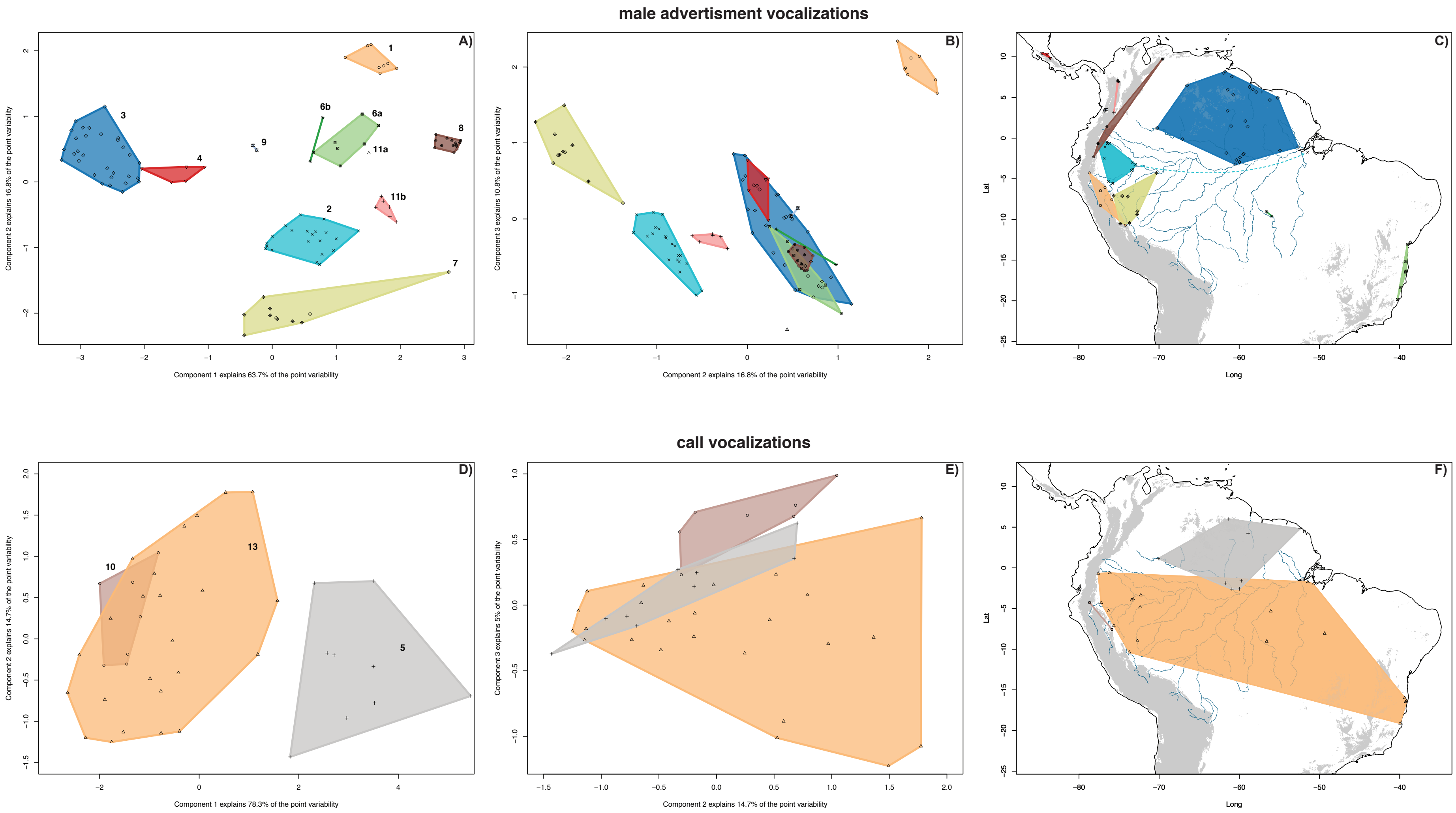
male advertisment vocalizations
A)
Component 1 explains 63.7% of the point variability
Component 2 explains 16.8% of the point variability
1
3
4
6b
6a
8
9
11a
11b
2
7
B)
Component 2 explains 16.8% of the point variability
Component 3 explains 10.8% of the point variability
C)
Long
Lat
call vocalizations
D)
Component 1 explains 78.3% of the point variability
Component 2 explains 14.7% of the point variability
10
13
5
E)
Component 2 explains 14.7% of the point variability
Component 3 explains 5% of the point variability
F)
Long
Lat

**Supplementary Figure 15. Vocalization loading plots**
PCA plots from Figure 14, shown here with loading vectors projected into principal component space. The direction and length of the vectors indicate the direction and strength of the variation in a particular direction. Panels A and B show data for lek vocalizations, while C and D show data for call vocalizations.

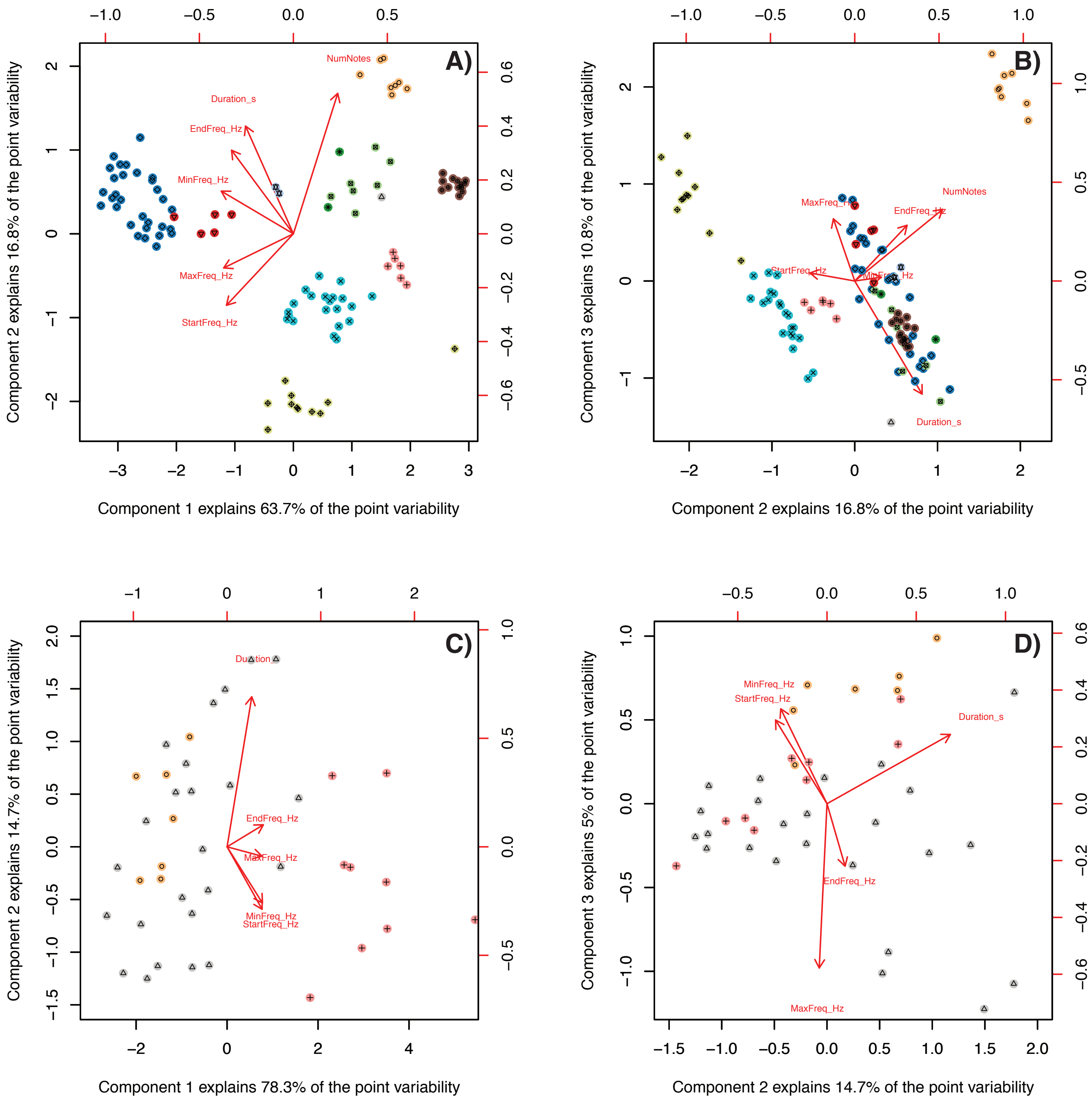
A)
NumNotes
Duration_s
EndFreq_Hz
MinFreq_Hz
MaxFreq_Hz
StartFreq_Hz
Component 1 explains 63.7% of the point variability
Component 2 explains 16.8% of the point variability
B)
MaxFreq_Hz
EndFreq_Hz
NumNotes
StartFreq_Hz
MinFreq_Hz
Duration_s
Component 2 explains 16.8% of the point variability
Component 3 explains 10.8% of the point variability
C)
Duration
EndFreq_Hz
MaxFreq_Hz
MinFreq_Hz
StartFreq_Hz
Component 1 explains 78.3% of the point variability
Component 2 explains 14.7% of the point variability
D)
MinFreq_Hz
StartFreq_Hz
Duration_s
EndFreq_Hz
MaxFreq_Hz
Component 2 explains 14.7% of the point variability
Component 3 explains 5% of the point variability

**Supplementary Figure 16 – Preliminary parsimony reconstruction of vocal types**
See description in Supplemental Appendix. Panel A shows a parsimony reconstruction of call vocal types, whereas Panel B shows a similar reconstruction for male advertisement identities. Black markers at tips indicate estimated states based on parsimony optimization.

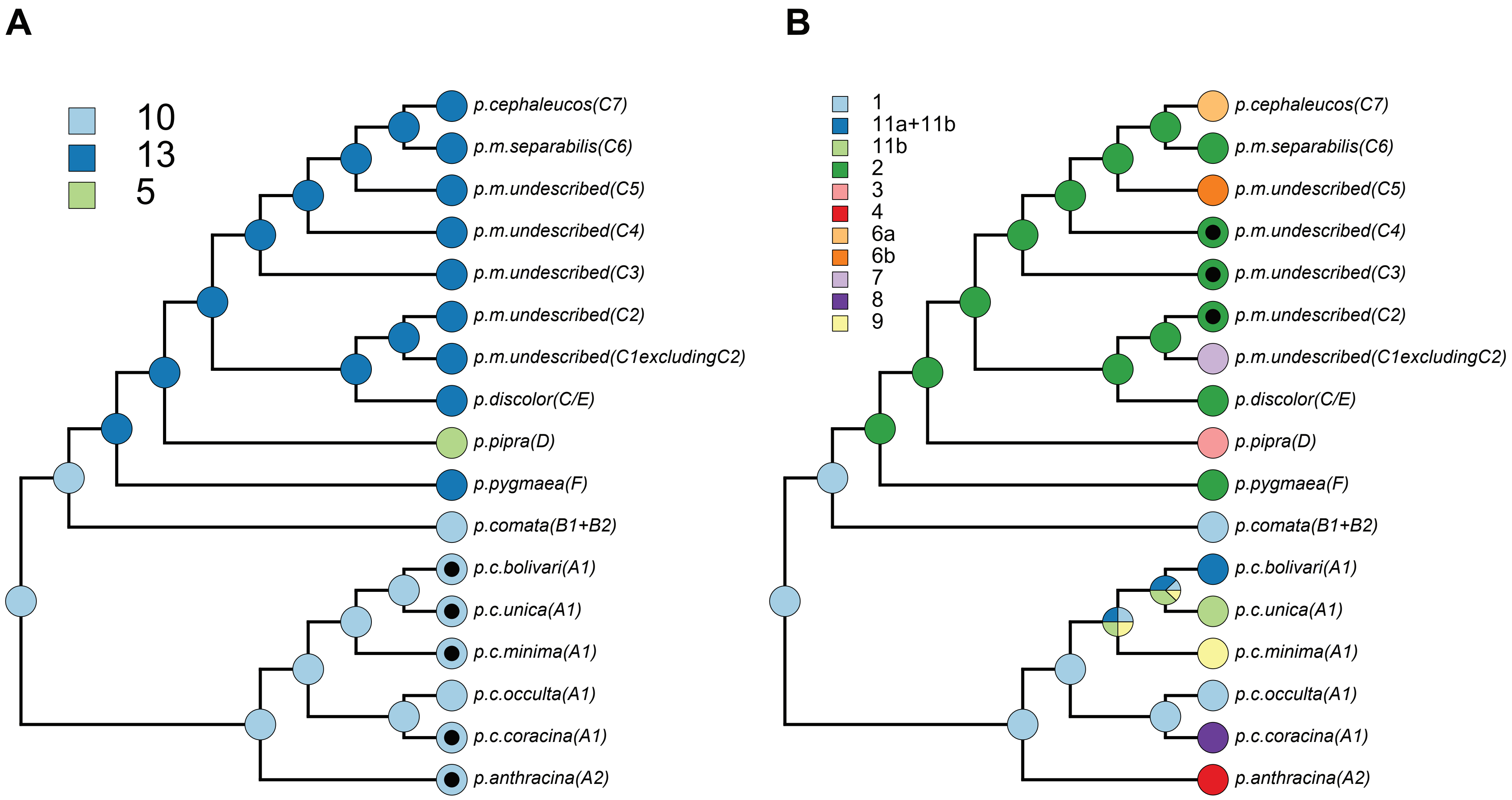
A
10
13
5
p.cephaleucos(C7)
p.m.separabilis(C6)
p.m.undescribed(C5)
p.m.undescribed(C4)
p.m.undescribed(C3)
p.m.undescribed(C2)
p.m.undescribed(C1excludingC2)
p.discolor(C/E)
p.pipra(D)
p.pygmaea(F)
p.comata(B1+B2)
p.c.bolivari(A1)
p.c.unica(A1)
p.c.minima(A1)
p.c.occulta(A1)
p.c.coracina(A1)
p.anthracina(A2)
B
1
11a+11b
11b
2
3
4
6a
6b
7
8
9
p.cephaleucos(C7)
p.m.separabilis(C6)
p.m.undescribed(C5)
p.m.undescribed(C4)
p.m.undescribed(C3)
p.m.undescribed(C2)
p.m.undescribed(C1excludingC2)
p.discolor(C/E)
p.pipra(D)
p.pygmaea(F)
p.comata(B1+B2)
p.c.bolivari(A1)
p.c.unica(A1)
p.c.minima(A1)
p.c.occulta(A1)
p.c.coracina(A1)
p.anthracina(A2)

**Supplementary Figure 17 – Mitochondrial ND2 gene tree**
Nodes with ultrafast bootstrap scores of lower than 95 are collapsed. The inferred topology is congruent with the topology presented in the main text derived from ddRAD sequencing data, with one exception: Western Napo haplotypes cluster with southern Amazonian lowland haplotypes (see results and discussion). Otherwise, there are no strongly supported conflicts (see discussion above) with our signal from nuclear genomic DNA. Also see supplementary data files.

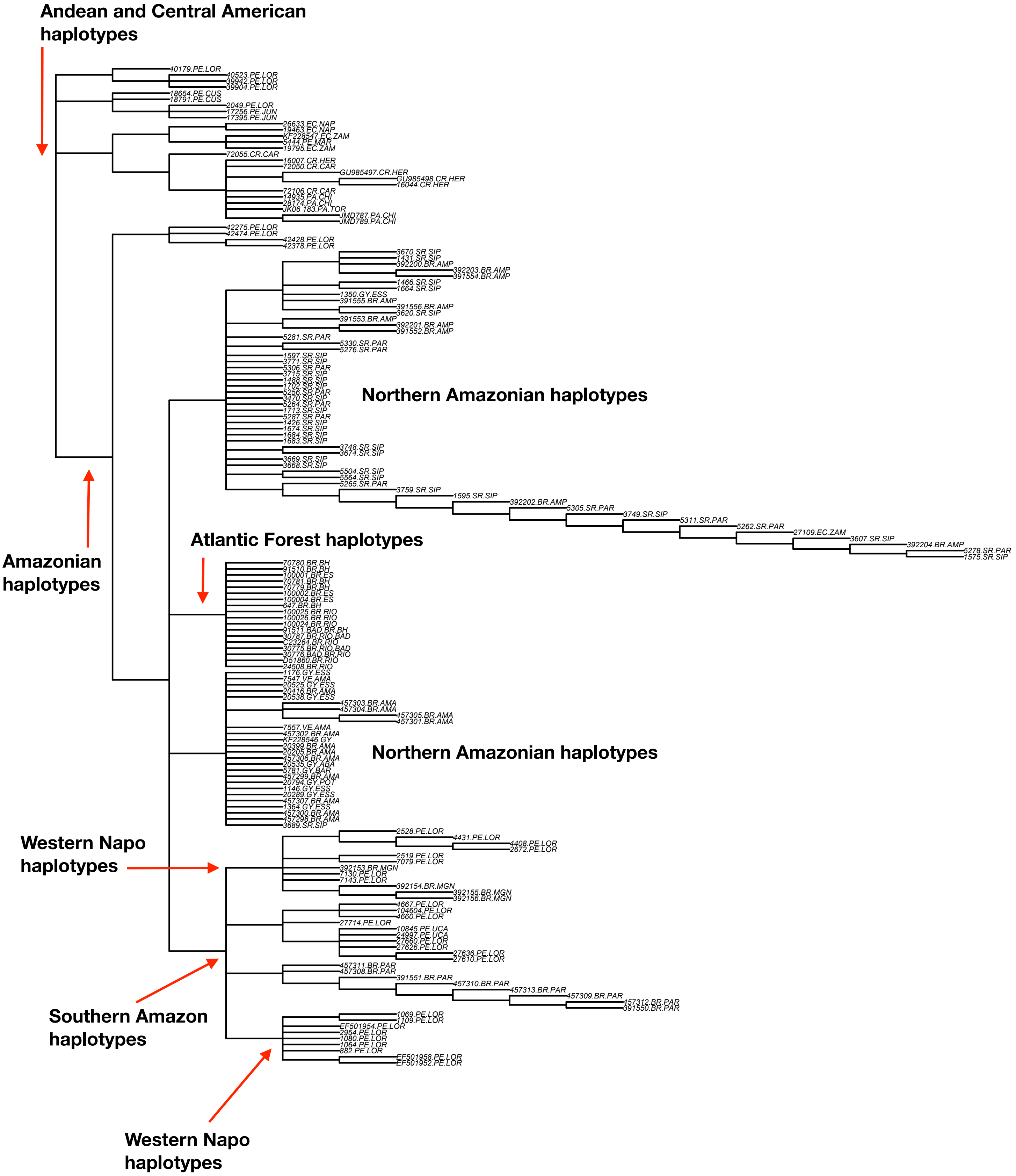

Andean and Central American haplotypes
40179.PE.LOR
40523.PE.LOR
39942.PE.LOR
39904.PE.LOR
18654.PE.CUS
18791.PE.CUS
2049.PE.LOR
17256.PE.JUN
17395.PE.JUN
26633.EC.NAP
19463.EC.NAP
KF228547.EC.ZAM
5444.PE.MAR
19795.EC.ZAM
72055.CR.CAR
16007.CR.HER
72050.CR.CAR
GU985497.CR.HER
GU985498.CR.HER
16044.CR.HER
72106.CR.CAR
14935.PA.CHI
28174.PA.CHI
JK06 183.PA.TOR
JMD787.PA.CHI
JMD789.PA.CHI
42275.PE.LOR
42474.PE.LOR
42428.PE.LOR
42378.PE.LOR
3670.SR.SIP
1431.SR.SIP
392200.BR.AMP
392203.BR.AMP
391554.BR.AMP
1466.SR.SIP
1664.SR.SIP
1350.GY.ESS
391555.BR.AMP
391556.BR.AMP
3620.SR.SIP
391553.BR.AMP
392201.BR.AMP
391552.BR.AMP
5281.SR.PAR
5330.SR.PAR
5276.SR.PAR
1597.SR.SIP
3771.SR.SIP
5306.SR.PAR
3715.SR.SIP
1488.SR.SIP
1702.SR.SIP
5256.SR.PAR
3470.SR.SIP
5264.SR.PAR
1713.SR.SIP
5287.SR.PAR
1426.SR.SIP
1674.SR.SIP
1684.SR.SIP
1683.SR.SIP
3748.SR.SIP
3674.SR.SIP
3669.SR.SIP
3668.SR.SIP
5504.SR.SIP
5564.SR.SIP
5265.SR.PAR
3759.SR.SIP
1595.SR.SIP
392202.BR.AMP
5305.SR.PAR
3749.SR.SIP
5311.SR.PAR
5262.SR.PAR
27109.EC.ZAM
3607.SR.SIP
392204.BR.AMP
5278.SR.PAR
1575.SR.SIP
Northern Amazonian haplotypes
Amazonian haplotypes
Atlantic Forest haplotypes
70780.BR.BH
91510.BR.BH
100001.BR.ES
70781.BR.BH
70779.BR.BH
100002.BR.ES
100004.BR.ES
647.BR.BH
100025.BR.RIO
100026.BR.RIO
100024.BR.RIO
91511.BAD.BR.BH
30787.BR.RIO.BAD
C23264.BR.RIO
30775.BR.RIO.BAD
30776.BAD.BR.RIO
D51860.BR.RIO
24508.BR.RIO
1176.GY.ESS
7547.VE.AMA
20525.GY.ESS
20416.BR.AMA
20538.GY.ESS
457303.BR.AMA
457304.BR.AMA
457305.BR.AMA
457301.BR.AMA
7557.VE.AMA
457302.BR.AMA
KF228546.GY
20399.BR.AMA
20205.BR.AMA
457306.BR.AMA
20535.GY.ABA
5781.GY.BAR
457299.BR.AMA
20794.GY.POT
1146.GY.ESS
20289.GY.ESS
457307.BR.AMA
1364.GY.ESS
457300.BR.AMA
457298.BR.AMA
3689.SR.SIP
Northern Amazonian haplotypes
Western Napo haplotypes
2528.PE.LOR
4431.PE.LOR
4408.PE.LOR
2672.PE.LOR
2519.PE.LOR
7079.PE.LOR
392153.BR.MGN
7130.PE.LOR
7143.PE.LOR
392154.BR.MGN
392155.BR.MGN
392156.BR.MGN
4667.PE.LOR
104604.PE.LOR
4660.PE.LOR
27714.PE.LOR
10845.PE.UCA
24997.PE.UCA
27660.PE.LOR
27626.PE.LOR
27636.PE.LOR
27610.PE.LOR
457311.BR.PAR
457308.BR.PAR
391551.BR.PAR
457310.BR.PAR
457313.BR.PAR
457309.BR.PAR
457312.BR.PAR
391550.BR.PAR
1069.PE.LOR
1109.PE.LOR
EF501954.PE.LOR
2954.PE.LOR
1080.PE.LOR
1064.PE.LOR
882.PE.LOR
EF501958.PE.LOR
EF501952.PE.LOR
Southern Amazon haplotypes
Western Napo haplotypes
